# Recurrent dynamics underlying transient neural representations

**DOI:** 10.1101/2025.06.17.659440

**Authors:** Lars Schutzeichel, Jan Bauer, Peter Bouss, Simon Musall, David Dahmen, Moritz Helias

## Abstract

Brain networks are high-dimensional and interacting complex systems that exhibit substantial structural heterogeneity as well as temporal variability. Yet, when exposed to a stimulus, their recurrent circuits perform reliable computations. The mechanisms underlying this robustness are, however, still mostly unknown. Here, we combine analyses of Neuropixels recordings of awake, behaving mice with models and theory of recurrent neural networks to identify three core computational characteristics that emerge from the interplay of many network constituents and drive dynamic, reliably classifiable stimulus representations. We find that the level of recurrent inhibition in circuits and the microscopic chaos of dynamics respectively drive mean population responses as well as the within– and across-class stimulus response similarities. These core characteristics in turn non-trivially interact to predict and shape the experimentally observed separability of visual and tactile stimulus representations in mouse superior colliculus. Using these characteristics to assess the information transmitted through the network for multiple stimuli reveals a trade-off in coding space: increasing the number of stimuli conveys more information but also reduces their separability due to their larger overlap in the finite-dimensional neuronal space. Our analysis predicts that, only for the experimentally observed small population activity, information keeps increasing with the number of stimuli, revealing another crucial advantage of sparse coding.

## Introduction

Animals perceive external stimuli through an intricate network of neurons. Information about these stimuli must therefore be encoded in the activity of neuronal populations. A puzzling observation, however, is the ubiquitously observed neural variability: even when presenting identical stimuli, the evoked responses share some similarity, but they also show considerable trial-to-trial variability across repeated presentations [1, 2, 3]. Such variability is, moreover, amplified by the microscopically chaotic dynamics of recurrent networks [4, 5], so the miniscule changes between stimuli potentially cause large differences in the internal representations already after short times. Variability, in addition, not only exists in terms of the temporal dynamics, but also in terms of the network structure. Basic dynamical properties, such as stabilization of activity by inhibitory feedback [4], oscillations [6], and low average correlations [7, 8, 9] can be explained even by randomly connected network models. But it is unclear how useful computation can arise in such networks despite their temporal and structural variability that seem to stand at odds with the observed reliability of behavior.

Classical proposals to solve this conundrum consist of population codes that have first been found in motor activity and have been associated with motor output [10, 11, 12, 13, 14]; here groups of cells fire coordinately, such that a weighted average over the population provides a reliable signal. Alternatively, averaging could be performed over time, so that the averaged firing rate of a neuron is the meaningful quantity [15], rather than the sequence of action potentials. The obvious drawback of such a “rate code” is its low temporal resolution that contradicts the fast processing times observed in mammals [16, 17]. Recent years have brought new insights into the meaning of population patterns by investigating neural representations in both artificial networks [18, 19, 20] and in high-level areas of the brain [21, 22, 23]. We here address the question of reliable representations in lower-level brain areas, such as the superior colliculus in mice, from a different point of view. We start with the central set of prerequisites for a neural code: To identify and classify stimuli, neural representations must preserve similarities between two repetitions of the same stimulus, but they must also be distinguishable between different stimulus classes. This is illustrated in Figure 1a, showing the *N*-dimensional activity space spanned by a set of *N* recorded neurons. Such neural representations are not static, but they evolve over time, where recurrence due to the connectivity plays a crucial role [24]. In the current work, we develop a network theory on how such stimulus representations are deformed by the recurrent dynamics and we gauge and validate this theory by experiments using parallel Neuropixels recordings. Our main finding is that, despite randomness in connectivity, stochasticity in time, and chaoticity of the network dynamics, there exists a robust computational backbone that arises from a concentration phenomenon, namely because networks contain a large number of constituents. We find that despite the high dimensionality of the problem, this backbone is faithfully described by only three key characteristics with a simple geometric interpretation: Each representation is characterized as an ellipsoid, where the distance of the ellipsoid’s center from the origin is the firing rate averaged over the neuronal population (characteristics 1: population mean *R*). The typical extent of the ellipsoid reflects the diversity of responses, its typical trial-to-trial variability. We denote the angle corresponding to this width as 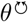 (characteristics 2); the smaller this angle, the more similar are two responses to the same stimulus. When comparing one neural representation with that of another stimulus, the two are separated by a typical angular distance *θ*^↔^ characteristics 3), which reflects the overlap between neural representations of different stimuli, as shown in Figure 1b. A dynamical mean-field theory, valid for large-scale neuronal networks, shows that these three characteristics are robust observables readily accessible in experiments. We expose the fundamental dynamical mechanisms by which recurrence shapes the dynamical interplay of these characteristics. We then show that these characteristics are also sufficient to predict stimulus separability obtained with an optimally trained readout neuron. We consider separability a minimal requirement for meaningful information processing: The signal propagates to downstream areas, where, along the processing hierarchy, representations of different perceptual objects must become increasingly separable to enable high-level cognition.

**Figure 1.**
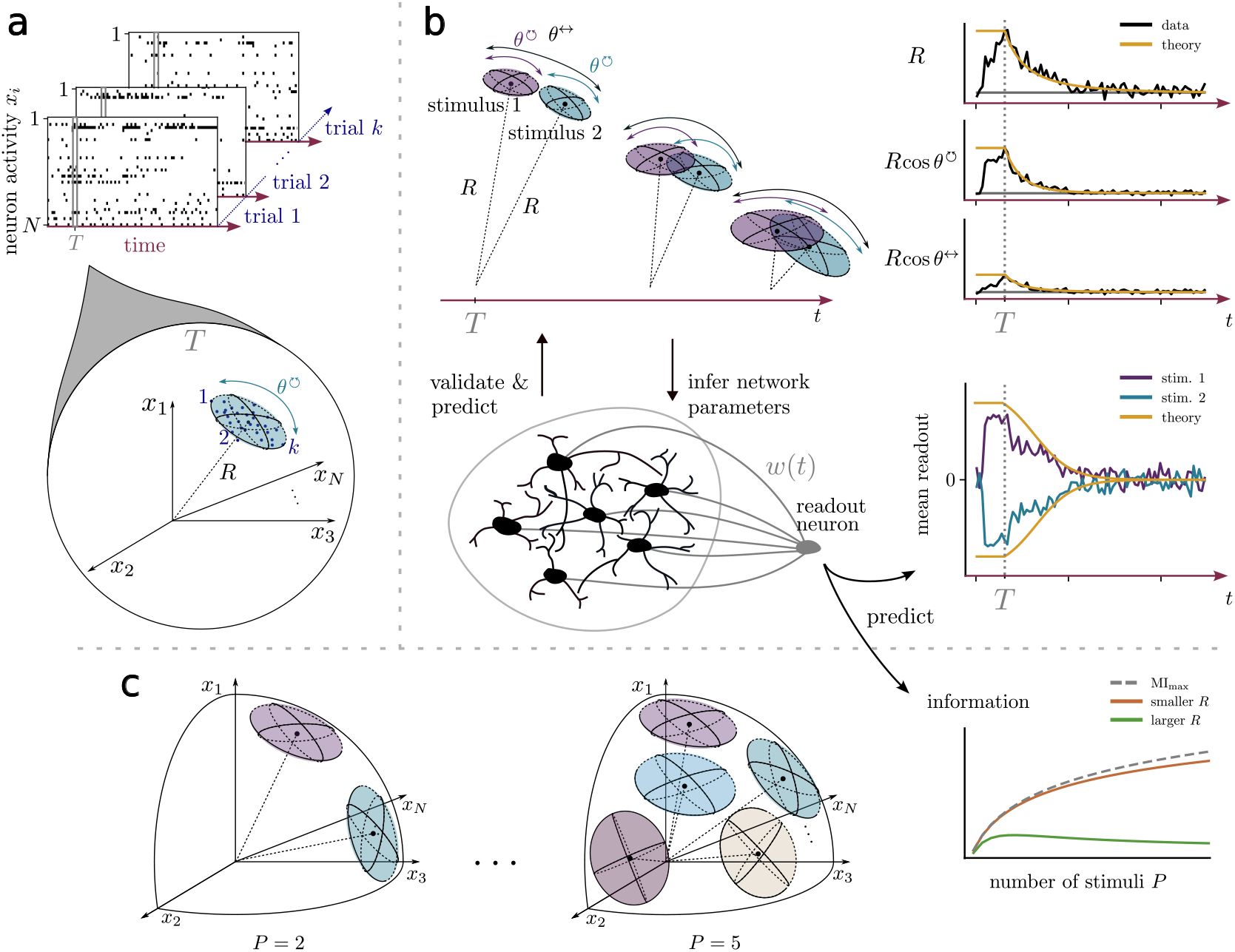
Overview. **a**: Raster plots of parallel neural recordings in superior colliculus from awake, behaving mice in response to stimulation, e.g. a tactile stimulus (air puff to whisker). For any point in time *T* and any trial *k* 1, …, *N*_trials_, the network activity is represented by a neural state vector, depicted as one point in neuronal space. We refer to different types of stimuli (e.g. visual or tactile) as “stimulus classes”. Multiple trials of the same stimulus class form a point cloud in neuron space representing this stimulus class. Point clouds in neuron space are characterized by their distance from the origin *R* and their extent 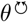. These two quantities yield a representative ellipsoid for each cloud. **b**: Temporal evolution of the representation of stimuli from two classes (with relative angle *θ*^↔^), for example, tactile (blue) and visual (dark purple), after stimulus offset at time *T*, shown for data and and mean-field theory. An optimally trained linear readout ***w***(*t*) quantifies the separability between the classes of stimuli for every point in time, which is determined by the time courses of *R*(*t*), 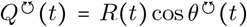, and *Q*^↔^(*t*) = *R*(*t*) cos *θ* ^↔^(*t*). **c**: The separability measure also determines the information content contained in the population signal in the case of *P* ≥ 2 stimuli. Encoding more stimuli generally increases information, while crowding of neuron space opposes this effect. The average population rate *R* controls a sharp transition between expanding (brown, small *R*) and vanishing (green, larger *R*) asymptotic information transfer.

The remainder of the manuscript is organized as follows: In Section 1.1 we analyze stimulus responses in the mouse superior colliculus (SC) obtained by massively parallel single unit Neuropixels recordings in awake behaving mice that were passively exposed to different stimuli. The SC is not only important for the processing of visual information in mice [25, 26], but it is also highly multimodal and therefore suitable for the analysis of different stimulus modalities [27, 28, 29]. Using large-scale recurrent network models and tools from statistical physics of disordered systems [30, 31], we show in Section 1.1 that, in the limit of many neurons, the separability of two neural representations is fully characterized by the three key characteristics, *R*, 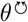, and *θ*^↔^. Furthermore, we make use of the large-neuron limit to derive an effective mean-field description of the network dynamics. We show that the three characteristics form a closed set of equations which entirely governs their dynamics. For the recurrent network model to reproduce the three experimental characteristics, we show that it is sufficient that parameters assume the right statistics; a fine-tuning on the level of individual synaptic weights, for example, is not required. This suggests universal computational properties inherent to such networks. By fitting the model to data, we find parameters that yield a balanced network in which negative feedback stabilizes the dynamics and decorrelates neurons. The intrinsically chaotic network dynamics, in addition, promotes dissimilarities across stimulus responses [4, 8, 32, 33].

In Section 1.2, we study how this dynamic interplay shapes stimulus separability. To assess separability, we use a linear readout neuron that is optimally trained on experimental data using a Bayesian framework. Therefore, we obtain a signal that separates the two stimulus representations (Figure 1b). We show that the three characteristics also determine the separability achieved by the downstream neuron, establishing a direct link between these variables and the separability of neural representations.

In Section 1.3, we highlight the importance of the dynamic interplay that shapes separability: We present two settings of the network model which show, counterintuitively, that the population mean *R* is not the main indicator of information retention – different temporal dynamics of *R* may indeed lead to identical separability. To better understand the mechanism behind this invariance, we derive a time evolution equation for the separability itself. This formulation allows us in Section 1.4 to predict stimulus settings in which the population mean decays while, at the same time, separability transiently increases as a result of recurrent network processing [34, 31]. This setting relies on the chaotic dynamics that are not captured by a rate code but emerge from the discrete signaling of neurons, amplifying initially small differences over time [4, 35] and thereby facilitating classification [31].

The developed theory allows us to predict the classification of multiple stimuli *P* in Section 1.5. For the time of maximal response, we compute the information content of the population signal. To this end, we evaluate the mutual information between stimulus and response, which represents the information transmitted from the stimuli, through the network, to the responses. Equivalently the mutual information reflects the ability to infer the stimulus class by observing the readouts. For optimal information transmission, the stimulus representations must overlap as little as possible within the neuronal space. This raises the question of how to optimally distribute these representations (Figure 1c), which is linked to the capacity of neural networks [36]. In contrast to Gardner, who studied the average overlap, we are interested in the *distribution* of mutual overlaps for a given number of stimuli in a high-dimensional neural space by the help spin-glass methods.

We find two distinct regimes, shown in Figure 1c, for the mutual information by studying the asymptotic information transmission (*P* → ∞). They arise from the trade-off between two antagonistic mechanisms: With more stimuli, more information can be transmitted, but at the same time a larger number of stimuli incurs a higher mutual overlap, thus reducing their separability. We observe that the experimental data lie within a regime that is advantageous for information representation. The sparseness of the code is crucial for the separation between these regimes: not only is it beneficial for information representation [37] but, as we show here, a sufficient degree of sparseness is even necessary for non-vanishing asymptotic information transfer.

## 1 Results

### 1.1 Stimulus responses in mouse superior colliculus

The application of different stimuli to mice causes a transient increase of activity in the superior colliculus, an area which shows responses to both visual and tactile stimuli. The mice are exposed to these stimuli over multiple trials. Two of these trials are shown in Figure 2a, where a visual or tactile stimulus was presented. In this raster plot, the stimuli evoke different characteristic responses. The firing rate or population mean 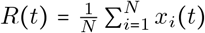, shown in Figure 2b, measures the population activity at a given point in time averaged over both stimuli. Here, *x*_*i*_(*t*) is the binned and binarized activity of neuron *i* in a time bin of 6 ms width at time *t*.

**Figure 2.**
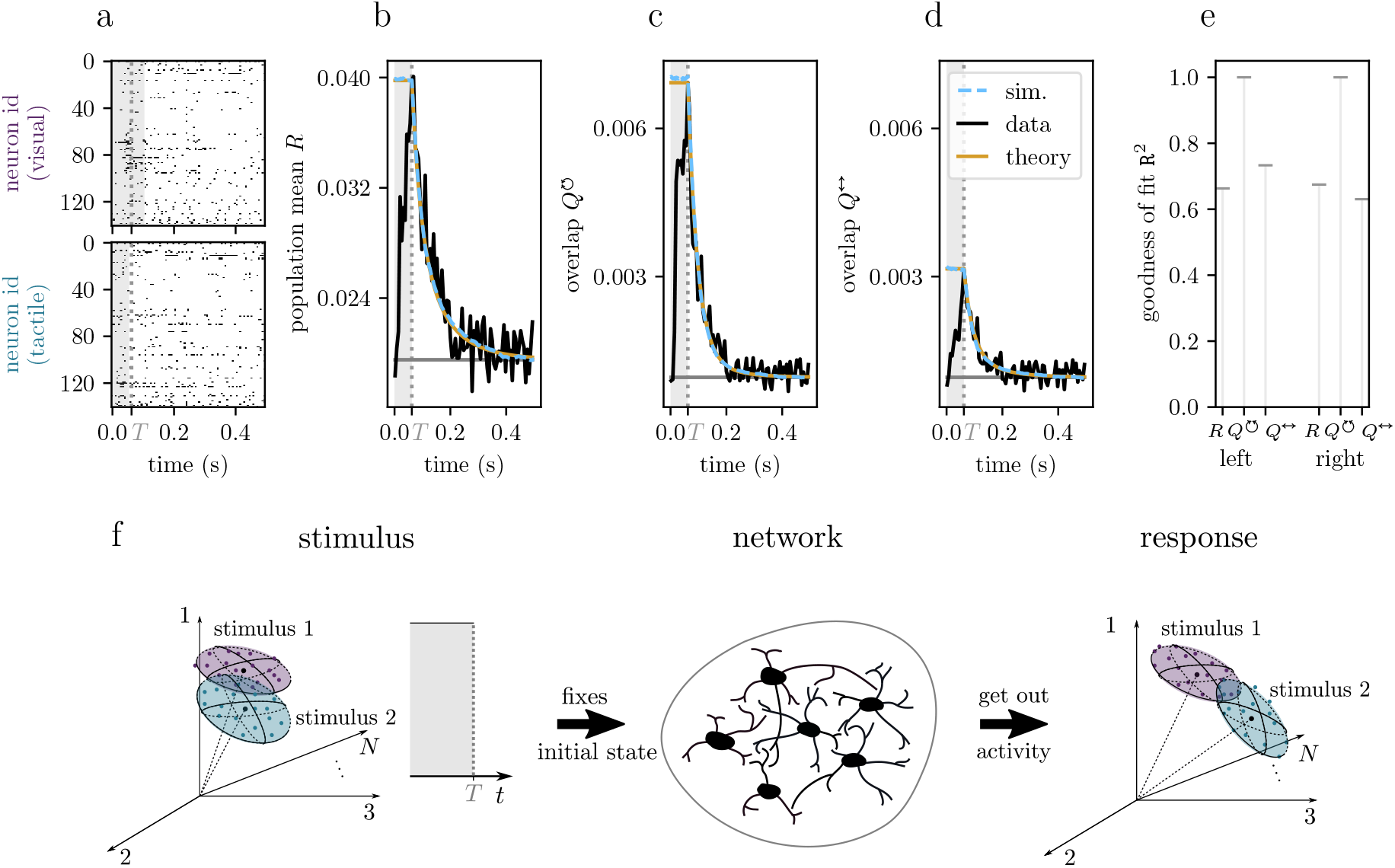
Experimental data and validation of the network model. **a**: Exemplary raster plots for visual (top) and tactile stimulus (bottom) in 6 ms bins. Shaded areas mark the duration of the stimulus, either a noise movie or an air puff, respectively. Neurons are sorted according to depth. **b**,**c**,**d:** Experimental characteristics (black) in counts/bin alongside mean-field theory predictions of the network model (yellow). The overlaps 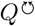 and *Q*^*↔*^ are averages over all pairs of trials of identical or different stimulus classes, respectively. The decay of the experimental characteristics, their baselines, and peak heights are the quantities that are used to determine the statistics of the network model parameters (detailed in Methods 3.3). Numerical simulations of network models (light blue) (for network parameters, see Supplement 3.6). **e:** Goodness of fit R^2^ for the three characteristics from two sessions for two different mice, in which both stimuli (visual and tactile) were on the left or right side, respectively; see Methods 3.3. **f**: Network theory yields a simplified description of the dynamics: A pair of stimuli is represented by two ellipsoids in stimulus space. After fixing the initial state, the stimulus is switched off at time *T* and the recurrent dynamics start to deform these initial representations. The time of stimulus application is indicated by a boxcar ending at *T*. The corresponding gray dotted vertical lines in panels a,b,c,d indicate the stimulus switch-off at *T*.

The population mean increases with stimulus onset and then relaxes back to baseline. Due to the binary nature of the spiking data, the population mean is directly related to the length of the neural state vector *R*(*t*) = ∥ ***x*** (*t*)∥^2^.

The responses exhibit variability across trials. We measure the similarity of responses over time between two trials *α* and *β* by the overlap of two neural states: 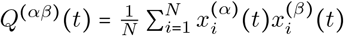. Comparing Figure 2c with Figure 2d, as expected, the variability of responses between different stimulus classes is larger than the variability of responses within the same stimulus classes, corresponding to a smaller overlap *Q*^↔^ in panel d compared to a larger overlap *Q*^↔^ in panel c. These overlaps are directly related to the angles introduced in Figure 1. As time progresses, both overlaps decrease and ultimately approach the same asymptotic value, indicative of chaotic network dynamics which tend to amplify initial differences of states over time.

To quantitatively understand the dynamics of *R*, 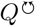, and *Q*^↔^, we build a recurrent network model of *N*_model_ neurons. We here employ a binary recurrent neural network model where neurons have a threshold, an average time *τ* between neuron updates and are connected by a random connectivity matrix with Gaussian distributed weights [31] (see Methods 3.2). The thresholds have an average value δ and a variability *γ*, while the connectivity has mean 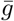 and variance 1/*N*. For large numbers of neurons *N*_model_, we show with mean-field theory (Methods 3.2), that the three quantities *R*,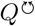, and *Q*^↔^ follow a closed set of equations that governs the network dynamics. Moreover, the dynamics of the three quantities, to leading order in model size, do not depend on the precise realizations of the connectivity and the neurons’ parameters, but only depend on their statistics, thus exposing a stereotypical collective behavior emerging from the recurrent network. The population mean *R* follows a self-consistency equation

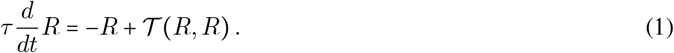

The concrete form of the function *T* and its dependence on the network parameters is given in Methods 3.2. The structure of the equation for *R* is consistent with previous works [4, 38, 39] and, in particular, is independent of the overlaps.

Mean-field theory for both *Q*s requires measuring the similarity of responses to different stimuli between two copies of the same network. This is accomplished using a two-replica approach, where each replicon corresponds to one of two trials, yielding the pair of equations

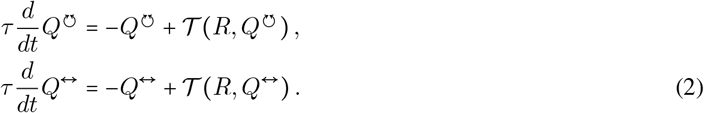

In these three equations lie the core characteristics that govern computation in recurrent networks. The coupling of the differential equation for the population mean (1) and (2) indicates a *dynamic interplay*: The population mean *R* influences the time evolution of both similarity measures 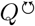 and *Q*^↔^ via *T* (*R, Q*). This function is the same as in the time evolution equation for *R* (1) and can be interpreted as a correlation transmission function, which quantifies the correlation of the output activity of a pair of neurons when driven by a pair of inputs of mean *R* and of mutual correlation *Q*.

The mean-field equations are used to determine the statistics of the network model parameters so that *R*, 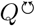, and *Q*^*↔*^ reproduce the experimental characteristics. We determine the statistics of network parameters by first constraining them by the experimental data and by then performing a least-squares fit (see Methods 3.3).

The mean-field theory, the experiment, and the simulations for the similarities are presented in Figure 2b-d. The goodness of fit of the theory to the experiment is shown in Figure 2e, evaluated here using the coefficient of determination `R`^2^ (see Methods 3.3). The fit yields similar statistics of the network parameters for both mice (see Supplement 3.6). In particular, we find that the network connectivity has to be inhibition-dominated 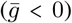 to reproduce the experimentally observed sparse activity.

The inference of the statistics of the network parameters only requires the study of the coupled set of differential equations (1) and (2) for initial conditions on *R* and *Q* given by the experimental data. To bring a pair of network models into such an initial state, one needs to apply a pair of artificial stimuli that drive the response into subspaces of the activity space that correspond to this initial state (Figure 2f). In the limit of large networks, only the Gaussian correlations of these artificial stimuli matter. They can thus be represented by ellipsoids in stimulus space. When the networks have settled into their statistically stationary states at time *T*, the network reproduces the initial activity in the experimental data by the three characteristics, which corresponds to the ellipsoid abstraction of the point cloud. From this initial time point on, the artificial stimuli are switched off and the ellipsoids resembling the initial representation are continuously deformed by the recurrent dynamics in line with the experimental observations.

### 1.2 Intrinsic dynamics shape separability of stimulus representations

The three characteristics *R*, 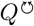, and *Q*^*↔*^ provide an intuitive understanding of how different stimuli are represented. Subsequently, we aim to quantitatively assess the separability of these representations to characterize their computational properties. In experimental recordings, we have access to a limited number of neurons. After measuring the characteristics, determining the parameters of the model, and validating the model, we can use the network model to assess separability for a larger number of neurons than the recorded subset. This will allow us to make predictions for the separability of stimuli in the superior colliculus.

To decode stimulus classes from neuronal activity, we use a linear readout, which may be interpreted as a readout neuron, given that neurons perform a weighted summation of their inputs. At each time point *t*, the readout is *y*_α_ (*t*) = ***w***(*t*)^T^ ***x***_α_ (*t*) + *ϵ*(*t*), with the time-dependent readout vector ***w***(*t*) ∈ ℝ^*N*^ and Gaussian noise *ϵ* (*t*), accounting for stochasticity, for example due to afferent signals to the readout neuron from other brain areas (see Methods 3.4). Using Bayesian inference, we determine the optimal readout vector for each time point *t* to assess what the downstream neuron could optimally capture.

We split the data set into training 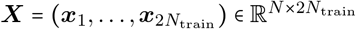 and test data. The binary classification task is mapped to a regression problem by assigning either of two outputs *y*_*α*_ ∈ {−1,1} depending on the stimulus class. These labels are gathered into a vector 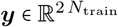.

In the limit of many neurons, the expected label of a test point follows a Gaussian process posterior with mean 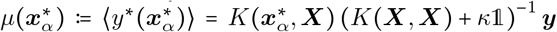, where *K*(○,○) is the kernel matrix of the process [40, 41]. The entries of this kernel are pairwise similarities of network responses, 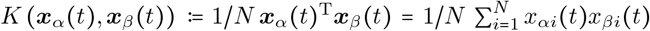.Importantly, in this limit the kernel concentrates around its mean value and can hence be expressed in terms of the key characteristics introduced earlier, i.e. 1/*N* ***x***_*α*_ (*t*)^T^ ***x***_*α*_ (*t*) →*R*(*t*), 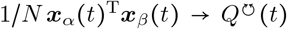 for trials *α* and *β* belonging to the same class, and 1/*N* ***x***_*α*_(*t*)^T^***x***_*β*_(*t*) → *Q*^↔^ (*t*) for trials *α* and *β* belonging to different classes. This limit corresponds to the abstraction step from a point cloud in Figure 3a_1_ to the ellipsoid picture in panel a_2_. Both the experimental and the mean-field kernel show a block-like structure in Figure 3b_1_,b_2_. The mean-field kernel has entries *R* on the diagonal, 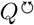 for entries corresponding to the same class (diagonal blocks), and *Q*^↔^ for entries corresponding to different stimulus classes (off-diagonal blocks).

**Figure 3.**
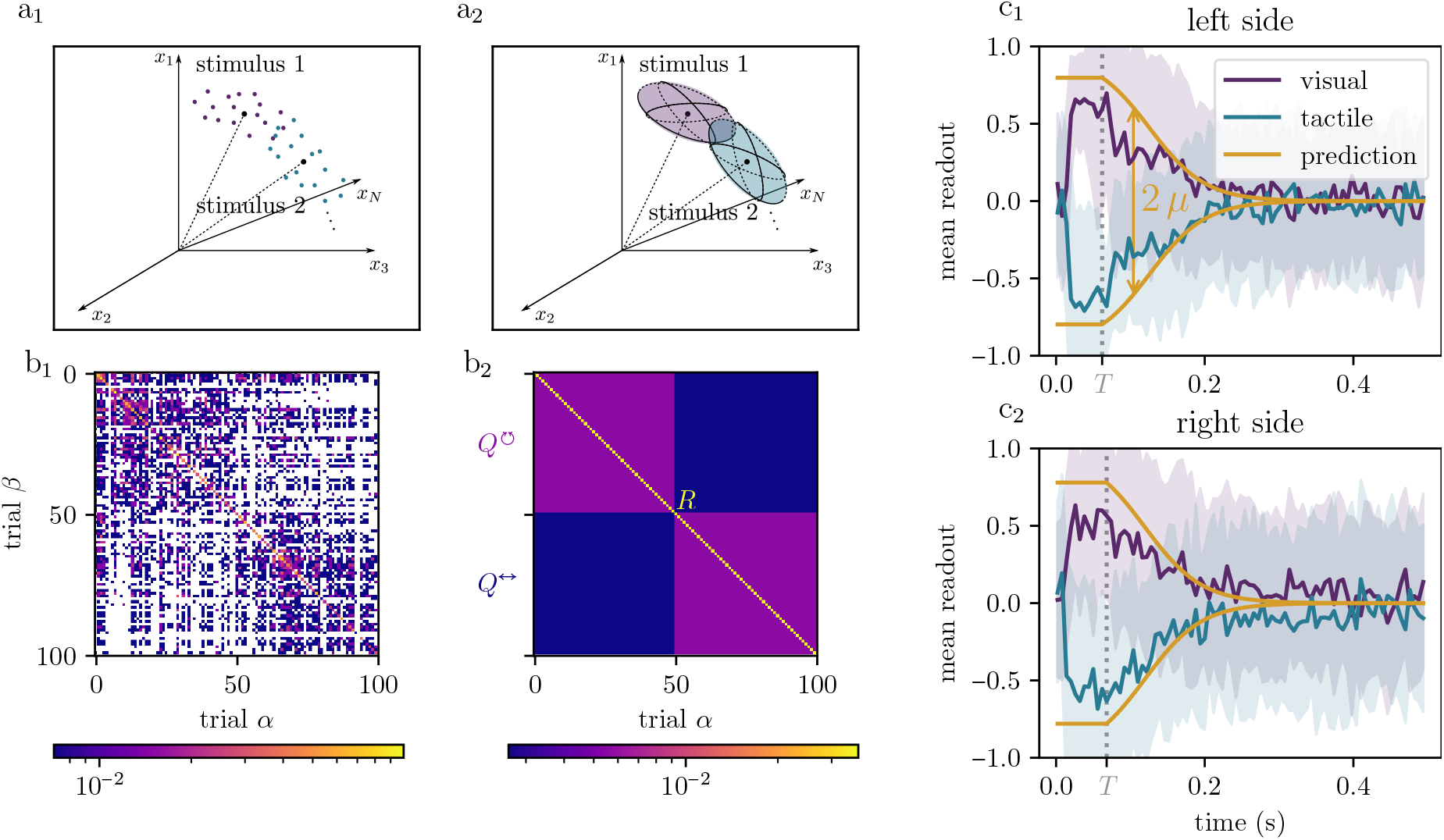
Separability of neural representations: Experiment and mean-field theory. Separability of stimuli is shaped by three characteristics. **a**_1_: Illustration of neural representations for the experimental recordings with a finite number of neurons. **a**_2_: Illustration of many-neuron limit of the mean-field theory, abstracting the experiment. **b**_1_: Experimental kernel matrix corresponding to a_1_ (at *t* = *T*). White spots indicate vanishing entries. The trials are ordered such that the first half belongs to the first stimulus class and the second half to the other class, revealing a block-like structure. **b**_2_: Kernel matrix for the mean-field theory. In the many-neuron limit, the kernel matrix between visual (with label *y*_*α*_ = 1) and tactile stimulus (with label *y*_*α*_ = −1) in mouse superior colliculus (SC), only has three distinct values: *R*, 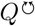, and *Q*^↔^, indicated by different colors. **c**_1_, **c**_2_: Time course of separability obtained with the kernels from b_1_,b_2_. **c**_1_: Visual and tactile stimulus on the left side. **c**_2_: Visual and tactile stimulus on the right side. The shaded area represents the standard deviation of predicted labels over test data points and seeds from sampling the test data. For parameters of network model and simulation parameters, see Supplement 3.6.

The scalar separability 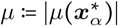 can be derived analytically

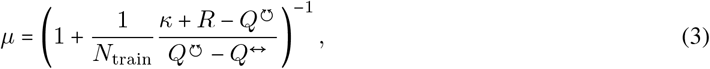

where *κ* is the variance of the readout noise (derived in Methods 3.4).

Therefore, we can understand separability assessed by a downstream neuron by the interplay of the three characteristics *R*, 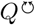, and *Q*^↔^. Specifically, the separability depends on a ratio of differences: It decreases with the difference between the population mean and the within-class overlap (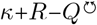), and increases with the difference between the within-class and the between-class overlap (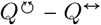). The former is a measure for the variability within one class compared to the maximal overlap *R*, while the latter can be understood as the signal, such that the ratio of differences in equation (3) resembles a ratio of the total variability to the signal. Notably, separability vanishes if stimuli within one class are as similar as stimuli between the two different classes, 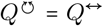. Furthermore, separability increases with the amount of training data per class, *N*_train_, and decreases with the readout noise *κ*. Adding noise has the same effect as increasing the population mean, while keeping the overlaps the same.

The experimentally observed separability and the extrapolation towards larger numbers of neurons than those recorded, obtained by equation (3), are shown in Figure 3c_1_ and c_2_. As expected, the extrapolated separability from mean-field theory exceeds the one obtained from the experimentally subsampled neurons; this is because the kernel matrix in the many-neuron limit is homogeneous within each block (Figure 3b_2_), while the experimental one has some heterogeneity (Figure 3b_1_). Separability is highest immediately after stimulus application, and the time courses of the experimental and the predicted separability match well.

### 1.3 Firing rate decay alone does not reflect stimulus information retention

Having obtained an expression for the separability *µ*, we now ask how it is affected by the quantities *R*, 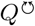, and *Q*^↔^. Traditionally, a persistent firing rate or population mean is *R* considered indicative of information retention. In the model, the time course of *R* is controlled by inhibitory balancing, whereas 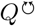 and *Q*^↔^ are driven by chaotic dynamics. Specifically, the rate of decay of *R* can be adjusted by modifying the ratio between inhibition and threshold (see Methods 3.2). All other parameters are obtained from the network adapted to the dataset of mouse 1 (stimuli on left side). In this particular case, despite the differences in the decay of *R* (Figure 4b_1_, b_2_), the decay of the overlaps in Figure 4c_1_, c_2_ changes in such a way that the separability in Figure 4d_1_, d_2_ evolves similarly. The three important characteristics *R*, 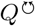, and *Q*^↔^ enter the mean-field equations (2) and the equation for the predictive mean (3) in a dynamic fashion, but remarkably, the alterations to *R* cancel each other out in this particular case and yield a similar decay of the separability. Why does such a cancellation happen?

**Figure 4.**
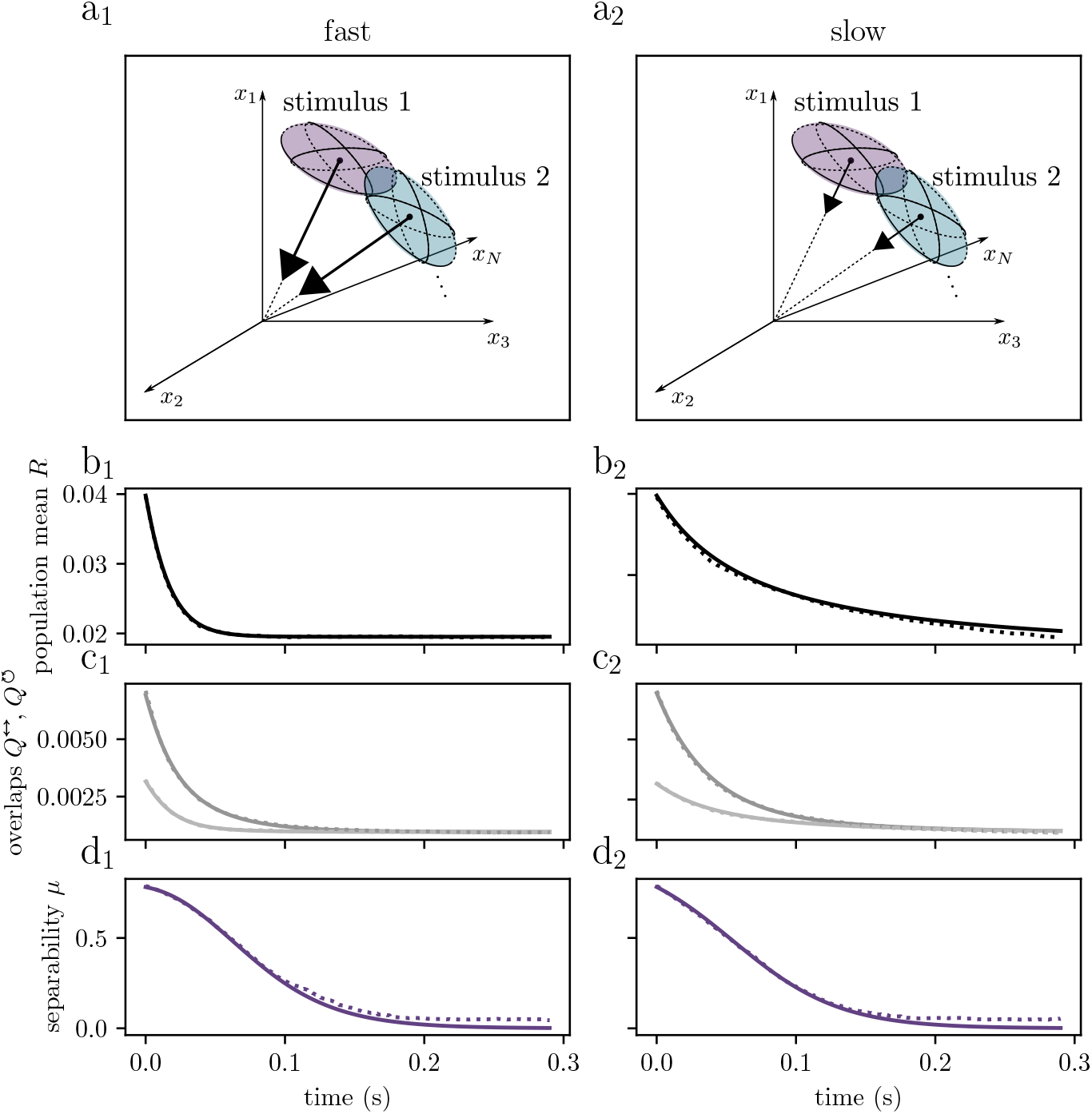
Two scenarios of fast and slow decay of population mean with similar decay of separability. **a**_1_, **a**_2_: Initial representations in neural state space. We compare two scenarios with different parameters that affect the decay of the population mean. The arrows indicate the different velocities of the population mean decay. **b**_1_, **b**_2_: Decay of the population mean that alters the decay of the within-class overlaps (dark gray) and between-class overlaps (light gray) in **c**_1_ and **c**_2_. Solid curves are theoretical predictions, dotted curves are simulations. **d**_1_, **d**_2_: Decay of the predictive mean for the different velocities of the decays of the population mean. The network parameters are those from the network model (see “left side” in Supplement 3.6), except for the ratio between inhibition 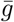 and threshold δ, respectively. The decay of the population mean is modulated by changing the slope of *T* (*R, R*) (Methods 3.2). Resulting parameters in Supplement 3.6.

We gain insight into the decay of the separability by deriving its time evolution analytically (additional details in Section 3.4)

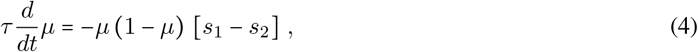

which shows that the separability evolves slowly whenever *µ ≈* 0 or *µ ≈* 1. The right hand side is, however, not expressible in terms of *µ* alone, which means that the time evolution of the separability is not completely determined by the separability at an initial time – rather, the time course of the three characteristics *R*, 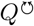, and *Q*^↔^ explicitly influence the separability via the two terms *s*_1_, *s*_2_.

The first term, *s*_1_, measures the relative change of the total variability 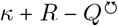 while the second term *s*_2_ measures the relative change of the signal 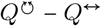. The difference between these two terms, by equation (4), modulates the speed by which *µ* evolves. Although the total variability decays faster with a rapidly decreasing population mean, the separability can exhibit a similar decay in both cases if the signal decay adjusts accordingly. The time evolution of the separability may also be understood geometrically, see Supplement 4.1.1.

This analysis highlights that firing rate alone is insufficient to infer information retention. Instead, our model suggests that the dynamic interplay of the three characteristics *R*, 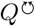, and *Q*^↔^ is crucial. This interplay can lead to a similar decay in separability even when the firing rates decay on different time scales. In particular, even a quickly decaying firing rate may be accompanied by correlations *Q* that are informative about the stimulus for a considerably longer time. Thus, stimuli can still be separable after the transient in the population mean has significantly subsided.

### 1.4 Transient increase in separability for low response variability and hard tasks

The analytical form of the time evolution of separability (equation (4)) allows us to explore how the decay changes for different values of *Q*^↔^ and 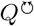.

In the experimental data, tactile and visual stimuli can be clearly distinguished as shown by the separability in Figure 3c_1_,c_2_. To increase the difficulty of distinguishing stimuli, we decrease the distance between stimulus classes in the model, thus increasing their overlap *Q*^↔^: the centers of the ellipsoids in Figure 5a are closer together. Simultaneously, we increase the overlap within classes to ensure that it remains larger than the overlap between classes. This corresponds to the ellipsoids becoming narrower in Figure 5a compared to Figure 4. The ratio between the distance of the ellipsoids and their extent 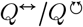 reflects the task difficulty.

**Figure 5.**
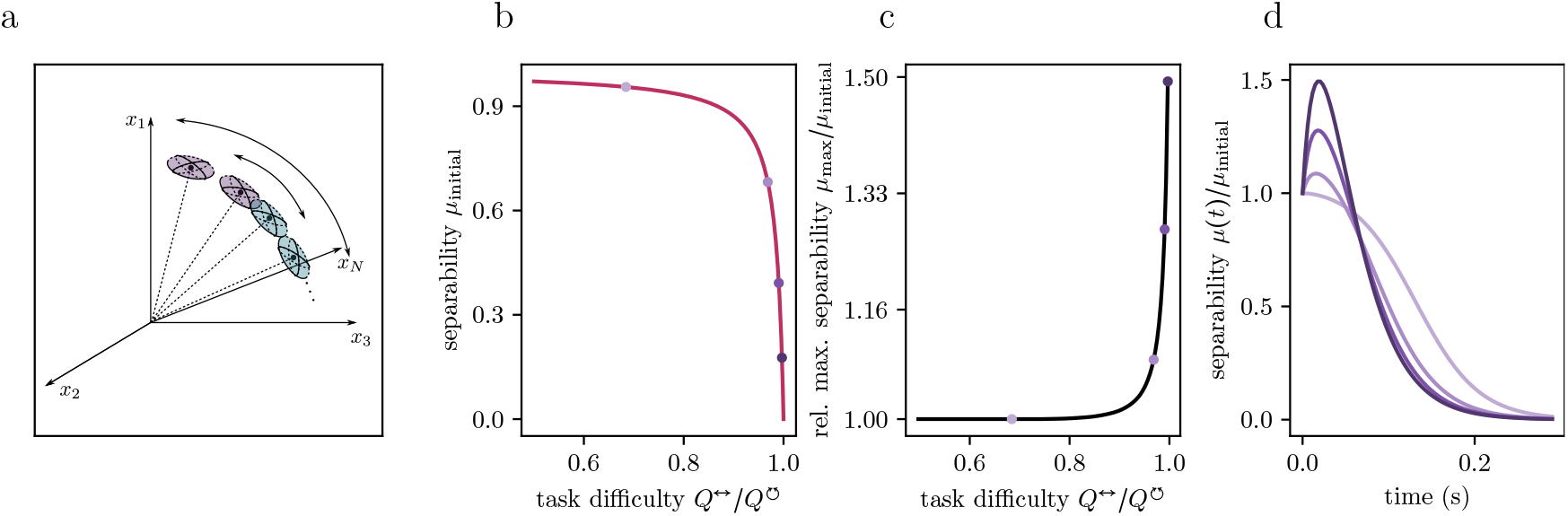
Separability transiently increases for stimuli that are initially hard to separate. **a**: Illustration of the initial state that is modified compared to the experiment. The overlap within one stimulus class 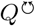 is increased, reflected in narrower ellipsoids compared to Figure 4. The hardness of the task 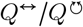 is varied as indicated by the black arrows. **b**: Initial value of separability *µ*_initial_ decreases with task difficulty. **c**: The maximum value of separability over time, *µ*_max_ as a function of the task difficulty in relation to the initial value *µ*_initial_. **d**: Separability over time normalized by the initial value. Different shadings correspond to different task difficulties, as indicated by markers in b and c. Parameter values: 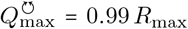, ratio of similarities: 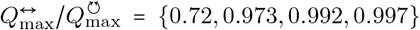, variance of noise *κ* = 0.5*R*_max_, amount of training data per class *N*_train_ = 35.

In the regime of larger task difficulty, we observe a transient increase in separability over time (Figure 5d). The reason is that chaos drives smaller overlaps (or larger distances) faster apart than larger overlaps (or smaller distances), particularly in biologically-inspired neuronal networks that have threshold-like activation functions [31]. This is due to the correlation transmission function *T* (*R, Q*) becoming infinitely steep as *Q* approaches *R*, inducing a transient increase in separability.

According to equation (4), a transient expansion occurs when the signal grows faster than the total variability, i.e., when the second term *s*_2_ is larger than the first term *s*_1_ in equation (4). So for a transient increase the signal has to increase (*s*_2_ > 1) and a sufficiently large portion of the total variability must be accounted for by the readout noise *κ* (see Methods 3.4 for more details).

While separability decreases with task difficulty in Figure 5b, the expansion is more pronounced for harder tasks (Figure 5c,d), where the ellipsoids are initially closer together. The transient increase in separability occurs at lower response variability than seen in the current set of experimental data. We expect that more similar pairs of stimuli will lead to representations that are harder to distinguish and may therefore enter the regime of transient increase.

### 1.5 Information in population signal

Having elucidated the mechanisms by which recurrent dynamics shape stimulus representations, we next investigate how much information about the stimulus class is encoded in the neural activity. As the brain must routinely distinguish among a far greater number of sensory inputs, we leverage the model to extrapolate beyond the case of binary classification studied so far. The aim is to examine how information encoding scales with the number of stimulus classes. This allows us to assess the model’s capacity for stimulus discrimination in more realistic settings.

In addition, we systematically vary the parameters of the model to explore whether different regimes emerge with respect to information transmission. To obtain the maximal information capacity across time, we choose the time point of peak population response, providing a snapshot of maximal representational separation. We then analyze the asymptotic behavior as the number of stimulus classes increases, which serves as a proxy for the model’s information capacity. Finally, by locating the parameters in the phase space of the model, we characterize the regime in which the superior colliculus (SC) operates.

We consider *P* stimuli, each eliciting a characteristic network state, see Figure 6a. Rather than focusing on the distance between two stimulus representations, we must now consider the joint embedding of *P* stimuli, which inevitably generates overlap between pairs of representations. This overlap *Q*^↔^ (*P*) in the limit of large numbers of neurons, can be derived by help of spin-glass methods (more details in Methods 3.5) as

**Figure 6.**
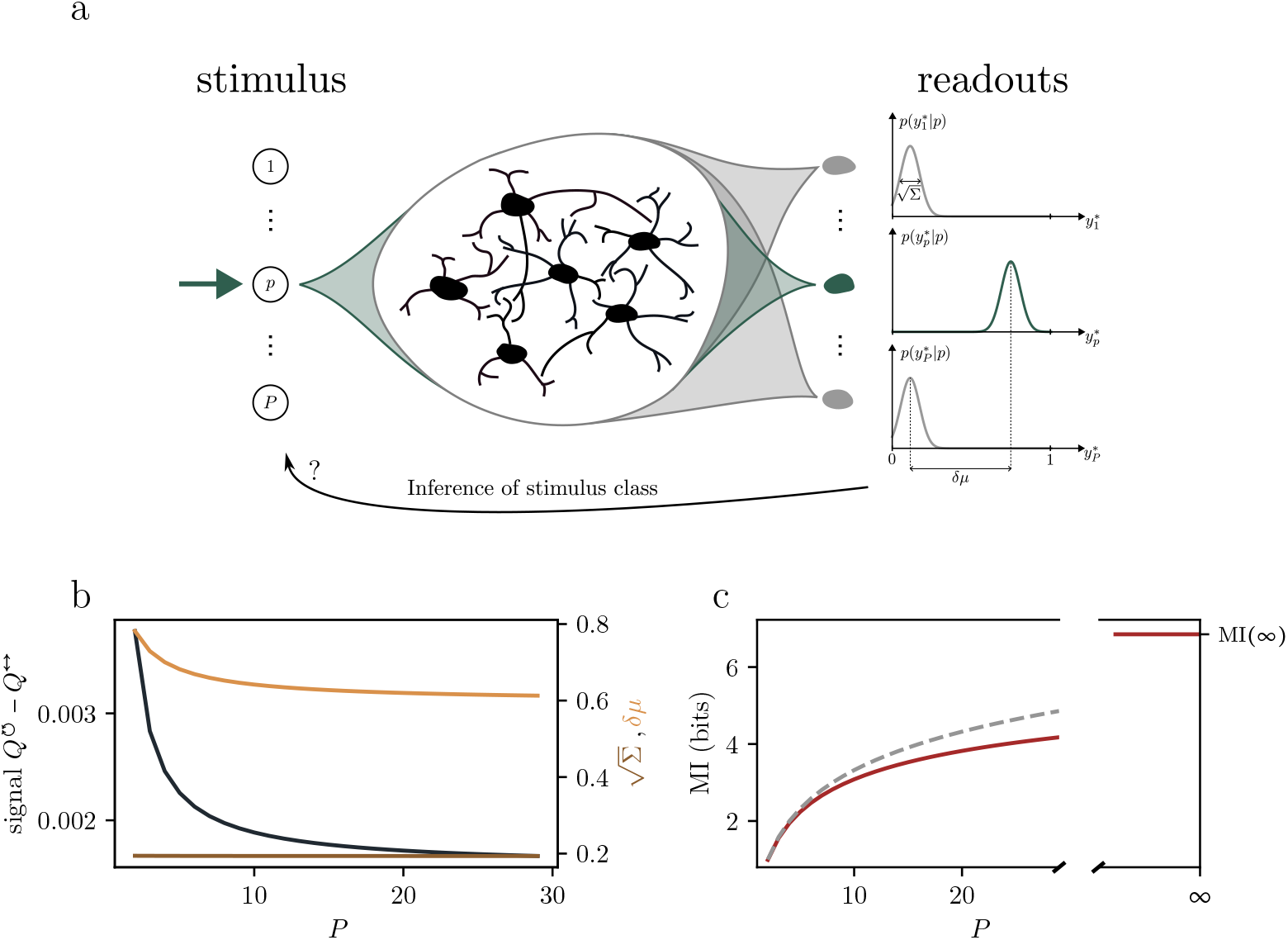
Separability for more than two stimuli: Information transmitted from stimuli through the network to readouts. **a**: Schematic drawing: The network is subject to one of *P* stimuli. The elicited network state is read out by *P* linear readout neurons, one for each stimulus: The *p*-th linear readout is trained such that its response is 1 for the *p*-th stimulus and zero otherwise (one-hot encoding). Through the overlap with the other stimuli, the other linear readouts also show a response for the *p*-th stimulus. **b**: The effect of crowding in neuron space. Left axis: The signal (the difference between within– and between-class overlap) decreases with increasing *P* (black). Right axis: Separability *δµ* (orange) decreases accordingly and posterior variance (brown) approximately stays constant as a function of *P*. **c**: Left: Mutual information 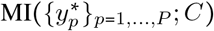; *C* between readouts 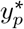 and stimulus classes *C* as a function of number of stimuli *P* for the experimental characteristics (red curve). The dashed curve indicates the maximally possible mutual information log_2_ *P*. The asymptotic information transmission (the saturation value of the mutual information for *P→* ∞) is indicated in the broken axis panel. Parameter values are as in Supplement 3.6.

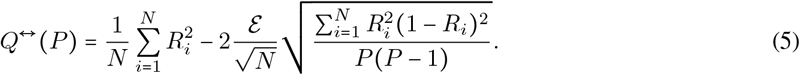

The elicited network states in response to stimuli are binary vectors in neuron space, where each neuron *i* has a firing rate *R*_*i*_, measured from trial averages in the experiment. The first term 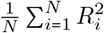 in equation (5) is the typical overlap of representations if neurons were firing independently across stimuli. An optimal system will use the space more efficiently, which is captured by the second term reducing the overlap *Q*^↔^ (*P*). The degree to which the brain is close to this optimum is controlled by the efficiency 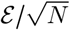, which is determined using the experimentally measured overlap of two stimuli 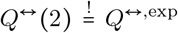. From equation (5), we observe that as *P* increases, distributing stimuli becomes more difficult because of their finite extent: The overlap increases and asymptotically approaches the average overlap given by the first term. This leads to a reduction in the signal, as shown in Figure 6b. Consequently, the separability in the limit of many neurons also decreases with *P*, as seen in Figure 6b. In the limit of many neurons, the separability is again given by equation (3), denoted as *δµ* (*P*) here, with *Q*^↔^ = *Q*^↔^(*P*) depending on *P* (for more details see Supplement 5).

#### Balancing information and overlap

How much information is contained in the population signal for *P* stimuli? In general, more stimuli provide more information; however, the neuronal space becomes increasingly crowded, thereby increasing *Q*^↔^ (*P*), which makes it more difficult to infer the stimulus class from the responses in Figure 6a. To quantify the trade-off between the information gain and the decreasing separability, we use the Kullback-Leibler divergence *D*_KL_ (○‖○) to assess the mutual information between stimuli and responses

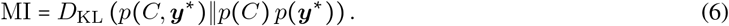

It captures the information that is transmitted from the stimuli through the network to the responses. We consider the case of all *P* stimuli being equally likely 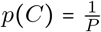. The responses are given by the readouts. Each stimulus evokes a distinct network state, which is subsequently decoded by the set of *P* linear readout neurons. Here, the *p*-th readout neuron is trained so that its response is largest for the *p*-th stimulus and is supposed to be zero else. However, non-matching readout neurons also show non-vanishing responses because of the mutual overlap between different stimulus representations. In this context, separability refers to the mean difference between the target readout’s output and that of all others, see Figure 6a. We denote the variability of the outputs with Σ. In comparison to the separability, this variability Σ (*P*) ≈ Σ approximately stays constant, as shown in Figure 6c.

The ratio between these two quantities

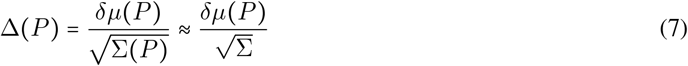

is the effective separability, which, in addition to *P*, fully determines the mutual information in equation (6) as

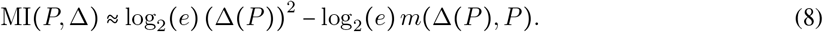

Here, *m*(Δ (*P*), *P*)is a parameter resulting from an approximation of a sum of random variables (see Methods 3.5 for more details).

Using the experimentally measured values, we find that the mutual information increases as the number of stimuli grows, following a trend similar to its theoretical maximum, log_2_ *P*, as shown in Figure 6c.

For an infinite number of stimuli, the mutual information, however, saturates

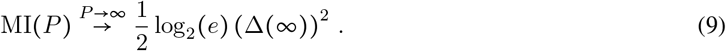

This limit is formally derived in Supplement 5.3, and it admits an intuitive interpretation: Each new readout provides an additional projection of the network state. However, as the number of readouts increases, the incremental benefit of an additional projection diminishes, as most of the network has already been effectively accessed.

The effect that limits the growth of the mutual information is the crowding of stimulus representations in neural space. The saturation value of the mutual information arises from the balance between the crowding and the information gained from more stimuli in the population signal. How does this balance depend on the population mean *R*? To operationalize this question, we scale each individual neurons’ rate *R*_*i*_ by the same factor, which preserves the shape of the experimentally measured firing rate distribution {*R*_*i*_}_*i*_. This scaling argument is based on the typically observed exponential dependence of the firing rate on the mean input to the neuron [42] (for more details, see Supplement 5.5). From this consideration it follows how the overlap between classes changes with the population mean in equation (5).

In Figure 7a, we observe two factors that enhance mutual information: (1) low response variability within classes and (2) a small population mean. A cross-section of mutual information along the line preserving the experimentally measured neuron class-selectivity is shown in Figure 7b. Every neurons’ class-selectivity is quantified by the correlation coefficient *ρ*_*i*_, see Supplement 5.5. For a very small population mean, the mutual information decreases due to the finite noise *κ* on the readout, as seen in Figure 7b. Two phases emerge in Figure 7a: In the expanding regime (for small population mean), the saturation value of mutual information is positive, and mutual information increases with *P* in a similar fashion to its theoretical maximum, log_2_ *P*. In the vanishing regime (for higher population mean), the mutual information initially follows the log_2_ *P* trend, reaches a peak, but then declines when *P* increases further, eventually approaching zero. Examples for the two cases in comparison to the case for the experimentally observed population mean are shown in Figure 7c. The transition between the regimes is continuous, with the point of vanishing mutual information corresponding to where the overlap within classes equals the overlap between classes 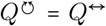. For approximately constant neuron class selectivity, the overlap between classes *Q*^*↔*^ grows more rapidly with the firing rate than the overlap within classes 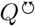, leading to such a transition point. This can be seen from Figure 7a, where the phase boundary between the expanding and the vanishing regime corresponds to 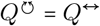. As the population mean *R* increases, the line indicating this boundary has a steeper slope than the black line indicating the overlap within classes 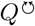. The transition point between the two phases – where these two lines intersect (near the green point) – remains the same when scaling class selectivities *ρ* → *ρ*^small^, *ρ*^large^ (dashed gray lines).

**Figure 7.**
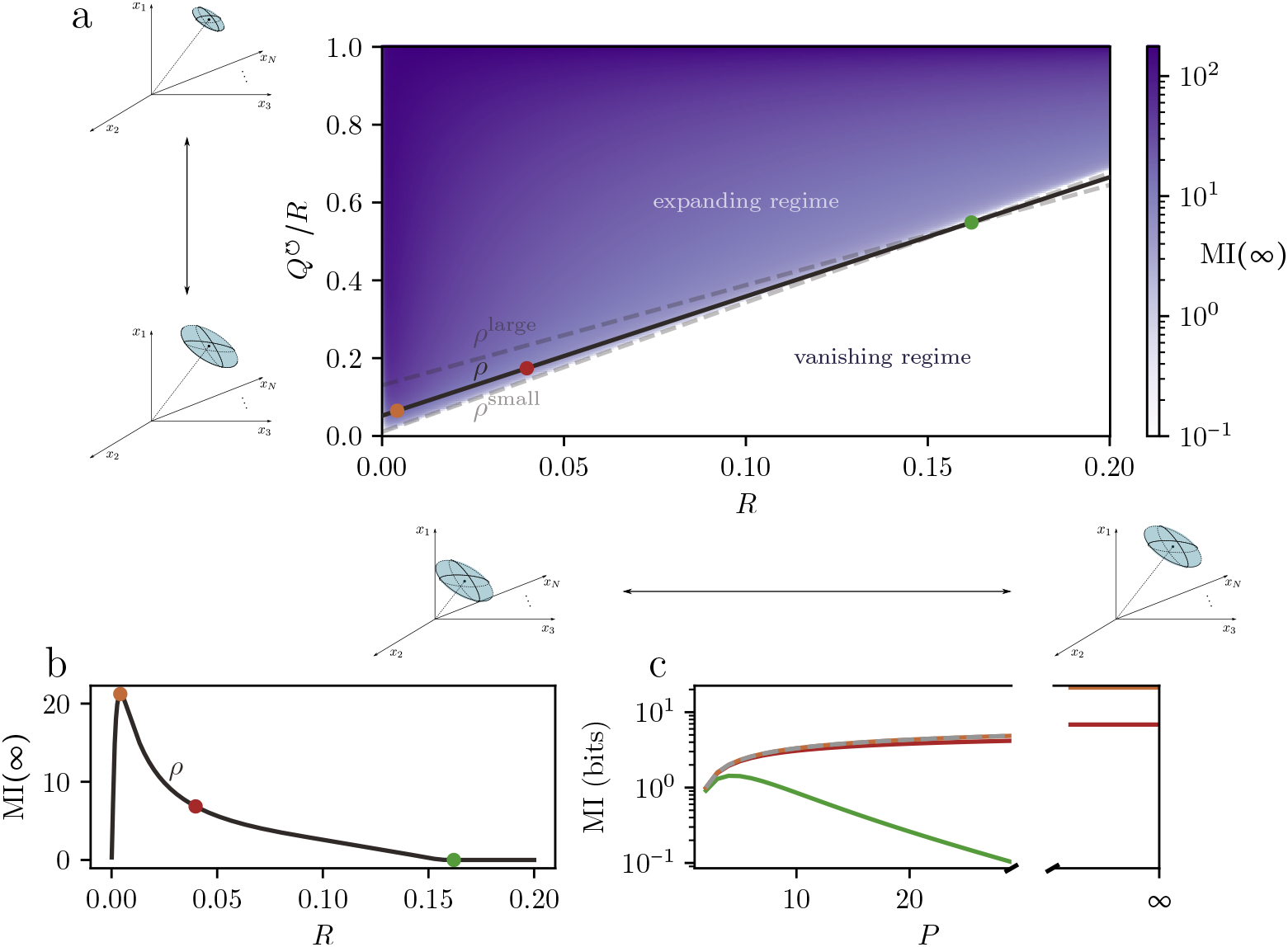
Regimes for asymptotic information transmission. **a**: Asymptotic value MI(∞) for different population means *R* and overlap within classes 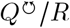. 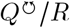 can be understood as the (cosine of the) angle 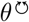 that measures the extent (variability) of the neural representation ellipsoid for one stimulus. The colored area represents the regime where MI (∞) ≠ 0. White space represents the vanishing regime, MI(∞) = 0. The vanishing regime (white) is separated from the expanding regime (purple) by a line indicating the phase boundary. Along this line,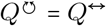. Values smaller than 0.1 bits are colored white. The black curve in panel a represents values of 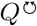 as a function of *R* for constant Pearson correlation coefficient *ρ*_*i*_ for every neuron *i* (the gray dashed line indicates smaller (*ρ*^small^) or larger (*ρ*^large^) Pearson correlation coefficients than in the experiment; see Supplement 5.5). Dots indicate different population means. The intermediate population mean (red) is identical to the experiment with *R*^exp^ = 0.04 (same as in Figure 6c). The small population mean (brown dot) indicates optimal sparseness. The large population mean (green dot) is an example of vanishing asymptotic information transfer in the vanishing regime. **b**: Asymptotic mutual information MI(∞) along the black line of panel a. **c**: Asymptotic mutual information MI(∞) as a function of *P*. Colors correspond to the different population means that are indicated as dots (green, brown) in panel b. The curve for the optimal asymptotic (brown) value starts very close to its theoretical maximum MI_max_ = log_2_ *P* (dashed, gray). The mutual information curve decreases and ultimately vanishes if the crowding effect is too strong (green). The saturation value of the MI is indicated by the broken axis. Parameter values are as in Supplement 3.6.

In conclusion, a lower population mean *R* expands the effective encoding space, leading to better information representation. Sparse coding, therefore, is not only advantageous for information representation but is also a necessary condition for non-vanishing asymptotic information transfer.

## 2 Discussion

In this study, we discover a central computational mechanism at work in recurrent brain networks. We arrive at this insight by a combination of high density Neuropixels recordings, models, and theory. The neural data show that activity organizes in neural trajectories [11, 43, 44, 22, 45, 46] that first align upon stimulus presentation and subsequently decorrelate. We demonstrate by theory and in the data that in large networks only three core characteristics generally govern the evolution of this dynamics and thus of neural representations: the *population firing rate* (*R*) and two types of *correlations* 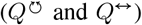 between pairs of neural trajectories. These characteristics provide summary statistics of neural trajectories across stimuli and trials, that are invariant under the ubiquitous variability and heterogeneity of brain networks, thus capturing a robust and universal mode of information processing. We demonstrate this further by showing that a downstream neuron’s ability to separate two neural representations is determined by the same three key characteristics alone.

Our analysis concerns the superior colliculus, a midbrain structure that integrates high-dimensional input from the retina, the primary visual cortex, and primary somatosensory cortex in addition to having recurrent connections. It distills this multisensory information and initiates motor commands [26]. As neural representations of different stimuli are transmitted from the superior colliculus to higher-level areas, they need to become separable. In this work, we demonstrate that this separability in recurrent networks is in general shaped by the dynamic interplay of the three core characteristics *R*, 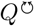, and *Q*^↔^: We extend the two-replica mean-field theory from [31] by incorporating transient firing rate dynamics and temporally sparse activity, and thereby reveal the critical role of these three characteristics in the characterization of neural representations. Moreover, this theory provides analytically tractable time-evolution equations that describe the dynamical interaction of these characteristics in recurrent neural networks in a self-consistent manner. Specifically, it uncovers how inhibitory balancing controls the time course of the population mean *R*, while chaotic dynamics drive the overlaps 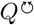 and *Q*^↔^. Together, these three characteristics govern stimulus separation in a downstream readout neuron, uncovering the mechanisms that underlie separability.

We test this theory by applying it to experimental data. The theory shows that network parameters tuning inhibition-excitation balance, thresholds, and the neuronal time constant enter only at the level of their statistics. To explain the time courses observed in experimental data, it is thus sufficient to fit these statistics – there is no fine-tuning needed on the level of individual neurons or synapses. This suggests that the network model contains the crucial building blocks for network-level processing. Furthermore, the agreement between the theoretical predictions of separability dynamics and experimental results supports the sufficiency of this theoretical framework.

With this network model, we can explore further implications of the theory. We highlight that information retention crucially depends on the mutual interplay between the three characteristics rather than on the firing rate alone: We demonstrate two cases where separability decays similarly, despite differences in firing rate decay. Previous studies have shown that a transient change in firing rate causes long lasting trial-to-trial decorrelation in disordered balanced networks [15, 35]. Our findings suggest that information is contained within the degree of decorrelation. We predict a transient increase in separability for stimulus representations that are hard to separate and exhibit low within-class variability. This transient increase occurs because after stimulus offset, the signal grows faster than the total variability, proposing a hypothesis on how chaotic dynamics on the network level contribute to computation, even if the firing rate only decreases.

By considering multiple stimuli, we identify two regimes for information transfer in the limit of large numbers of stimuli, that are separated by a sharp phase boundary: the expanding regime at low firing rate and a vanishing regime at higher rates. The experimental data with its sparse activity and corresponding low rate lies within the expanding regime. We conclude that sparse coding is not only beneficial, but also necessary for information representation and information transmission.

Our network model has only few parameters, which, surprisingly, are sufficient to reproduce and explain the main features of the data. In this sense, the theory is minimal. Specifically, we assume random Gaussian connectivity, whose sufficiency lies in the fact that the resulting model tracks the key characteristics relevant for a downstream neuron reading out from a large network. In particular this connectivity gives rise to the overlap *Q*, which captures the variability homogeneously across neurons, leading to the depiction of the neural activity over multiple trials as ellipsoids in Figure 1a. This formulation is advantageous because it remains agnostic to the specific direction of variability.

In reality, the superior colliculus is a midbrain structure composed of different laminae, which in turn are composed of different types of neurons that differ in their responses [26]. For neurons reading out from a small subset of neurons (specific parts of a brain area), it may become important to incorporate structure in the connectivity. Accounting for structured connectivity [45, 47, 48] would skew the ellipsoids, altering the directions of variability they measure and thereby affecting their subsequent decorrelation. Measuring the variability along a specific axis may therefore be advantageous when neural activity is confined to a manifold with a specific directionality [19, 49], and may become important for more complex task setups and recordings in higher-level areas. In our formalism, including structure in the connectivity can be achieved by relaxing the assumption of statistical homogeneity across all neurons, thereby introducing heterogeneity.

The prediction of separability in the mean-field approach corresponds to an extrapolation to large networks, allowing a readout neuron to achieve a higher signal-to-noise ratio than that directly estimated from the observed experimental data. This is consistent with previous findings showing that finite-size corrections reduce separability [50]. The expression for separability is derived in the limit of large network size [40, 41], while keeping the amount of data finite. In machine learning, this corresponds to the lazy-learning regime [51], where training the network is equivalent to training only the readout [52, 53]. In other words, in this limit, training the whole network does not substantially change the recurrent weights relative to their initial values. Consequently, the simple and analytically tractable expression for separability remains valid.

For networks that learn features, one could adopt a Bayesian approach similar to that used to study artificial networks, which is agnostic to the learning rule [54, 55], or implement plasticity into the field-theoretic model [56, 57]. This would enable modeling of learning and task adaptation, with a focus on how the interaction between neuronal and plasticity timescales influences neural representations and their separability.

In the experimental data, a transient increase of separability after stimulus offset has not been observed. However, we predict a transient increase in separability after stimulus offset for stimuli that are more difficult to separate (i.e., higher task difficulty) and exhibit lower variability within stimulus classes. Such a transient increase would correspondingly enhance the amount of information conveyed in the population signal, with a maximum occurring later than immediately after stimulus application.

In addition to stimulus-related sources, variability can also stem from ongoing neural activity, presumably reflecting varying brain states [1] or corresponding to unobserved behavior [3, 58, 59]. It would be interesting to measure how trial-to-trial variability changes with learning [60, 3], particularly between and within highly similar stimuli, and to test the theory under conditions of lower response variability.

To quantify mutual information, we consider a set of *P* stimuli. Rather than considering the full network state, we project it onto a number of readout neurons equal to the number of stimuli, which then defines the response. This presents a dimensionality reduction, where every readout neuron is optimal in a Bayesian sense to maximize the mutual information between inputs and outputs [61, 62, 63, 64]. We again leverage the fact that a downstream neuron separates network states according to the geometry in neural activity space given by the three characteristics *R*,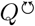, and *Q*^↔^. This allows us to relate mutual information directly to separability.

The amount of mutual information depends on the choice of the prior variance for the readout weights (*g*_*w*_), which sets the scale for the noise in the readout neurons. We choose this variance such that the correlation between readout neurons matches that of the experimentally recorded neurons.

The asymptotic amount of information increases with decreasing firing rate, supporting the sparse coding strategy of the brain [37]. This occurs because the total variability, 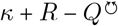, decreases faster with decreasing population mean than the signal,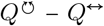, leading to an enhanced signal-to-noise ratio that determines the separability *δµ*.

Sparse coding enables efficient storage [36, 65], communication, and computation despite drastic compression [66]. In addition to these computational advantages, it also reduces energetic costs: another benefit of sparse coding [67]. Our findings do not rely on these previous arguments and thus offer a novel perspective on the classical advantages provided by sparse coding.

For sufficiently high rates (low levels of sparseness) we observe a regime in which the asymptotic information transfer vanishes and a maximum amount of information appears at a certain number of stimuli. This optimal number of separable stimuli arises from a trade-off between transmitting more information per stimulus and the neuron space becoming too crowded, which increases the mutual overlap of responses to unrelated stimuli. The vanishing regime arises for a larger population mean than that observed in the experiment. To extrapolate to such higher population means, we scaled the firing rate of each neuron by a common factor, which is valid for small population means, where the neuronal activation function behaves approximately like an exponential [42]. As the activity increases, the activation function becomes more linear. Reassuringly, however, the vanishing regime remains prevalent and is robust to these changes, as discussed in Supplement 5.5.

It would be interesting to experimentally test whether neural representations of stimuli in a finite neuronal space are organized in a way that optimally transmits information according to the mutual information considered here. Concretely, the behavioral accuracy of mice as a function of task complexity could be investigated to test whether the data falls into the expanding or vanishing regime.

## 3 Methods

### 3.1 Experimental design and statistical analysis

We analyze parallel neuronal in vivo electrophysiology recordings from mouse superior colliculus. All experiments were carried out in accordance with the German animal protection law and local ethics committee (LANUV, NRW)

#### Animals

Animals were 16-week old male mice. Surgical details for electrophysiological recordings are described in [68].

#### Recordings

To record from superior colliculus (SC), mice were head-fixed on a rubber coated wheel and Neuropixels probes [69, 70] were inserted into the left hemisphere of the brain. The bottom 384 channels of the Neuropixels probe were then used to record a high pass filtered action potential signal (30 kHz). We simultaneously recorded *N* = 141(mouse 1) and *N* = 65 (mouse 2) neurons in the SC.

#### Stimulation

Visual stimulation was performed using Gaussian noise movies displayed at 60 fps for 100 ms at a 17 cm distance from the right eye on a gamma-corrected LED-backlit LCD monitor (Viewsonic VX3276-2K-MHD-2, 32”, 60 Hz refresh rate). Monitor brightness was calibrated to 60 lux. Tactile stimulation consisted of puffs of air that were given in short bursts with a duration of 50 ms and a pressure of 0.03 bar. Spouts delivering the stimulation were targeted to hit the whiskers of the mouse and aimed so that they did only elicit whisker deflections but no additional startle reactions. Stimuli were delivered for 50 (mouse 1) and 40 (mouse 2) trials each. The time between individual trials was 3-5 s.

#### Pre-processing

The Neuropixel recordings provide voltage traces for each channel, from which spike waveforms stemming from different neurons have to be identified and classified. For spike sorting, Kilosort 2 was utilized for the data set of mouse 1 and Kilosort 2.5 for mouse 2 [71]. We bin the data in 6 ms bins, leaving us with a discretized time axis and an entry *x*_*αi*_ (*t*) ∈ ℕ_0_ for every neuron *i*, trial *α*, and time *t*. For this bin size only 1.2% (mouse 1) and 1.0% (mouse 2) of all non-zero entries are greater than 1. In the following we set these entries to 1 to ensure Boolean values *x*_*αi*_ (*t*) ∈ {0, 1}. Additional details can be found in Supplement 1.

### 3.2 Recurrent neural network model

We use a network of binary neurons, where the neurons can be in either of two states: actively firing *x*_*i*_ (*t*)=1 or quiescent *x*_*i*_ (*t*) = 0 [72, 73, 4, 74, 75, 76]. The state of each neuron *i* is dynamically updated according to the synaptic input 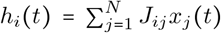, which is the weighted sum of the neurons’ activities *x*_*j*_ (*t*) with the connectivity matrix *J*. If a neuron’s input *h*_*i*_ exceeds its threshold, it transitions to the active state; otherwise, it will be in the quiescent state *x*_*i*_ (*t*) = 0. Neuronal activation is thus described by the Heaviside function *H*(○). This constitutes a well established simple model for brain cortical activity [77, 4, 8]. For propagating the state of the neurons, we use Glauber dynamics [78], selecting neurons for update according to a Poisson process with exponential interval distribution of mean *τ*.

#### Network parameters

The network model is minimal in the sense that it is described by only four network parameters: The process of updating neurons is governed by a Poisson process with time constant *τ*. The connectivity matrix is drawn randomly 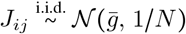, where the average strength of neuronal couplings is 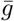, the sign of which determines whether the network is excitation– or inhibition-dominated. Every neuron has an individual threshold, which the synaptic input has to surpass for the neuron to fire. Thresholds are distributed with mean *δ* and variability *γ* across neurons. The variance of the connectivity is fixed to 1 *N* without loss of generality; a different numerator would merely rescale the fields and could be absorbed by the other parameters. This is due to the Heaviside activation function having no inherent scale *H* (*ax*)= *H* (*x*) ∀ *a* >0 [76].

#### Dynamical mean-field theory

A recurrent network model with binary units has been shown to be analytically tractable using dynamical mean-field theory [4, 31, 30]. We here extend this theory by a time-varying mean activity to account for stimulus-driven network transients. To study how a network responds to two different stimuli, we consider two instances of the network (replicas) that share their connectivity, resembling two trials from the experiment. We obtain self consistent equations for the fields *R*(*t*), 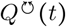 and *Q*^↔^(*t*). For more details, see Supplement 2.

### 3.3 Determining network parameters from experimental data

The right-hand sides of the time evolution equations (1),(2) only depend on certain constellations of observables and parameters. By inferring the experimental values of these constellations we obtain constrains on the parameters. Under these contraints we fit the decay by minimizing a weighted mean-squared error. More details on the fitting procedure in Supplement 3.

#### Goodness of fit

For the experimental characteristics (expm) and the model fit (model_fit) of *R*, 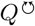, *Q*^↔^, we perform a smoothing-window approach with a window size of 3 bins (= 18ms). Using the sliding window, we calculate the average 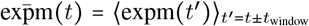 and variance 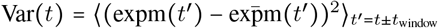 over this window for the experimental characteristics. The goodness of fit is then evaluated using the coefficient of determination

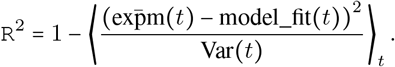

This leaves us with three coefficients of determination, evaluating the goodness of fit for each of the three characteristics separately (see Figure 2e).

### 3.4 Separability

We quantify separability using a linear readout model 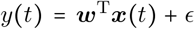 with 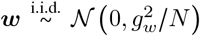 and 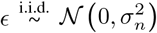. After conditioning on training data 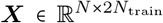 with labels 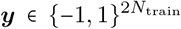in a Bayesian manner, we can obtain the predictive statistics for the test data 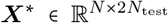. The posterior distribution 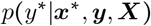 follows a Gaussian process [40]. The two cumulants of this Gaussian process can be calculated using the kernel: The kernel matrix directly follows as the dot-product between two neural state vectors 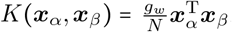. In the limit of large numbers of neurons the entries of this kernel concentrate around the three distinct values 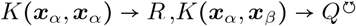 for *C*(*α*) = *C* (*β*) and *K* (***x***_*α*_, ***x***_*β*_) → *Q*^↔^ for *C*(*α*) ≠ *C* (*β*). Here, *C*(○) denotes the class membership of the sample. When sorting the trials by class-membership the block structure becomes evident. The predictive mean in equation (3) depends on the three characteristics *R*, 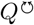 and *Q*^↔^ in addition to the noise 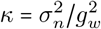 and the number of training points *N*_train_. (more details in Supplement 4)

#### Time evolution of separability

For the time evolution of the separability, the total vairability 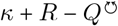and the signal 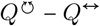 entering the formula (3) are of particular importance. The relative change of the total variability and the signal are governed by the terms *s*_1_ and *s*_2_, respectively. This can be seen by rewriting equations (1) and (2) as

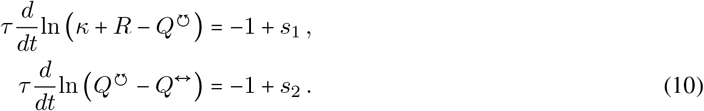

For more details, see Supplement 4.1.1.

### 3.5 Information transmission

We take *P* linear readouts 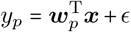 with one weight vector ***w***_*p*_ for every stimulus class *p* = 1, …, *P*. The *p*-th readout is obtained through Bayesian inference to produce the highest output when applying the *p*-th stimulus. Prior weights are as before chosen according to 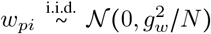. Because prior and likelihood factorize over *p* [79, Chapter 11, (11.2)], the posterior covariance is also diagonal between readouts *p*. The Bayesian posterior quantifies the conditional probability distribution of responses *p*(*y*^*^|*C*), where 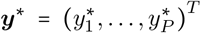, which is necessary for evaluating the mutual information. Together with the probability distribution across stimuli *p* (*C*) = 1 /*P*, the mutual information is fully determined. Since the mutual information between the stimuli and responses depends only on the ratio 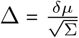 it suffices to consider the difference of the mean readouts. The latter is given by equation (3). The overlap within classes 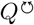 is kept constant across the different numbers of stimuli, i.e. the extent of each stimulus is kept the same. This assumption is based on experimental observations. More information in Supplement 5.

#### Optimal overlap between classes

We here calculate the optimal overlap *Q*^*↔*^ between different stimulus representations in equation (5) for the case that stimuli are distributed efficiently in the available neuronal space. The assumptions entering the derivation of this overlap are informed by both the data and network model. We observe that stimuli evoke similarly high responses with similar population firing rates and similar extents. Therefore, we keep *R* and 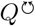 the same for all stimulus classes. Network responses are captured well by the mean-field theory, in which cross-correlations between neurons vanish. Therefore, we assume that neurons *i* fire independently from each other as 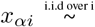 Bernoulli (*R*_*i*_), but coordinated across trials. More details in Supplement 5.4.

## Code and data availability

The datasets generated and analyzed during the current study are available from the corresponding author on reasonable request.

## Acknowledgments

We thank Tobias Kühn, Inés Samengo, Severin Graff and Thomas Rüland for helpful discussions. This work is funded by the Deutsche Forschungsgemeinschaft (DFG, German Research Foundation) as part of the SPP 2205 – 533396241 and the Gatsby Charitable Foundation. Open access publication funded by the Deutsche Forschungs-gemeinschaft (DFG, German Research Foundation) − 491111487. The authors gratefully acknowledge the computing time granted by the JARA Vergabegremium and provided on the JARA Partition part of the supercomputer JURECA at Forschungszentrum Jülich (computation grant JINB33).

