## Supplementary Material for "Recurrent dynamics underlying transient neural representations"

October 1, 2025

### 1 Experimental data

Neuropixel electrodes are an electrophysiological tool for recording spiking data. They reliably record individual neurons over long timescales with $384$ channels and can locate the depth by recording one spiking neuron with multiple channels [1].

In both sessions, mice have electrodes inserted into the left hemisphere of the brain into primary visual cortex (V1) going down into the superior colliculus (SC). We thereby obtain simultaneous recordings of $N = 141$ (mouse 1) and $N = 65$ (mouse 2) neurons in the superior colliculus. The mice are fixated on a rubber coated wheel that allows for leg movement. While mouse 1 has two screens for visual stimulation (noise movie) in addition to one air puff directed at each whisker (ipsi- and contralateral stimulation), mouse 2 only has one screen and one air puff on its right side (contralateral stimulation). Both mice receive passive stimulation with a chosen stimulus modality, which is applied twice in every trial.

The Neuropixel recordings provide one voltage trace for each channel, from which spike waveforms that stem from different neurons have to be identified and classified. For spike sorting, Kilosort 2 was utilized for the data set of mouse 1 and Kilosort 2.5 for mouse 2 [1, 2].

We bin the data in 6ms bins, leaving us with a discretized time axis and an entry $x_{\alpha i}(t) \in \mathbb{N}_0$ for every neuron $i$, trial $\alpha$, and time $t$. For this bin size, only $1.2\%$ / $1.0\%$ of all non-zero entries are greater than $1$. In the following we set these entries to $1$ to ensure Boolean values $x_{\alpha i}(t) \in \{0, 1\}$.

### 2 Binary networks as a model for spiking data

#### 2.1 Field theory for random networks

In this section we derive the mean-field equations for $R$, $Q^{\circlearrowleft}$, and $Q^{\leftrightarrow}$. The derivation uses the field-theoretic framework of [3]. In our model, the state of a neuron takes on Boolean values $\{0, 1\}$ resembling spiking data instead of Ising values $\{-1, 1\}$ used in [3]. This changes the formulas in comparison to [3].

The interplay of neurons and their mutual influence is modeled by considering an input field $\boldsymbol{h} \in \mathbb{R}^N$ influencing the neuronal state $\boldsymbol{x} \in \mathbb{R}^N$ via a (potentially stochastic) input-to-output relation of neurons that can in general be described by a probability functional $\rho[\boldsymbol{x}|\boldsymbol{h}]$. The joint probability density of neuronal inputs and states is given by

$$\rho[\boldsymbol{x}, \boldsymbol{h}] = \rho[\boldsymbol{x}|\boldsymbol{h}]\, \rho[\boldsymbol{h}], \quad \rho[\boldsymbol{x}|\boldsymbol{h}] := \prod_{i=1}^{N} \rho[x_i|h_i], \tag{1}$$

where the latter factorization arises from the conditional independence of the neurons when conditioned on their input $h_i$. By providing the relation from the field back to the neuronal state $\rho[\boldsymbol{h}(t)|\boldsymbol{x}(t)] = \delta\left[\boldsymbol{h}(t) - J\boldsymbol{x}(t)\right]$ which is mediated by the connectivity matrix $\boldsymbol{J}$, these equations can be closed

$$\begin{aligned}\rho[\boldsymbol{h}] &= \int \mathcal{D}\boldsymbol{x}\;\rho[\boldsymbol{x}|\boldsymbol{h}]\delta[\boldsymbol{h} - \boldsymbol{J}\boldsymbol{x}] \\ &= \int \mathcal{D}\boldsymbol{x} \int \mathcal{D}\hat{\boldsymbol{h}} \exp\bigl(\hat{\boldsymbol{h}}^{\mathrm{T}}\boldsymbol{h}\bigr) \exp\bigl(-\hat{\boldsymbol{h}}^{\mathrm{T}}\boldsymbol{J}\,\boldsymbol{x}\bigr)\rho[\boldsymbol{x}|\boldsymbol{h}] \end{aligned} \tag{2}$$

The scalar products can be understood as contracting the unbound indices from either the neuron or time dimensions

$$\hat{\boldsymbol{h}}^{\mathrm{T}}\,\boldsymbol{J}\,\boldsymbol{x} = \sum_{i,j=1}^{N} \hat{h}_i^{\mathrm{T}}(t) J_{ij} x_j(t) \quad \text{and} \quad \hat{h}_i^{\mathrm{T}} x_j = \int dt \hat{h}_i(t) x_j(t).$$

We assume Gaussian connectivity

$$J_{ij} \overset{\text{i.i.d.}}{\sim} \mathcal{N}(\bar{g}/N,\, 1/N). \tag{3}$$

For simplicity, we do not scale the variance of the connectivity as we will later assume discretely communicating neurons with an activation function that has no inherent scale. For one instance of the network, the connectivity matrix is drawn once and the couplings are kept constant. In the following, we are interested in the statistics over an ensemble of networks with different realizations of the connectivity matrix $J_{ij}$,

$$\langle \rho[\boldsymbol{h}] \rangle_J = \int \mathcal{D}\boldsymbol{x}\;\rho[\boldsymbol{x}|\boldsymbol{h}]\langle \delta[\boldsymbol{h} - J\boldsymbol{x}]\rangle_J.$$

The reason for this is that in the limit of many neurons, the number of synaptic inputs becomes large and thus we expect the distribution $\langle \rho[\boldsymbol{h}]\rangle_J$ over $J_{ij}$ to become strongly peaked around its mean value. In other words, we expect that for the field of one neuron $i$ the variability between different realizations of $J_{ij}$ becomes small, which subsequently implies that the variability between neuron $i$ and neuron $k$ for one fixed realization of $J_{ij}$ becomes small. This property is referred to as self-averaging and conveniently lets us reduce the neuronal dynamics to an equation for only one representative neuron. The appearing term in equation (2)

$$\begin{aligned}&\Bigl\langle \exp\bigl(-\hat{\boldsymbol{h}}^{\mathrm{T}}\boldsymbol{J}\,\boldsymbol{x}\bigr)\Bigr\rangle_{\boldsymbol{J} \overset{\text{i.i.d.}}{\sim} \mathcal{N}(\frac{\bar{g}}{N}, \frac{1}{N})} \\ &= \exp\Bigl(-\frac{\bar{g}}{N}\sum_{i=1}^{N}\hat{h}_i^{\mathrm{T}}\sum_{j=1}^{N} x_j + \frac{1}{2N}\sum_{i,j=1}^{N}\bigl(\hat{h}_i^{\mathrm{T}} x_j\bigr)^2\Bigr) \\ &= \prod_{i=1}^{N} \exp\Bigl(-\hat{h}_i^{\mathrm{T}}\mathcal{R}\bar{g} + \frac{1}{2}\hat{h}_i^{\mathrm{T}}\mathcal{Q}\hat{h}_i\Bigr)\end{aligned} \tag{4}$$

motivates the definition of the fields

$$\mathcal{R}(t) := \frac{1}{N}\sum_{j=1}^{N} x_j(t) \quad \text{and} \quad \mathcal{Q}(t,s) := \frac{1}{N}\sum_{j=1}^{N} x_j(t) x_j(s). \tag{5}$$

The scalar products are again

$$\hat{h}_i^{\mathrm{T}}\mathcal{R} = \int dt \hat{h}_i(t) R(t) \quad \text{or} \quad \hat{h}_i^{\mathrm{T}}\mathcal{Q}\hat{h}_i = \iint dt\, ds\, \hat{h}_i(t)\mathcal{Q}(t,s)\hat{h}_i(s).$$

Enforcing the definitions in equations (5) by Dirac-$\delta$ constraints gives rise to (imaginary) auxiliary fields [4, Chapter 12] $\hat{\mathcal{R}}$ and $\hat{Q}$, leaving us with the disorder averaged functional

$$
\begin{aligned}
\big\langle\rho[\boldsymbol{h}]\big\rangle_{\boldsymbol{J}} &= \int \mathcal{D}\boldsymbol{x}\,\rho[\boldsymbol{x}|\boldsymbol{h}]\,\big\langle\delta[\boldsymbol{h}-\boldsymbol{J}\boldsymbol{x}]\big\rangle_{\boldsymbol{J}} \\
&= \int \mathcal{D}\{\mathcal{Q},\mathcal{R},\hat{\mathcal{Q}},\hat{\mathcal{R}}\}\exp\big(-N\hat{\mathcal{R}}^{\mathrm{T}}\mathcal{R}-N\hat{\mathcal{Q}}^{\mathrm{T}}\mathcal{Q}\big) \\
&\times\prod_{i=1}^{N}\int \mathcal{D}\{x_i,\hat{h}_i\}\,\rho[x_i|h_i] \\
&\times\exp\Big(\hat{h}_i^{\mathrm{T}}h_i-\bar{g}\hat{h}_i^{\mathrm{T}}\mathcal{R}+\frac{1}{2}\,\hat{h}_i^{\mathrm{T}}\mathcal{Q}\hat{h}_i+\hat{\mathcal{R}}^{\mathrm{T}}x_i+x_i^{\mathrm{T}}\hat{\mathcal{Q}}x_i\Big).
\end{aligned}
\tag{6}
$$

Note that the integral over components of $\boldsymbol{x}$, $\boldsymbol{h}$, and $\hat{\boldsymbol{h}}$ factorizes and integrates out the microscopic degrees of freedom, allowing us to extract a common factor $N$ and state the action $\Omega$ as

$$
\begin{aligned}
&\int \mathcal{D}\boldsymbol{h}\,\big\langle\rho[\boldsymbol{h}]\big\rangle_{\boldsymbol{J}} \\
&= \int \mathcal{D}\{\mathcal{Q},\mathcal{R},\hat{\mathcal{Q}},\hat{\mathcal{R}}\}\exp\big(N\,\Omega[\mathcal{R},\mathcal{Q},\hat{\mathcal{R}},\hat{\mathcal{Q}}]\big),
\end{aligned}
$$

$$
\Omega[\mathcal{R},\mathcal{Q},\hat{\mathcal{R}},\hat{\mathcal{Q}}] := -\mathcal{R}^{\mathrm{T}}\hat{\mathcal{R}}-\mathcal{Q}^{\mathrm{T}}\hat{\mathcal{Q}}+\ln\int \mathcal{D}\{x,h,\hat{h}\}\,\rho[x|h]\exp\Big(\hat{h}^{\mathrm{T}}h-\bar{g}\hat{h}^{\mathrm{T}}\mathcal{R}+\frac{1}{2}\,\hat{h}^{\mathrm{T}}\mathcal{Q}\hat{h}+\hat{\mathcal{R}}^{\mathrm{T}}x+x^{\mathrm{T}}\hat{\mathcal{Q}}x\Big). \tag{7}
$$

We now want to find those values of the fields $\mathcal{R},\mathcal{Q},\hat{\mathcal{R}},\hat{\mathcal{Q}}$ called $R,Q,\hat{R},\hat{Q}$ that have the largest contribution to the probability mass which amounts to doing a saddle point approximation. The probability concentrates around the maximum of $\Omega$ for large $N$. This leads to

$$
\begin{aligned}
0 &= \frac{\delta\Omega[\mathcal{R},\mathcal{Q},\hat{\mathcal{R}},\hat{\mathcal{Q}}]}{\delta\hat{\mathcal{R}}(t)}\Big|_{\substack{\mathcal{Q}=Q,\hat{\mathcal{Q}}=\hat{Q}\\ \mathcal{R}=R,\hat{\mathcal{R}}=\hat{R}}} = -R(t)+\langle x(t)\rangle_{\Omega(\bar{g}R,Q)}\ , \\
0 &= \frac{\delta\Omega[\mathcal{R},\mathcal{Q},\hat{\mathcal{R}},\hat{\mathcal{Q}}]}{\delta\mathcal{R}(t)}\Big|_{\substack{\mathcal{Q}=Q,\hat{\mathcal{Q}}=\hat{Q}\\ \mathcal{R}=R,\hat{\mathcal{R}}=\hat{R}}} = -\hat{R}(t)-\underbrace{\big\langle\hat{h}(t)\big\rangle_{\Omega(\bar{g}R,Q)}}_{=0}, \\
0 &= \frac{\delta\Omega[\mathcal{R},\mathcal{Q},\hat{\mathcal{R}},\hat{\mathcal{Q}}]}{\delta\hat{\mathcal{Q}}(t,s)}\Big|_{\substack{\mathcal{Q}=Q,\hat{\mathcal{Q}}=\hat{Q}\\ \mathcal{R}=R,\hat{\mathcal{R}}=\hat{R}}} = -Q(t,s)+\langle x(t)x(s)\rangle_{\Omega(\bar{g}R,Q)}\ , \\
0 &= \frac{\delta\Omega[\mathcal{R},\mathcal{Q},\hat{\mathcal{R}},\hat{\mathcal{Q}}]}{\delta\mathcal{Q}(t,s)}\Big|_{\substack{\mathcal{Q}=Q,\hat{\mathcal{Q}}=\hat{Q}\\ \mathcal{R}=R,\hat{\mathcal{R}}=\hat{R}}} = -\hat{Q}(t,s)+\frac{1}{2}\underbrace{\big\langle\hat{h}(t)\hat{h}(s)\big\rangle_{\Omega(\bar{g}R,Q)}}_{=0},
\end{aligned}
$$

where all the expectation values of $\hat{h}$ vanish as detailed in [4, Sec 10.1]. The average $\langle\dots\rangle_{\Omega(\bar{g}R,Q)}$ is taken with respect to the single-site action,

$$
\begin{aligned}
\langle\dots\rangle_{\Omega(\bar{g}R,Q)} &= \int \mathcal{D}\{x,h,\hat{h}\}\,\rho[x|h]\,\dots\,\exp\Big(\hat{h}^{\mathrm{T}}h-\bar{g}\hat{h}^{\mathrm{T}}\mathcal{R}+\frac{1}{2}\,\hat{h}^{\mathrm{T}}\mathcal{Q}\hat{h}+\hat{\mathcal{R}}^{\mathrm{T}}x+x^{\mathrm{T}}\hat{\mathcal{Q}}x\Big)\Big|_{\substack{\mathcal{Q}=Q,\hat{\mathcal{Q}}=\hat{Q}\\ \mathcal{R}=R,\hat{\mathcal{R}}=\hat{R}}} \\
&= \int \mathcal{D}\{x,h\}\,\rho[x|h]\cdots\prod_{t}\int_{-i\infty}^{i\infty}\frac{d\hat{h}(t)}{2\pi i}\exp\Big(\hat{h}^{\mathrm{T}}h-\bar{g}\hat{h}^{\mathrm{T}}R+\frac{1}{2}\,\hat{h}^{\mathrm{T}}Q\hat{h}\Big) \\
&= \int \mathcal{D}\{x,h\}\,\rho[x|h]\,\dots\,\mathcal{N}(h|\bar{g}R,Q) \\
&= \int \mathcal{D}x\,\dots\,\langle\rho[x|h]\rangle_{h\sim\mathcal{N}(\bar{g}R(t),Q(t,s))}\ ,
\end{aligned}
\tag{8}
$$

where in the penultimate line, we identified the cumulant generating function of a Gaussian. Through this field formalism, we obtained mean-field values for the fields $\mathcal{R}$ and $\mathcal{Q}$,

$$\begin{aligned} R(t) &= \langle x(t)\rangle_{\Omega(\bar{g}R(t),Q(t,s))} \\ Q(t,s) &= \langle x(t)x(s)\rangle_{\Omega(\bar{g}R(t),Q(t,s))} \,. \end{aligned} \tag{9}$$

The variance $Q(t,s)$ in general couples different points in time rendering equation (9) non-local in time. These two equations provide a description of the network in terms of a single neuron $x$ representative of the whole network.

#### 2.2 Two-replica calculation

In the experiment, we compare different trials to each other. To account for this feature, we employ a two-replica calculation, where each replicon can be understood as one trial $\alpha$ of a stimulus presentation. The difference to the calculations before is that for the replica calculation, we take two instances of networks with the same realization of the connectivity $\boldsymbol{J}$ and then perform a disorder average over the randomness in $\boldsymbol{J}$. Later, we will see that this results in a newly appearing term which shapes the joint time evolution of the networks.

Because of the two replicas, the functional equation (1) is reformulated with two fields $\boldsymbol{h}^{(\alpha)}$ and neuron states $\boldsymbol{x}^{(\alpha)}$

$$\rho[\boldsymbol{x}^{(1)},\boldsymbol{x}^{(2)}|\boldsymbol{h}^{(1)},\boldsymbol{h}^{(2)}] = \prod_{i=1}^{N}\rho[x_i^{(1)},x_i^{(2)}|h_i^{(1)},h_i^{(2)}].$$

The disorder average is now taken over the functional

$$\begin{aligned} \rho[\boldsymbol{h}^{(1)},\boldsymbol{h}^{(2)}] = \int \mathcal{D}\{\boldsymbol{x}^{(1)},\boldsymbol{x}^{(2)}\}\, &\rho[\boldsymbol{x}^{(1)},\boldsymbol{x}^{(2)}|\boldsymbol{h}^{(1)},\boldsymbol{h}^{(2)}] \\ &\times \prod_{\alpha=1}^{2}\delta[\boldsymbol{h}^{(\alpha)} - \boldsymbol{J}\,\boldsymbol{x}^{(\alpha)}], \end{aligned} \tag{10}$$

which gives rise to a coupling term $\hat{h}_i^{(1)\mathrm{T}}x_j^{(1)}\,\hat{h}_i^{(2)\mathrm{T}}x_j^{(2)}$ between the networks

$$\begin{aligned} &\Big\langle \exp\big(-\hat{\boldsymbol{h}}^{(1)\mathrm{T}}\boldsymbol{J}\boldsymbol{x}^{(1)} - \hat{\boldsymbol{h}}^{(2)\mathrm{T}}\boldsymbol{J}\boldsymbol{x}^{(2)}\big)\Big\rangle_{\boldsymbol{J}\overset{\text{i.i.d.}}{\sim}\mathcal{N}(\frac{\bar{g}}{N},\frac{1}{N})} \\ &= \prod_{i=1}^{N}\prod_{\alpha=1}^{2}\exp\Big(-\frac{\bar{g}}{N}\hat{h}_i^{(\alpha)\mathrm{T}}\sum_{j=1}^{N}x_j^{(\alpha)} + \frac{1}{2}\frac{1}{N}\sum_{j=1}^{N}\big(\hat{h}_i^{(\alpha)\mathrm{T}}x_j^{(\alpha)}\big)^2\Big) \\ &\quad\times\exp\Big(\frac{1}{N}\sum_{j=1}^{N}\hat{h}_i^{(1)\mathrm{T}}x_j^{(1)}\,\hat{h}_i^{(2)\mathrm{T}}x_j^{(2)}\Big). \end{aligned} \tag{11}$$

The last line suggests the introduction of an additional field $\mathcal{Q}^{(12)}$. This definition is again enforced by a $\delta$-constraint, yielding one additional auxiliary field. This motivates a unified notation

$$\mathcal{Q}^{(\alpha\beta)}(t,s) \coloneqq \frac{1}{N}\sum_{j=1}^{N}x_j^{(\alpha)}(t)\,x_j^{(\beta)}(s)$$

for the cross- and autocorrelation of the replicas with $\alpha,\beta\in\{1,2\}$. Similar to the previous section, we can now formulate a disorder averaged functional with an action $\Omega$ of which we calculate the saddle point. Using the saddle point approximation, one arrives at the mean-field equations

$$
\begin{aligned}
R^{(\alpha)}(t) &= \langle x^{(\alpha)}(t) \rangle_{\Omega(\{\bar{g}R^{(\alpha)}, Q^{(\alpha\beta)}\})} \\
Q^{(\alpha\beta)}(t,s) &= \langle x^{(\alpha)}(t) x^{(\beta)}(s) \rangle_{\Omega(\{\bar{g}R^{(\alpha)}, Q^{(\alpha\beta)}\})}
\end{aligned}
$$

with

$$
\langle \dots \rangle_{\Omega(\{\bar{g}R^{(\alpha)}, Q^{(\alpha\beta)}\})} = \int \mathcal{D}\{x^{(1)}, x^{(2)}\} \dots \langle \rho[x^{(1)}, x^{(2)} | h^{(1)}, h^{(2)}] \rangle_{(h^{(1)}, h^{(2)}) \sim \mathcal{N}(\{\bar{g}R^{(\alpha)}, Q^{(\alpha\beta)}\})}, \tag{12}
$$

where $\mathcal{N}(\{\bar{g}R^{(\alpha)}, Q^{(\alpha\beta)}\})$ is a shorthand for the bivariate Gaussian $\mathcal{N}\left(\begin{pmatrix} \bar{g}R^{(1)} \\ \bar{g}R^{(2)} \end{pmatrix}, \begin{pmatrix} Q^{(11)} & Q^{(12)} \\ Q^{(21)} & Q^{(22)} \end{pmatrix}\right)$. We now have to specify $\rho[x^{(1)}, x^{(2)} | h^{(1)}, h^{(2)}]$ to tailor this general framework to our model of Boolean neurons.

#### 2.3 Time evolution of the mean-field quantities

In the calculations so far, we have not yet specified our neuron model. As mentioned at the beginning of this section, we take binary neurons with Boolean values $x_i \in \{0, 1\}$ instead of Ising values $x_i \in \{1, -1\}$ as in [3]. The time dynamics are governed by specifically chosen update times that serve as possible points in time allowing one neuron to switch its state (Glauber dynamics [5]). We assume that updates are equally likely between any two time intervals of equal length. Hence, they follow a Poisson process which is specified by stating the probability distribution of intervals $\Delta t$ between two update times to be

$$
p(\Delta t = t) = \frac{1}{\tau} e^{-\frac{t}{\tau}}
$$

with the Poisson rate given by the time constant $\tau^{-1}$. The mean of this distribution $\tau$ reflects the characteristic timescale on which an active neuron affects the receiving neurons. This is because once updated, a neuron will remain active until the next time step of update, after which it will most likely deactivate provided the activity is low. The probability for a neuron $i$ to be active at some time point $t$ only depends on the field $h_i^{(\alpha)}$ of the replicon it belongs to, and is influenced by the probability to be active at a prior time point $t'$

$$
\rho[x_i^{(\alpha)}(t) = 1 | h_i^{(\alpha)}] = \int_{-\infty}^{t} \frac{dt'}{\tau} e^{-\frac{t-t'}{\tau}} H\left(h_i^{(\alpha)}(t')\right).
$$

$H(\circ)$ is the activation function, which is chosen to be the deterministic Heaviside function, i.e. an update to the active state always happens if the input is large enough to cross the threshold (here 0). Because the Heaviside function has no scale, we can choose the variance of the connectivity matrix (equation (3)) to be $1/N$ without loss of generality. We can now evaluate equation (9) by using the above equation for binary neurons:

$$
\begin{aligned}
R^{(\alpha)}(t) =& \langle x^{(\alpha)}(t) \rangle_{\Omega(\{\bar{g}R^{(\alpha)}, Q^{(\alpha\beta)}\})} \\
=& 0 \cdot \langle \rho[x^{(\alpha)}(t) = 0 | h^{(1)}, h^{(2)}] \rangle_{(h^{(1)}, h^{(2)}) \sim \mathcal{N}(\{\bar{g}R^{(\alpha)}, Q^{(\alpha\beta)}\})} \\
&+ 1 \cdot \langle \rho[x^{(\alpha)}(t) = 1 | h^{(1)}, h^{(2)}] \rangle_{(h^{(1)}, h^{(2)}) \sim \mathcal{N}(\{\bar{g}R^{(\alpha)}, Q^{(\alpha\beta)}\})} \\
=& \langle \rho[x^{(\alpha)}(t) = 1 | h^{(\alpha)}] \rangle_{h^{(\alpha)} \sim \mathcal{N}(\bar{g}R^{(\alpha)}, Q^{(\alpha\alpha)})} \\
=& \int_{-\infty}^{t} \frac{dt'}{\tau} e^{-\frac{t-t'}{\tau}} \left\langle H\left(h^{(\alpha)}\right) \right\rangle_{h^{(\alpha)} \sim \mathcal{N}(\bar{g}R^{(\alpha)}(t'), Q^{(\alpha\alpha)}(t',t'))}.
\end{aligned} \tag{13}
$$

The mere appearance of the equal-time autocorrelation $Q^{(\alpha\alpha)}(t,t)$ mirrors the Markov property of the system; the time evolution at time point $t$ does not depend on the history, but rather only on the statistics of this very time point. Note here that due to our choice of Boolean values, the equal-time autocorrelation can be written as

$$Q^{(\alpha\alpha)}(t,s=t)=\left\langle x^{(\alpha)}(t)x^{(\alpha)}(t)\right\rangle_{\Omega(\{\bar{g}R^{(\alpha)},Q^{(\alpha\beta)}\})}=R^{(\alpha)}(t). \tag{14}$$

Taking the time derivative of (13), we obtain

$$\tau\frac{d}{dt}R^{(\alpha)}(t)+R^{(\alpha)}(t)=\langle H(h)\rangle_{h\sim\mathcal{N}(\bar{g}R^{(\alpha)}(t),Q^{(\alpha\alpha)}(t,t))}\,. \tag{15}$$

Next, we want to derive an equation for the equal-time cross-correlation. For the product $x^{(1)}(t)x^{(2)}(t)$ to become zero, it is sufficient if one of the $H\left(h^{(\alpha)}(t)\right)$ is zero. Thus the minimum of these activation functions determines the transition and it follows for the integral over $\{x^{(1)},x^{(2)}\}$ in equation (12)

$$\begin{aligned}
&\sum_{x^{(1)}(t),x^{(2)}(t)=0}^{1} x^{(1)}(t)x^{(2)}(t)\rho[x^{(1)}(t),x^{(2)}(t)|h^{(1)},h^{(2)}]\\
&=\int_{-\infty}^{t}\frac{dt'}{\tau}e^{-\frac{t-t'}{\tau}}\min\{H(h^{(1)}(t')),H(h^{(2)}(t'))\}\\
&=\int_{-\infty}^{t}\frac{dt'}{\tau}e^{-\frac{t-t'}{\tau}}H(h^{(1)}(t'))H(h^{(2)}(t')).
\end{aligned}$$

Taking into account the average over the fields, we obtain

$$\begin{aligned}
Q^{(12)}(t,t)=&\langle x^{(1)}(t)x^{(2)}(t)\rangle_{\Omega(\{\bar{g}R^{(\alpha)},Q^{(\alpha\beta)}\})}\\
=&\sum_{x^{(1)}(t),x^{(2)}(t)=0}^{1} x^{(1)}(t)x^{(2)}(t)\langle\rho[x^{(1)}(t),x^{(2)}(t)|h^{(1)},h^{(2)}]\rangle_{(h^{(1)},h^{(2)})\sim\mathcal{N}(\{\bar{g}R^{(\alpha)},Q^{(\alpha\beta)}\})}\\
=&\int_{-\infty}^{t}\frac{dt'}{\tau}e^{-\frac{t-t'}{\tau}}\left\langle H(h^{(1)})H(h^{(2)})\right\rangle_{(h^{(1)},h^{(2)})\sim\mathcal{N}(\{\bar{g}R^{(\alpha)},Q^{(\alpha\beta)}\})}.
\end{aligned}$$

Taking the derivative with respect to time yields

$$\tau\frac{d}{dt}Q^{(12)}(t,t)+Q^{(12)}(t,t)=\left\langle H(h^{(1)})H(h^{(2)})\right\rangle_{(h^{(1)},h^{(2)})\sim\mathcal{N}(\{\bar{g}R^{(\alpha)},Q^{(\alpha\beta)}\})}. \tag{16}$$

The equal-time cross-correlation $Q^{(12)}(t,t)$ is a similarity measure between the neuron states $\boldsymbol{x}^{(1)}$ and $\boldsymbol{x}^{(2)}$ of the two replicas. In the following we refer to this $Q^{(12)}(t,t)$ as the "overlap".

Note how the population mean enters the time evolution equation of the correlation in the mean and variance of the bivariate Gaussian. For simplicity, we assume that the responses in different trials are similar in strength $R^{(1)}=R^{(2)}=R$. Because of the Boolean values, $0\le Q^{(12)}\le R$, and, as anticipated for perfect correlation $Q^{(12)}=R$, the correlation equation becomes that of the mean. Also note how the correlation for unequal time points does not show up, so these ordinary differential equations (ODEs) $\{d/dt\,R^{(\alpha)}(t),\,d/dt\,Q^{(\alpha\beta)}(t)\}$ are closed as a result of the Markov property.

#### 2.4 Network parameters

Here we provide details on the parametrization of the network model, which will enable us to connect it to the experimental data.

##### 2.4.1 Thresholds

We introduce thresholds by adding a static, external field $h_{\mathrm{stat},i} \overset{\text{i.i.d.}}{\sim} \mathcal{N}(\delta,\gamma)$ to every neuron, where we only fix its mean $\delta$ and variance $\gamma$ across neurons. We assume that the thresholds are constant over the time course of the experiment (or between trials) and unaffected by the stimulus applied, making it the same for each replicon. The external fields change as $\boldsymbol{h}^{(\alpha)} \to \boldsymbol{h}^{(\alpha)} + \boldsymbol{h}_{\text{stat.}}$.

The additional term in the disorder average

$$\begin{aligned}
&\Big\langle \exp\big( - \hat{\boldsymbol{h}}^{(1)T}\boldsymbol{h}_{\text{stat.}} - \hat{\boldsymbol{h}}^{(2)T}\boldsymbol{h}_{\text{stat.}}\big)\Big\rangle_{h_{\mathrm{stat},i} \overset{\text{i.i.d.}}{\sim} \mathcal{N}(\delta,\gamma)} \\
&= \prod_{i=1}^{N_{\mathrm{nrn}}} \prod_{\alpha=1}^{2} \exp\left(-\delta\, \hat{h}_i^{(\alpha)\mathrm{T}} 1 + \frac{\gamma}{2} \hat{h}_i^{(\alpha)\mathrm{T}} 1 \hat{h}_i^{(\alpha)}\right) \\
&\quad \times \exp\Big(\gamma\, \hat{h}_i^{(1)\mathrm{T}} 1 \hat{h}_i^{(2)}\Big),
\end{aligned}$$

with

$$\hat{h}_i^{(\alpha)\mathrm{T}} 1 = \int dt\, \hat{h}_i^{(\alpha)}(t) \qquad \text{or} \qquad \hat{h}_i^{(\alpha)\mathrm{T}} 1 \hat{h}_i^{(\alpha)} = \int ds \int dt\, \hat{h}_i^{(\alpha)}(s) \hat{h}_i^{(\alpha)}(t).$$

These additional terms are accounted for by replacing the Gaussian measure as

$$\mathcal{N}\left(\begin{pmatrix} \bar{g}R \\ \bar{g}R \end{pmatrix}, \begin{pmatrix} R & Q^{(12)} \\ Q^{(21)} & R \end{pmatrix}\right) \to \mathcal{N}\left(\begin{pmatrix} \bar{g}R + \delta \\ \bar{g}R + \delta \end{pmatrix}, \begin{pmatrix} R + \gamma & Q^{(12)} + \gamma \\ Q^{(21)} + \gamma & R + \gamma \end{pmatrix}\right),$$

because the cumulants of the independently distributed Gaussian fields add up for the sum of those fields $\boldsymbol{h}^{(\alpha)} + \boldsymbol{h}_{\text{stat.}}$.

##### 2.4.2 External stimulus

In addition to the static field contribution, we want to be able to induce transient behavior by sending a stimulus into the replica that carries the input information for different trials. We here take a boxcar stimulus $f(t)$

$$f(t) = \begin{cases} 1 & 0 \le t < t_{\max} \\ 0 & \text{else} \end{cases}$$

multiplied by two different vectors $\boldsymbol{v}^{(1)}$ and $\boldsymbol{v}^{(2)}$ whose entries are correlated with a parameter $\nu^{\circlearrowright} \in [0,1]$ if the replica represent trials within one stimulus class, and $\nu^{\leftrightarrow} \in [0,1]$ otherwise ($\nu^{\leftrightarrow} < \nu^{\circlearrowright}$).The stimulus vectors are drawn according to a Gaussian

$$\begin{pmatrix} v_i^{(1)} \\ v_i^{(2)} \end{pmatrix} \overset{\text{i.i.d.}}{\sim} \mathcal{N}\left(\begin{pmatrix} \eta \\ \eta \end{pmatrix}, \Theta \begin{pmatrix} 1 & \nu^{\circ} \\ \nu^{\circ} & 1 \end{pmatrix}\right), \tag{17}$$

where $\circ \in \{\leftrightarrow, \circlearrowright\}$, $\eta$ is the average input strength, and $\Theta$ is the overall scale of the variability in input.

The fields correspondingly get transformed as

$$\boldsymbol{h}^{(\alpha)} + \boldsymbol{h}_{\text{stat.}} \to \boldsymbol{h}^{(\alpha)} + \boldsymbol{h}_{\text{stat.}} + \boldsymbol{v}^{(\alpha)} f(t). \tag{18}$$

This transforms the bivariate Gaussian of the fields

$$
\begin{aligned}
&\mathcal{N}\left(\begin{pmatrix}\bar{g}R+\delta\\ \bar{g}R+\delta\end{pmatrix},\begin{pmatrix}R+\gamma & Q^{(12)}+\gamma\\ Q^{(12)}+\gamma & R+\gamma\end{pmatrix}\right)\\
\to&\mathcal{N}\left(\begin{pmatrix}\bar{g}R+\delta+\eta\\ \bar{g}R+\delta+\eta\end{pmatrix},\begin{pmatrix}R+\gamma+\Theta & Q^{(12)}+\gamma+\nu\Theta\\ Q^{(12)}+\gamma+\nu\Theta & R+\gamma+\Theta\end{pmatrix}\right).
\end{aligned}
$$

The parameters $\eta, \Theta, \nu^{\leftrightarrow}, \nu^{\circlearrowright}$ allow us to fit the peak-heights of the experimental observables.

###### 2.4.3 Mean-field equations

The external stimulus fixes the initial state and leads to two different overlaps $Q^{\circlearrowright}$ and $Q^{\leftrightarrow}$ depending on the class membership of the trials. Subsequently, the stimulus is switched off and the recurrent dynamics evolve autonomously. The time-evolution is then governed by

$$
\begin{aligned}
\tau\frac{d}{dt}R &= -R+\mathcal{T}(R,R)\,,\\
\tau\frac{d}{dt}Q^{\circlearrowright} &= -Q^{\circlearrowright}+\mathcal{T}(R,Q^{\circlearrowright})\,,\\
\tau\frac{d}{dt}Q^{\leftrightarrow} &= -Q^{\leftrightarrow}+\mathcal{T}(R,Q^{\leftrightarrow})\,,
\end{aligned}
\tag{19}
$$

where we introduced the correlation-transmission function $\mathcal{T}(\circ,\circ)$, defined and evaluated through equations (15), (16) and the additional network parameters

$$
\begin{aligned}
\mathcal{T}(R,R) &= \langle H(h)\rangle_{h\sim\mathcal{N}(\bar{g}R+\delta,R+\gamma)}\\
&= \frac{1}{2}\left(1+\operatorname{erf}\left(\frac{\bar{g}R+\delta}{\sqrt{2}\sqrt{R+\gamma}}\right)\right).
\end{aligned}
\tag{20}
$$

Thus, $\mathcal{T}(R,R)$ is a function of $c_1 \coloneqq \frac{\bar{g}R+\delta}{\sqrt{R+\gamma}}$. For the overlaps, we have

$$
\begin{aligned}
\mathcal{T}(R,Q) &= \left\langle H(h^{(1)})H(h^{(2)})\right\rangle_{\left(\begin{smallmatrix}h^{(1)}\\ h^{(2)}\end{smallmatrix}\right)\sim\mathcal{N}\left(\left(\begin{smallmatrix}\bar{g}R+\delta\\ \bar{g}R+\delta\end{smallmatrix}\right),\left(\begin{smallmatrix}R+\gamma & Q+\gamma\\ Q+\gamma & R+\gamma\end{smallmatrix}\right)\right)}\\
&= \left[G(c_1)-2T\left(c_1,\sqrt{\frac{1-c_2}{1+c_2}}\right)\right]\Bigg|_{c_1=\frac{\bar{g}R+\delta}{\sqrt{R+\gamma}},\,c_2=\frac{Q+\gamma}{R+\gamma}}
\end{aligned}
\tag{21}
$$

with $c_1 = \frac{\bar{g}R+\delta}{\sqrt{R+\gamma}}$ and $c_2 = \frac{Q+\gamma}{R+\gamma}$ and $G(x) = \frac{1}{2}\left(1+\operatorname{erf}\left(\frac{x}{\sqrt{2}}\right)\right)$ and Owen's T function $T(h,a) = \frac{1}{2\pi}\int_0^a dx\,\frac{\exp(-\frac{1}{2}h^2(1+x^2))}{1+x^2}$ [6]. We thereby extracted the arguments $c_1$ and $c_2$, on which $\mathcal{T}(R,Q)$ depends. We have used here

$$
\begin{aligned}
&\left\langle H(h^{(1)})H(h^{(2)})\right\rangle_{\left(\begin{smallmatrix}h^{(1)}\\ h^{(2)}\end{smallmatrix}\right)\sim\mathcal{N}\left(\left(\begin{smallmatrix}R\\ R\end{smallmatrix}\right),\left(\begin{smallmatrix}Q_1 & Q_2\\ Q_2 & Q_1\end{smallmatrix}\right)\right)}\\
=&G\left(\frac{R}{\sqrt{Q_1}}\right)-2T\left(\frac{R}{\sqrt{Q_1}},\sqrt{\frac{1-Q_2/Q_1}{1+Q_2/Q_1}}\right).
\end{aligned}
\tag{22}
$$

We can obtain equation (22) by taking $\beta\to\infty$ of the more general expression

$$
\begin{aligned}
&\left\langle G(\beta h^{(1)})G(\beta h^{(2)})\right\rangle_{\begin{pmatrix}h^{(1)}\\h^{(2)}\end{pmatrix}\sim\mathcal{N}\left(\begin{pmatrix}R\\R\end{pmatrix},\begin{pmatrix}Q_1 & Q_2\\Q_2 & Q_1\end{pmatrix}\right)}\\
&=G\left(\frac{R}{\sqrt{\beta^{-2}+Q_1}}\right)-2T\left(\frac{R}{\sqrt{\beta^{-2}+Q_1}},\sqrt{\frac{\beta^{-2}+1-Q_2/Q_1}{\beta^{-2}+1+Q_2/Q_1}}\right),
\end{aligned}
$$

see [7].

Knowing that $\mathcal{T}(R,Q)$ depends on $c_1$ and $c_2$ will facilitate the process of fitting the model to the data.

### 3 Fitting the model to experimental data

In this section we will show how the mean-field time-evolution equations are fit to the experimental data using the parameters $\tau,\bar{g},\delta,\gamma$ introduced in the previous section. We distinguish between fitting the baseline values, the highest (peak) values, and the decay of the three quantities. We indicate experimental values with a superscript $\circ^{\mathrm{exp}}$.

#### 3.1 Fitting the baselines

The baselines are indicated with a subscript $\circ_0$. We obtain baseline values for the observables by averaging over the time points $\{t_0\}$ after the transients where $t_0 > 1.5\mathrm{s}$.

In addition to averaging over the neurons $i$, we average over trials $\alpha$ for the population mean,

$$
R_0^{\mathrm{exp}} = \langle\langle\langle x_{\alpha,i}(t)\rangle_i\rangle_{t\in\{t_0\}}\rangle_\alpha .
$$

For the baselines of the correlation, we average over pairs of trials $(\alpha\beta)$,

$$
Q_0^{\mathrm{exp}} = \langle\langle\langle x_{\alpha,i}(t)x_{\beta,i}(t)\rangle_i\rangle_{t\in\{t_0\}}\rangle_{(\alpha\beta)|\,\alpha\neq\beta} .
$$

Since both $Q^{\circlearrowleft}$ and $Q^{\leftrightarrow}$ relax to the same baseline after the transient we do not have to distinguish by the classes of trials $\alpha$ and $\beta$. We seek to find parameters $\bar{g},\delta,\gamma$ such that

$$
\begin{aligned}
R_0^{\mathrm{exp}} &= \mathcal{T}(R_0^{\mathrm{exp}},R_0^{\mathrm{exp}})\\
Q_0^{\mathrm{exp}} &= \mathcal{T}(R_0^{\mathrm{exp}},Q_0^{\mathrm{exp}}).
\end{aligned}
\tag{23}
$$

In equations (20), (21) we see that the correlation transmission function $\mathcal{T}$ only depends on the two constellations of parameters $c_1$ and $c_2$. To fulfill equation (23) we have to find these values. We can infer the baseline value $c_{1,0}$ straightforwardly by

$$
c_{1,0} = \sqrt{2}\,\mathrm{erf}^{-1}\left(2R_0^{\mathrm{exp}}-1\right).
$$

Because we only have an integral representation $\mathcal{T}(R,Q)$ for the correlations, we use a bisection-algorithm (Figure 1) to determine a set of parameters that fulfills equations (23). The underlying notion is that $\bar{g}$ tunes the inhibition and $\bar{g}R$ displaces the Gaussian of the input field $h$, whereas $\gamma$ broadens this Gaussian independent of $R$. So for a given left-hand side in the second equation in (23) and $\bar{g}$ inferred from the population mean baseline, one can find a $\gamma(\bar{g})$ that reproduces the baselines. We choose $\delta = 0$ as an initial parameter guess for the algorithm in

**def** fit_baselines_with_$\bar{g}$_and_$\gamma$($R_0^{\text{exp}}$, $Q_0^{\text{exp}}$, $\delta = 0$):

  $c_{1,0} = \sqrt{2}\,\text{erf}^{-1}(2\,R_0^{\text{exp}} - 1)$

  **def** root_corr($\gamma$):
    $\bar{g} = (c_{1,0}\sqrt{R_0^{\text{exp}} + \gamma} - \delta)/R_0^{\text{exp}}$
    $z = R_0(\bar{g}, \delta, \gamma)$
    **return** $Q_0(\bar{g}, \delta, \gamma, R = z) - Q_0^{\text{exp}}$

  $\gamma$ = bisect(root_corr, $\gamma_{\text{begin}}$, $\gamma_{\text{end}}$)
  $\bar{g} = (c_{1,0}\sqrt{R_0^{\text{exp}} + \gamma} - \delta)/R_0^{\text{exp}}$

  **return** $\bar{g}$, $\gamma$

Figure 1: **Bisection algorithm determining parameters that fit experimental baselines.** $R_0$ and $Q_0$ are obtained by integrating forward the equations (19) until a stationary point is reached. Initial guess for threshold $\delta = 0$. Start and end values $\gamma_{\text{begin}}, \gamma_{\text{end}}$ can be chosen to lie within $[0, \infty]$.

Figure 1 since for negative $\delta$ the self-consistent equation in (23) can have two solutions when a quiescent fixed-point ($R = 0$) arises.

Having found a set $\{\tau, \bar{g}, \delta, \gamma\}$ that fulfills equation (23) we can obtain all other sets that reproduce the baselines by demanding that $c_1$ and $c_2$ remain constant. Note that $\gamma$ is fixed by the baseline constraints

$$\gamma = \sqrt{\frac{R_0^{\text{exp}} c_{2,0} - Q_0^{\text{exp}}}{1 - c_{2,0}}}$$

while either $\bar{g}$ or $\delta$ is still free to choose

$$\delta = c_{1,0}\sqrt{R_0^{\text{exp}} + \gamma} - \bar{g} R_0^{\text{exp}}. \tag{24}$$

#### 3.2 Fitting the decay

Reproducing the experimental baselines leaves us with one free parameter ($\delta$ or $\bar{g}$) in addition to the time constant $\tau$. We will fit the time-course for the times $\{t_{\text{decay}}\}$ where $t_{\max} \leq t_{\text{decay}} \leq t_{\max} + 0.4$s, because we see a transient in activity in this interval.

The experimental population mean is obtained by

$$R^{\text{exp}}(t) = \langle\langle x_{\alpha,i}(t)\rangle_i\rangle_\alpha$$

and the experimental overlaps by

$$Q^{\circlearrowright,\text{exp}}(t) = \langle\langle x_{\alpha,i}(t) x_{\beta,i}(t)\rangle_i\rangle_{(\alpha\beta)|\,C(\alpha)=C(\beta)}\,,$$
$$Q^{\leftrightarrow,\text{exp}}(t) = \langle\langle x_{\alpha,i}(t) x_{\beta,i}(t)\rangle_i\rangle_{(\alpha\beta)|\,C(\alpha)\neq C(\beta)}\,,$$

where we average over pairs of trials that have the same/different class membership. In general the overlap $Q^{\circlearrowright,\text{exp}}$ (intra-overlap) is larger than $Q^{\leftrightarrow,\text{exp}}$ (inter-overlap), meaning that trials of the same class are more similar than trials of different classes.

We fit $\tau, \delta$ by minimizing the quadratic loss

$$\mathcal{L}(\tau, \delta) = \sum_{t \in \{t_{\text{decay}}\}} \left[ \frac{(R(t) - R^{\text{exp}}(t))^2}{\sigma_R^2(t)} + \frac{\left(Q^{\circlearrowleft}(t) - Q^{\circlearrowleft,\text{exp}}(t)\right)^2}{\sigma^2_{Q^{\circlearrowleft,\text{exp}}}(t)} + \frac{(Q^{\leftrightarrow}(t) - Q^{\leftrightarrow,\text{exp}}(t))^2}{\sigma^2_{Q^{\leftrightarrow,\text{exp}}}(t)} \right],$$

where the standard deviations are measured across trials/pairs of trials. Note that during this procedure, we keep the baseline constraints in equations (23) and update $\delta$ according to equation (24). The standard deviation tends to be very large for all three observables, stemming from short-timescale noise in the recorded traces. This still allows for a fit, but does not allow for the assessment of the goodness of fit. We will come back to this in Section 3.4.

#### 3.3 Fitting the peak heights

For simulations, we initialize the network in a random state and then let the dynamics evolve. We want to feed in stimuli that reproduce the experimental peak values of $R$ and the two $Q$s.

Peak values are indicated with a subscript $\circ_{\text{max}}$. $R^{\text{exp}}_{\text{max}}$ is given by the population mean at $t_{\text{max}}$

$$R^{\text{exp}}_{\text{max}} = \langle\langle x_{\alpha,i}(t_{\text{max}})\rangle_i\rangle_\alpha$$

and the overlaps by

$$Q^{\circlearrowleft,\text{exp}}_{\text{max}}(t_{\text{max}}) = \langle\langle x_{\alpha,i}(t_{\text{max}}) x_{\beta,i}(t_{\text{max}})\rangle_i\rangle_{(\alpha\beta)|C(\alpha)=C(\beta)},$$
$$Q^{\leftrightarrow,\text{exp}}_{\text{max}}(t_{\text{max}}) = \langle\langle x_{\alpha,i}(t_{\text{max}}) x_{\beta,i}(t_{\text{max}})\rangle_i\rangle_{(\alpha\beta)|C(\alpha)\neq C(\beta)},$$

where $Q^{\circlearrowleft,\text{exp}}_{\text{max}} \geq Q^{\leftrightarrow,\text{exp}}_{\text{max}}$ because stimuli of the same class are more similar to each other than stimuli of different classes.

The stimuli change the parameter constellations $c_1$ and $c_2$, on which the right-hand-side of the mean-field equations depend:

$$c_1 \to c_{1,\text{max}} = \frac{\bar{g}R + \delta + \eta}{\sqrt{R + \gamma + \Theta}}$$

and

$$c_2 \to \begin{cases} c^{\circlearrowleft}_{2,\text{max}} = \frac{Q+\gamma+\nu^{\circlearrowleft}\Theta}{R+\gamma+\Theta} \\ c^{\leftrightarrow}_{2,\text{max}} = \frac{Q+\gamma+\nu^{\leftrightarrow}\Theta}{R+\gamma+\Theta} \end{cases}.$$

For $c_2$ we have to distinguish between the intra-overlap and inter-overlap case. Using the bisection-algorithm in Figure 2 and an initial guess for $\nu^{\leftrightarrow}$ we can first fit $\eta, \Theta$ to match $R^{\text{exp}}_{\text{max}}, Q^{\leftrightarrow,\text{exp}}_{\text{max}}$.

As an initial guess, $\nu^{\leftrightarrow}$ can be chosen $\in [-1, 1]$ as long as one solution exists. It has to be chosen larger if the inter-overlap in relation to the population mean is large. The missing parameter $\nu^{\circlearrowleft}$ is found by bisectioning in $\nu^{\circlearrowleft} \in [-1, 1]$. This enables us to find a set $\eta, \Theta, \nu^{\leftrightarrow}, \nu^{\circlearrowleft}$ that reproduces the experimental peak heights from which the decay starts. Keeping the parameter constellations constant, we can always set $\nu^{\circlearrowleft} = 1$ for the parameters from the experiment. It is, however, beneficial to first treat it as a free parameter for an easier bisectioning, because it potentially could be chosen such that there is no solution for $\Theta$.

#### 3.4 Evaluating the goodness of fit

We want to have a measure for the quality of the fit. We do a smoothing window approach with a window size of 3 bins, i.e. $t_{\text{window}} = 3 \cdot 6\,\text{ms} = 18\,\text{ms}$. For the experimental observables (expm) and the model fit (model_fit) of $R, Q^{\circlearrowleft}, Q^{\leftrightarrow}$ we calculate the sliding window average

**def** fit_peak_heights_with_$\eta$_$\Theta$($R_{\max}^{\exp}$, $Q_{\max}^{\exp,\leftrightarrow}$, $\Omega_b$, $\nu^{\leftrightarrow}$):

    $c_{1,\max} = \sqrt{2}\,\mathrm{erf}^{-1}(2\,R_{\max}^{\exp} - 1)$

    **def** root_corr($\Theta$):

        $\eta = c_{1,\max}\sqrt{R_{\max}^{\exp} + \gamma + \Theta} - \delta - \bar{g}\,R_{\max}^{\exp}$

        $z = R_{\max}(\Omega_b, \Theta, \eta)$

        **return** $Q_{\max}^{\leftrightarrow}(\Omega_b, R = z, \Theta, \eta, \nu^{\leftrightarrow}) - Q_{\max}^{\leftrightarrow,\exp}$

    $\Theta$ = bisect(root_corr, $\Theta_{\mathrm{begin}}$, $\Theta_{\mathrm{end}}$)

    $\eta = c_{1,\max}\sqrt{R_{\max}^{\exp} + \gamma + \Theta} - \delta - \bar{g}\,R_{\max}^{\exp}$

    **return** $\eta$, $\Theta$

Figure 2: **Bisection algorithm determining parameters that fit experimental peak values.** $\Omega_b = \{\bar{g}, \delta, \gamma\}$ is a shorthand notation for the baseline parameters. $\nu^{\leftrightarrow}$ can be chosen as desired as long as there is a solution. Start and end values of $\Theta$ can be chosen to lie within $[0, \infty]$.

$$\mathrm{ex\bar{p}m}(t) = \langle \mathrm{expm}(t') \rangle_{t'=t\pm t_{\mathrm{window}}}$$

and variance

$$\begin{aligned}\mathrm{Var}(t) &= \left\langle \left(\mathrm{expm}(t') - \mathrm{ex\bar{p}m}(t')\right)^2 \right\rangle_{t'\in t\pm t_{\mathrm{window}}} \\ &= \langle\!\langle (\mathrm{expm}(t'))^2 \rangle\!\rangle_{t'\in t\pm t_{\mathrm{window}}}.\end{aligned}$$

The goodness of fit is then evaluated using the coefficient of determination

$$\mathrm{R}^2 = 1 - \left\langle \frac{(\mathrm{ex\bar{p}m}(t) - \mathrm{model\_fit}(t))^2}{\mathrm{Var}(t)} \right\rangle_t .$$

Here $\mathrm{R}^2$ is a standard measure that can become negative. This leaves us with three coefficients of determination, evaluating the goodness of fit for each of the three observables separately. Figure 3 and Figure 4 show that the deviation of the model to the data lies well within the error margins.

#### 3.5 Tuning the decay of the population mean

We want to investigate the role of the decay of the population mean $R$. For tuning its decay, we calculate the derivative of $\mathcal{T}(R, R)$, which determines the decay to linear order:

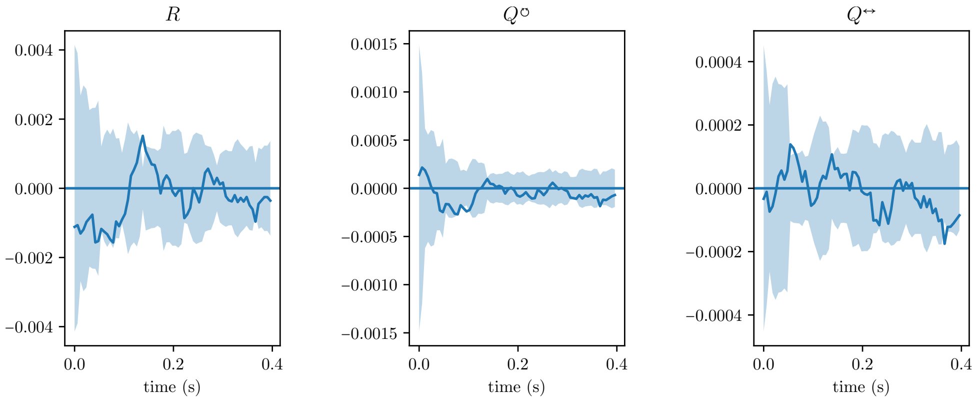

Figure 3: **Fitting for mouse 1.** Three plots for the three observables $R, Q^{\circlearrowleft}, Q^{\leftrightarrow}$. Opaque blue curve indicates the difference of the model to the smoothed data $\bar{\exp}\text{m}(t) - \text{model\_fit}(t)$. Shaded areas indicate the standard deviation $\pm\sqrt{\text{Var}(t)}$. Window size: $t_{\text{window}} = 18\,\text{ms}$.

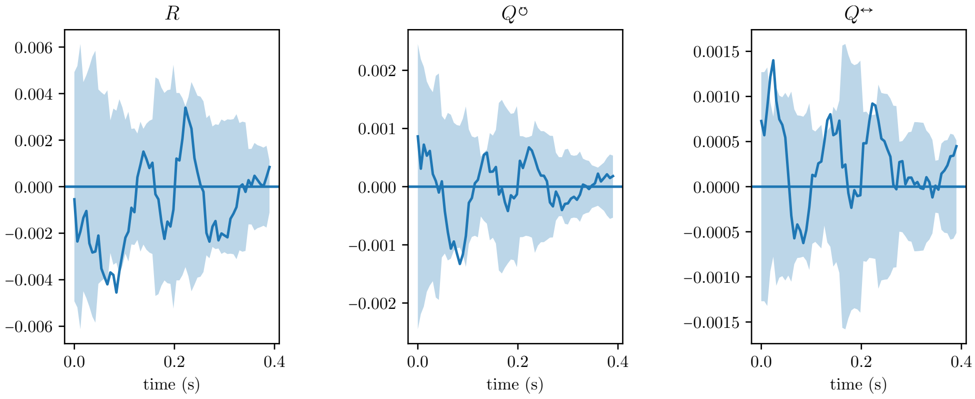

Figure 4: **Fitting for mouse 2.** Same as in Figure 3.

$$
\begin{aligned}
&\frac{d}{dR}\mathcal{T}\left(R,R\right)\\
&\overset{(20)}{=}\frac{d}{dR}\frac{1}{2}\left(1+\mathrm{erf}\left(\frac{\bar{g}R+\delta}{\sqrt{2}\sqrt{R+\gamma}}\right)\right)\\
&=\frac{d}{dR}\frac{1}{2}\left(1+\mathrm{erf}\left(\frac{c_1}{\sqrt{2}}\right)\right)\\
&=\frac{1}{\sqrt{2\pi}}e^{-c_1^2/2}c_1\left(\frac{\bar{g}}{\bar{g}R+\delta}-\frac{1}{2}\frac{1}{R+\gamma}\right).
\end{aligned}
$$

We can vary $\delta$ under the baseline constraint in equation (24) and set $\frac{d}{dR}\mathcal{T}\left(R,R\right)\big|_{R=R_0}$ to $1/{-1}$ to realize a slow/fast decay of the population mean. Importantly, this leaves the value of the peaks and the baselines invariant.

##### 3.6 Parameter table

In the main text, we use the considerations above to obtain $\bar{g}_{\mathrm{fast}} = -10.03$, $\delta_{\mathrm{fast}} = -0.11$ for the fast decay and $\bar{g}_{\mathrm{slow}} = -3.70$, $\delta_{\mathrm{slow}} = -0.24$ for the slow decay of the population mean.

| **Experimental observables** | left side | right side | |
|---|---|---|---|
| $R_{\mathrm{max}}$ | 6.63 | 12.05 | Initial population mean $[\mathrm{Hz}]$ |
| $R_0$ | 3.25 | 8.86 | Population mean baseline $[\mathrm{Hz}]$ |
| $Q^{\circlearrowleft}_{\mathrm{max}}$ | 3.08 | 2.53 | Initial variability within stimulus class $[\mathrm{Hz}^2]$ |
| $Q^{\leftrightarrow}_{\mathrm{max}}$ | 0.52 | 1.22 | Initial variability across stimulus classes $[\mathrm{Hz}^2]$ |
| $Q_0$ | 0.16 | 0.98 | Variability baseline $[\mathrm{Hz}^2]$ |
| $N$ | 141 | 65 | number of recorded neurons |

| **Inferred network parameters** | | | |
|---|---|---|---|
| $\tau$ | 29.29 | 25.65 | time-constant $[\mathrm{ms}]$ |
| $\bar{g}$ | −4.47 | −1.69 | average coupling strength (inhibition) |
| $\delta$ | −0.22 | −0.30 | average threshold |
| $\gamma$ | 0.0031 | 0.0062 | variability of thresholds |
| $T$ | 62 | 68 | stimulus switch-off $[\mathrm{ms}]$ |

| **Bayesian inference parameters** | | | |
|---|---|---|---|
| $N_{\mathrm{trials}}$ | 50 | 40 | |
| $N_{\mathrm{train}}$ | 35 | 28 | number of training samples/trials |
| $N_{\mathrm{test}}$ | 15 | 12 | number of test samples/trials |
| $\kappa$ | $0.1\,R_{\mathrm{max}}$ | $0.1\,R_{\mathrm{max}}$ | readout noise |
| $g_w/N$ | $1/N$ | $1/N$ | variance of prior |
| | 100 | 100 | number of seeds for sampling training, test data |

| **Simulation Parameters** | | |
|---|---|---|
| $N_{\mathrm{trials}}$ | 50 | |
| $\mathrm{d}t$ | 0.6 | Simulation step size $[\mathrm{ms}]$ |
| $N_{\mathrm{model}}$ | 35250 | Number of neurons |

Table 1: **Network parameters.** Left/right side refers to the side of stimulus application for the visual and tactile stimuli on the mouse. The experimental observable $R$ is averaged over all trials, and the overlaps $Q$ are averaged over pairs of trials. The overlap within classes $Q^{\circlearrowleft}$ is computed by averaging over all pairs of trials with the same stimulus class (both visual or both tactile).

### 4 Separability in the limit of many neurons

We quantify separability using a linear model $y(t) = \boldsymbol{w}^{\mathrm{T}}\boldsymbol{x}(t) + \epsilon$ with $\boldsymbol{w} \overset{\text{i.i.d.}}{\sim} \mathcal{N}\left(0, g_w^2/N\right)$ and $\epsilon \overset{\text{i.i.d.}}{\sim} \mathcal{N}\left(0, \sigma_n^2\right)$, where the readout $\boldsymbol{w} \in \mathbb{R}^N$ is identical across trials and the scalar readout noise $\epsilon \in \mathbb{R}$ is independent over trials. The vector $\boldsymbol{w}$ describes a linear downstream readout from the neural activity and $\epsilon$ describes input-independent variability in the neural response that cannot be used for inference, propagated through the readout through $\boldsymbol{w}$. After conditioning on training data $\boldsymbol{X} \in \mathbb{R}^{N\times 2N_{\mathrm{train}}}$, the predictive statistics for the test data $p(\boldsymbol{y}^*|\boldsymbol{X}^*, \boldsymbol{y}, \boldsymbol{X})$ follows from Bayesian inference for a Gaussian process [8]. This expression yields the Bayes-optimal predictor (i.e., with minimal mean-squared error on heldout test data). Equivalently, this predictor can be thought of as resulting from a regularized linear regression of $\boldsymbol{w}$ of the network responses $\boldsymbol{X}$ to labels $\boldsymbol{y}$.

The two cumulants of this Gaussian process can be calculated using its kernel matrix. The kernel directly follows as the dot-product between two neural state vectors

$$\begin{aligned} K(\boldsymbol{x}_\alpha, \boldsymbol{x}_\beta) &= \frac{g_w^2}{N}\boldsymbol{x}_\alpha^{\mathrm{T}}\boldsymbol{x}_\beta \\ &= \frac{g_w^2}{N}\sum_{i=1}^{N} x_{\alpha i} x_{\beta i}. \end{aligned}$$

For many neurons the entries of this kernel approach three distinct values, see Figure 5

$$\begin{aligned} K(\boldsymbol{x}_\alpha, \boldsymbol{x}_\alpha) &\to g_w^2 R \\ K(\boldsymbol{x}_\alpha, \boldsymbol{x}_\beta) &\to g_w^2 Q^{\circlearrowleft} \quad \text{for} \quad C(\alpha) = C(\beta) \\ K(\boldsymbol{x}_\alpha, \boldsymbol{x}_\beta) &\to g_w^2 Q^{\leftrightarrow} \quad \text{for} \quad C(\alpha) \neq C(\beta). \end{aligned}$$

#### 4.1 Predictive mean

We here derive analytical expressions for the mean of the predictive distribution. We start from the general formula for a Gaussian process [8]

$$\boldsymbol{\mu}(\boldsymbol{X}^*) = \boldsymbol{K}(\boldsymbol{X}^*, \boldsymbol{X})\left(\boldsymbol{K}(\boldsymbol{X}, \boldsymbol{X}) + \sigma_n^2 \mathbb{1}\right)^{-1}\boldsymbol{y} \tag{25}$$

with $\boldsymbol{K}(\boldsymbol{X}^*, \boldsymbol{X}) \in \mathbb{R}^{2N_{\mathrm{train}}\times 2N_{\mathrm{train}}}$.

In the mean-field setting the kernel is homogeneous in that it only has three different values which we call $a, b, c$. We can write the kernel as

$$K(\boldsymbol{x}_\alpha, \boldsymbol{x}_\beta) + \sigma_n^2\delta_{\alpha\beta} = \delta_{\alpha\beta} a + \left(\delta_{C(\alpha),C(\beta)} - \delta_{\alpha\beta}\right) b + \left(1 - \delta_{C(\alpha),C(\beta)}\right) c$$

We can visualize this kernel in matrix representation, also see Figure 5

$$\boldsymbol{K}(\boldsymbol{X}, \boldsymbol{X}) = \left(\begin{array}{cccc|cccc} a & b & \dots & b & c & \dots & \dots & c \\ b & \ddots & & \vdots & \vdots & \ddots & & \vdots \\ \vdots & & \ddots & b & \vdots & & \ddots & \vdots \\ b & \dots & b & a & c & \dots & \dots & c \\ \hline c & \dots & \dots & c & a & b & & b \\ \vdots & \ddots & & \vdots & b & \ddots & & \\ \vdots & & \ddots & \vdots & & & \ddots & b \\ c & \dots & \dots & c & b & & b & a \end{array}\right).$$

The other matrix we need for the predictive mean in equation (25) is

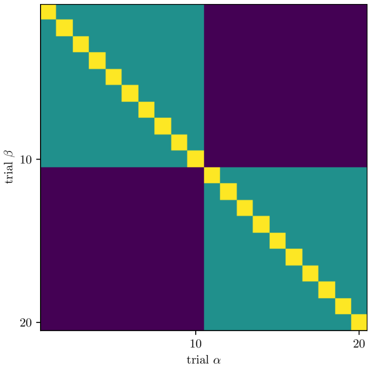

Figure 5: **Kernel in matrix representation.** The kernel matrix $\left[\boldsymbol{K}(\boldsymbol{X},\boldsymbol{X})\right]_{\alpha\beta}$ allows for the assessment of separability, see equation (25). Different colors correspond to the values $a$ (yellow), $b$ (green), $c$ (indigo). $N_{\text{train}} = 10$. The fluctuations around the three distinct entries of this kernel diminish for many neurons.

$$\boldsymbol{K}(\boldsymbol{x}_\alpha^*,\boldsymbol{x}_\beta) = \delta_{C(\alpha),C(\beta)} b + \left(1 - \delta_{C(\alpha),C(\beta)}\right) c.$$

The diagonal entry $a$ does not show up here since no test point is included in the training set $\boldsymbol{x}_\alpha^* \neq \boldsymbol{x}_\beta \;\forall\, \alpha,\beta$. Taking $\boldsymbol{y} = (-1,\dots,-1,1,\dots 1)^{\mathrm{T}} \in \mathbb{R}^{2N_{\text{train}}}$, i.e. the first half of the training points are from one stimulus class and the rest from the other, we are able to decompose

$$\boldsymbol{K}(\boldsymbol{X},\boldsymbol{X}) + \sigma_n^2 \mathbb{1} = l\mathbb{1} + p\,\mathbf{1}\mathbf{1}^{\mathrm{T}} + q\,\boldsymbol{y}\boldsymbol{y}^{\mathrm{T}}$$

into outer products of two mutually orthogonal vectors $\boldsymbol{y}$ and $\mathbf{1} = (1,\dots,1)^{\mathrm{T}} \in \mathbb{R}^{2N_{\text{train}}}$ with

$$\begin{aligned} b =& p + q\,, \\ c =& p - q\,, \\ a - b =& l\,. \end{aligned}$$

This enables us to carry out the matrix multiplication

$$\begin{aligned} \mu(\boldsymbol{x}^*) =& \boldsymbol{K}(\boldsymbol{x}^*,\boldsymbol{X})\left(\boldsymbol{K}(\boldsymbol{X},\boldsymbol{X}) + \sigma_n^2\mathbb{1}\right)^{-1}\boldsymbol{y} \\ =& \boldsymbol{K}(\boldsymbol{x}^*,\boldsymbol{X})\boldsymbol{y}\left[l + 2N_{\text{train}}q\right]^{-1} \\ =& C(\boldsymbol{x}^*)N_{\text{train}}(b-c)\left[l + 2N_{\text{train}}q\right]^{-1} \\ =& C(\boldsymbol{x}^*)N_{\text{train}}(b-c)\left[a - b + N_{\text{train}}(b-c)\right]^{-1} \\ =& C(\boldsymbol{x}^*)\frac{1}{1 + \frac{1}{N_{\text{train}}}\frac{a-b}{b-c}} \\ =:& C(\boldsymbol{x}^*)\mu, \end{aligned} \tag{26}$$

where $C(\boldsymbol{x}^*) \in \{-1,1\}$ is the true class label of the test point. We are particularly interested in the absolute value, which we have denoted as the separability $\mu$. Inserting $a = g_w^2 R + \sigma_n^2$, $b = g_w^2 Q^{\circlearrowleft}$, and $c = g_w^2 Q^{\leftrightarrow}$ we can write

$$
\begin{aligned}
\mu &= \left(1+\frac{1}{N_{\mathrm{train}}}\frac{a-b}{b-c}\right)^{-1} \\
&= \left(1+\frac{1}{N_{\mathrm{train}}}\frac{\kappa+R-Q^{\circlearrowleft}}{Q^{\circlearrowleft}-Q^{\leftrightarrow}}\right)^{-1}
\end{aligned} \tag{27}
$$

with the noise strength $\kappa = \sigma_n^2/g_w^2$. Note that this formula still holds if we were to include a bias term in the linear model because a global additive term on the kernel does not affect the separability in equation (27).

##### 4.1.1 Time evolution

For the dynamical quantities $R(t), Q^{\circlearrowleft}(t), Q^{\leftrightarrow}(t)$ we can calculate the time derivative of equation (27)

$$
\begin{aligned}
\tau\frac{d}{dt}\mu &= \left(1+\frac{a-b}{N_{\mathrm{train}}\,(b-c)}\right)^{-2}\tau\frac{d}{dt}\frac{a-b}{N_{\mathrm{train}}\,(b-c)} \\
&= -\mu^2\left[\frac{\frac{d}{dt}\,(a-b)}{N_{\mathrm{train}}\,(b-c)}-\frac{(a-b)}{N_{\mathrm{train}}\,(b-c)^2}\frac{d}{dt}\,(b-c)\right] \\
&= -\mu^2(\mu^{-1}-1)\left[\frac{\tau\frac{d}{dt}(a-b)}{a-b}-\frac{\tau\frac{d}{dt}(b-c)}{b-c}\right] \\
&= -\mu(1-\mu)\left[\frac{\tau\frac{d}{dt}(a-b)}{a-b}-\frac{\tau\frac{d}{dt}(b-c)}{b-c}\right] \\
&= -\mu(1-\mu)\left[\frac{\tau\frac{d}{dt}(\kappa+R-Q^{\circlearrowleft})}{\kappa+R-Q^{\circlearrowleft}}-\frac{\tau\frac{d}{dt}(Q^{\circlearrowleft}-Q^{\leftrightarrow})}{Q^{\circlearrowleft}-Q^{\leftrightarrow}}\right] \\
&= -\mu(1-\mu)\left[\frac{\kappa+\mathcal{T}(R,R)-\mathcal{T}(R,Q^{\circlearrowleft})}{\kappa+R-Q^{\circlearrowleft}}-\frac{\mathcal{T}(R,Q^{\circlearrowleft})-\mathcal{T}(R,Q^{\leftrightarrow})}{Q^{\circlearrowleft}-Q^{\leftrightarrow}}\right] \\
&\quad -\mu(1-\mu)\left[s_1-s_2\right],
\end{aligned} \tag{28}
$$

where we used equation (19) and defined the secants $s_1$, $s_2$ as

$$
\begin{aligned}
s_1 &= \frac{\kappa+\mathcal{T}(R,R)-\mathcal{T}(R,Q^{\circlearrowleft})}{\kappa+R-Q^{\circlearrowleft}} \\
s_2 &= \frac{\mathcal{T}(R,Q^{\circlearrowleft})-\mathcal{T}(R,Q^{\leftrightarrow})}{Q^{\circlearrowleft}-Q^{\leftrightarrow}}.
\end{aligned}
$$

This equation is not fully expressible solely in terms of $\mu$. This implies that the time course of separability is not fully determined by separability at an initial time – rather, the values of the underlying network observables $R, Q^{\circlearrowleft}, Q^{\leftrightarrow}$ continue to influence separability. For the time evolution of separability it is of particular importance how the differences $R-Q^{\circlearrowleft}$ and $Q^{\circlearrowleft}-Q^{\leftrightarrow}$ evolve in time. The two secants $s_1$, $s_2$ in equation (28) measure the strength of the correlation transmission function relative to the leak terms of $R-Q^{\circlearrowleft}$ and $Q^{\circlearrowleft}-Q^{\leftrightarrow}$. This can be seen by rewriting the mean-field equations (19) as

$$
\begin{aligned}
\tau\frac{d}{dt}\ln\left(\kappa+R-Q^{\circlearrowleft}\right) &= -1+s_1\,, \\
\tau\frac{d}{dt}\ln\left(Q^{\circlearrowleft}-Q^{\leftrightarrow}\right) &= -1+s_2\,.
\end{aligned} \tag{29}
$$

#### Geometric interpretation of the decay of separability

We can also understand the decay of the separability geometrically. The three observables $R$, $Q^{\circlearrowleft}$, and $Q^{\leftrightarrow}$ explicitly influence the separability via the two secants of the correlation transmission function $\mathcal{T}(R, \circ)$

$$
\begin{aligned}
s_1 &= \frac{\kappa + \mathcal{T}(R,R) - \mathcal{T}(R,Q^{\circlearrowleft})}{\kappa + R - Q^{\circlearrowleft}}\,,\\
s_2 &= \frac{\mathcal{T}(R,Q^{\circlearrowleft}) - \mathcal{T}(R,Q^{\leftrightarrow})}{Q^{\circlearrowleft} - Q^{\leftrightarrow}}\,.
\end{aligned}
$$

The secant $s_1$ measures the relative change of the total variability $\kappa + R - Q^{\circlearrowleft}$ and the secant $s_2$ measures the relative change of the signal $Q^{\circlearrowleft} - Q^{\leftrightarrow}$, both of which determine the separability through equation (27). The secant $s_1$ can be understood geometrically by connecting the points $\{Q^{\circlearrowleft}, \mathcal{T}(R,Q^{\circlearrowleft})\}$ and $\{R+\kappa, \mathcal{T}(R,R)+\kappa\}$ (for $\kappa = 0$). Likewise, we obtain the secant $s_2$ by connecting the points $\{Q^{\leftrightarrow}, \mathcal{T}(R,Q^{\leftrightarrow})\}$ and $\{Q^{\circlearrowleft}, \mathcal{T}(R,Q^{\circlearrowleft})\}$, see Figure 6. The difference between these two secants, by equation (28), modulates the speed by which $\mu$ evolves.

Figure 6$\mathbf{a}_1, \mathbf{a}_2$ explains why the separability decays similarly for the two cases shown in Figure 4 of the main text: the difference between the time evolution of the total variability minus the signal, $s_1 - s_2$, has a similar value and does not change significantly for $t \geq t_1$ (blue dot).

#### Condition for transient increase in separability

For a transient increase of the separability $\mu$, the right side of equation (28) has to become positive, i.e. $s_1 < s_2$. This means that the relative change of $Q^{\circlearrowleft} - Q^{\leftrightarrow}$ has to be larger than that of $R - Q^{\circlearrowleft}$.

For zero noise $\kappa = 0$, $s_1 > s_2$ because the derivative of the correlation transmission function ${}^{d}\!/_{dQ}\mathcal{T}(R,Q)$ is positive. With

$$
\begin{aligned}
&\frac{d}{dc}\big\langle H(h^{(1)})H(h^{(2)})\big\rangle_{\begin{pmatrix}h^{(1)}\\h^{(2)}\end{pmatrix}\sim\mathcal{N}\left(\begin{pmatrix}r\\r\end{pmatrix},q\begin{pmatrix}1&c\\c&1\end{pmatrix}\right)}\\
&\overset{\substack{\text{Price's}\\\text{Theorem}}}{=} q\big\langle \delta(h^{(1)})\delta(h^{(2)})\big\rangle_{\begin{pmatrix}h^{(1)}\\h^{(2)}\end{pmatrix}\sim\mathcal{N}\left(\begin{pmatrix}r\\r\end{pmatrix},q\begin{pmatrix}1&c\\c&1\end{pmatrix}\right)}\\
&= q\frac{1}{2\pi\sqrt{\det\left(q\begin{pmatrix}1&c\\c&1\end{pmatrix}\right)}}\exp\left(-\frac{1}{2}\begin{pmatrix}r\\r\end{pmatrix}^{\mathrm{T}} q^{-1}\begin{pmatrix}1&c\\c&1\end{pmatrix}^{-1}\begin{pmatrix}r\\r\end{pmatrix}\right)\\
&= q\frac{1}{2\pi\sqrt{q^2\left(1-c^2\right)}}\exp\left(-\frac{1}{2}\begin{pmatrix}r\\r\end{pmatrix}^{\mathrm{T}} q^{-1}\begin{pmatrix}1&c\\c&1\end{pmatrix}^{-1}\begin{pmatrix}r\\r\end{pmatrix}\right)\\
&= \frac{1}{2\pi\sqrt{1-c^2}}\exp\left(-\frac{1}{2}\frac{1}{q}\frac{1}{(1-c^2)}\begin{pmatrix}r\\r\end{pmatrix}^{\mathrm{T}}\begin{pmatrix}(1-c)r\\(1-c)r\end{pmatrix}\right)\\
&= \frac{1}{2\pi\sqrt{1-c^2}}\exp\left(-\frac{1}{q}\frac{1-c}{1-c^2}r^2\right)\\
&= \frac{1}{2\pi\sqrt{1-c^2}}\exp\left(-\frac{1}{q}\frac{r^2}{1+c}\right)
\end{aligned}
$$

we can write

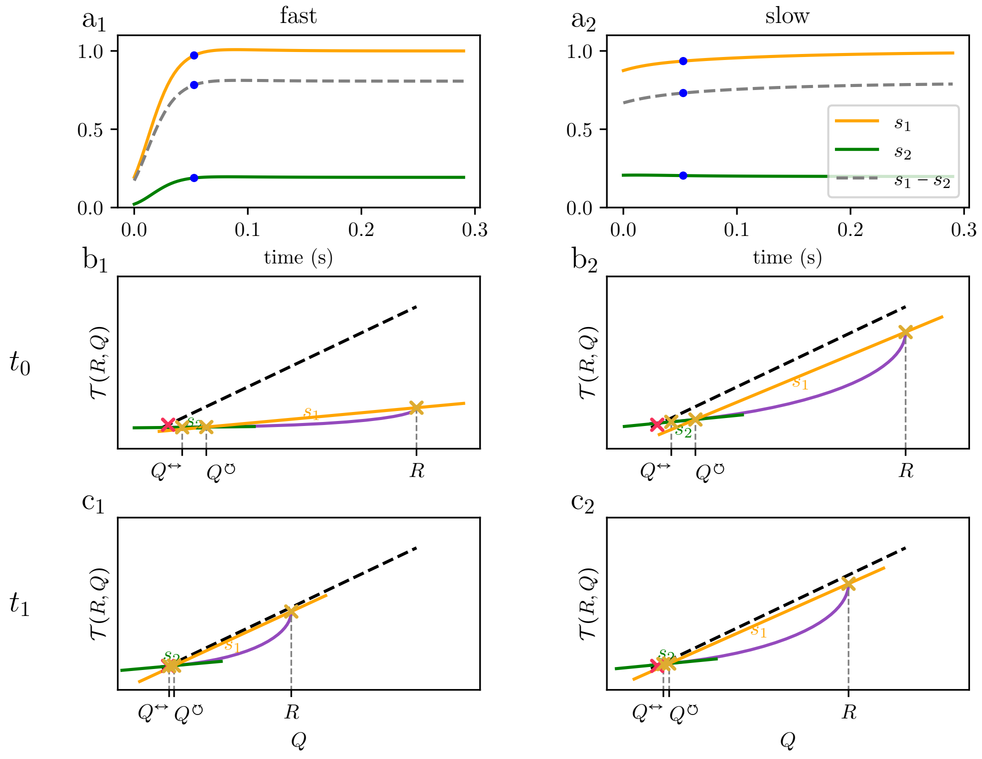

Figure 6: Understanding the time evolution of separability in a geometric picture through the total variability and the signal: $\mathbf{a}_1, \mathbf{a}_2$: Time evolution of $s_1$ and $s_2$, which measure the change of total variability and the signal, respectively. Additionally, the difference $s_1 - s_2$ is plotted (dashed line), which shapes the time evolution of the separability in equation (28). $\mathbf{b}_1, \mathbf{b}_2, \mathbf{c}_1, \mathbf{c}_2$: Illustration of equation (28): correlation transmission function $\mathcal{T}(R, \circ)$ (violet curve). The decay depends on the form of the correlation transmission function $\mathcal{T}(R, Q)$ along the time course of $R$ and $Q$. Intersections of $\mathcal{T}$ with the identity line (black dashed) determine fixed points (red crosses) for the overlaps $Q^{\circlearrowleft}, Q^{\leftrightarrow}$, cf. equation (19), which correspond to the baseline value of both $Q^{\circlearrowleft}$ and $Q^{\leftrightarrow}$ for long times. The first secant $s_1$ (orange line) connects $R$ and $Q^{\circlearrowleft}$, the second secant $s_2$ (green line) connects $Q^{\circlearrowleft}$ and $Q^{\leftrightarrow}$. $\mathbf{b}_1$ Initial state $t_0 = 0$ for the quickly decaying population mean in Figure 4 of the main text. $\mathbf{b}_2$ Initial state $t_0 = 0$ for the slowly decaying population mean. $\mathbf{c}_1, \mathbf{c}_2$: Secant picture for quickly/slowly decaying population mean at $t_1 = 3.69\,\tau = 0.052\,\mathrm{s}$ (blue dot). The example here is for zero noise $\kappa = 0$. The addition of noise $\kappa \neq 0$ in (28) corresponds to moving the marker at $R$ to larger values and simultaneously decreasing its distance to the dashed unit line ($s_1 \xrightarrow{\kappa\to\infty} 1$). The distance to the unit line represents the velocity of the decay according to (19).

$$
\begin{aligned}
&\frac{d}{dQ}\mathcal{T}(R,Q)\\
=&q\frac{d}{dc}\left\langle H(h^{(1)})H(h^{(2)})\right\rangle_{\begin{pmatrix}h^{(1)}\\h^{(2)}\end{pmatrix}\sim\mathcal{N}\left(\begin{pmatrix}r\\r\end{pmatrix},q\begin{pmatrix}1&c\\c&1\end{pmatrix}\right)}\\
=&\frac{q}{2\pi\sqrt{1-c^2}}\exp\left(-\frac{1}{q}\frac{r^2}{1+c}\right)\Bigg|_{\substack{r=\bar{g}R+\delta\\ q=R+\gamma\\ c=Q+\gamma/R+\gamma}}>0
\end{aligned}
$$

and observe that this is always a positive quantity.

Because $s_1 > s_2$ for $\kappa = 0$, a transient increase in separability can only occur when observations are noisy $\kappa \neq 0$. The introduction of noise blurs the relative change of $\kappa + R - Q^{\circlearrowleft}$

$$
\frac{\tau\frac{d}{dt}\left(\kappa+R-Q^{\circlearrowleft}\right)}{\kappa+R-Q^{\circlearrowleft}}\xrightarrow{\kappa\to\infty}0,
$$

enabling $s_1 < s_2$ when the relative change of $Q^{\circlearrowleft} - Q^{\leftrightarrow}$ is positive (compare to equation (29))

$$
\frac{\tau\frac{d}{dt}(Q^{\circlearrowleft}-Q^{\leftrightarrow})}{Q^{\circlearrowleft}-Q^{\leftrightarrow}}>0.
$$

### 5 Information transmission

#### 5.1 Bayesian inference of the stimulus label

We take $P$ linear readouts $y_p = \boldsymbol{w}_p^{\mathrm{T}}\boldsymbol{x} + \epsilon$ with $\boldsymbol{y}_p \in \{0,1\}^P$ and one weight vector $\boldsymbol{w}_p$ for every stimulus class $p = 1,\ldots,P$. The $p$-th readout is conditioned on the $p$-th stimulus using Bayes formula. To account for the different linear readouts, we introduce the weight matrix $\boldsymbol{W} = (\boldsymbol{w}_1,\ldots,\boldsymbol{w}_P) \in \mathbb{R}^{N_{\mathrm{nrn}}\times P}$, the data matrix with $N_{\mathrm{train}}$ vectors for every class $P$, $\boldsymbol{X} = (\boldsymbol{x}_1,\ldots,\boldsymbol{x}_{N_{\mathrm{train}}P}) \in \mathbb{R}^{N_{\mathrm{nrn}}\times N_{\mathrm{train}}P}$, and the associated label matrix $\boldsymbol{Y} = (\boldsymbol{y}_1,\ldots,\boldsymbol{y}_P) \in \mathbb{R}^{N_{\mathrm{train}}P\times P}$. The labels are one-hot encoded $(\boldsymbol{y}_p)_d = \begin{cases}1, & N_{\mathrm{train}}\cdot(p-1)\le d < N_{\mathrm{train}}\cdot p\\ 0, & \text{else}\end{cases}$.

For multivariate linear regression, the likelihood factorizes [9, Chapter 11, (11.2)]

$$
\begin{aligned}
p(\boldsymbol{Y}|\boldsymbol{X},\boldsymbol{W}) &= \prod_{p=1}^{P} p(\boldsymbol{y}_p|\boldsymbol{X},\boldsymbol{w}_p)\\
p(\boldsymbol{y}_p|\boldsymbol{X},\boldsymbol{w}_p) &= \mathcal{N}(\boldsymbol{X}^{\mathrm{T}}\boldsymbol{w}_p,\sigma_n^2).
\end{aligned}
\tag{30}
$$

We likewise choose independent weights for each $p$

$$
w_{pi} \overset{\text{i.i.d.}}{\sim} \mathcal{N}(0, g_w^2/N)\,,
$$

where $g_w^2$ has to be $\mathcal{O}(1)$ for the prior variance not to scale with the number of neurons. For this choice, the joint distribution $p(\boldsymbol{Y}|\boldsymbol{X},\boldsymbol{W})p(\boldsymbol{W})$ also factorizes over channels, and consequentially the posterior as well

$$
p(\boldsymbol{y}^*|\boldsymbol{X}^*,\boldsymbol{y},\boldsymbol{X}) = \prod_{p=1}^{P} p(\boldsymbol{y}_p^*|\boldsymbol{X}^*,\boldsymbol{y}_p,\boldsymbol{X}), \tag{31}
$$

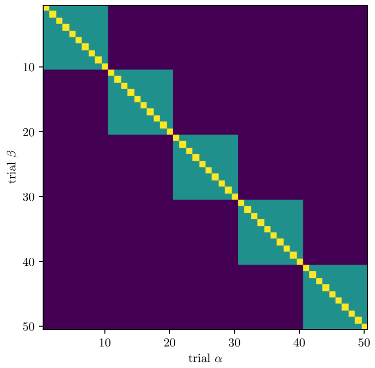

Figure 7: **Kernel in matrix representation.** We can assess the separability of $P$ stimuli using this kernel $[\boldsymbol{K}(\boldsymbol{X},\boldsymbol{X})]_{\alpha\beta}$. Different colors correspond to the values $a$ (yellow), $b$ (green), $c$ (indigo). $N_{\text{train}} = 10$ and $P = 5$. The fluctuations around the three distinct entries of this kernel diminish in the limit of large numbers of neurons.

which in particular makes the posterior covariance diagonal between readouts. We thus obtain the predictor statistics

$$\begin{aligned}\boldsymbol{\mu}_p(\boldsymbol{X}^*) =& \boldsymbol{K}(\boldsymbol{X}^*,\boldsymbol{X})\left(\boldsymbol{K}(\boldsymbol{X},\boldsymbol{X})+\sigma_n^2\mathbb{1}\right)^{-1}\boldsymbol{y}_p\,,\\ \boldsymbol{\Sigma}(\boldsymbol{X}^*) =& \boldsymbol{K}(\boldsymbol{X}^*,\boldsymbol{X}^*)+\sigma_n^2\mathbb{1} \quad - \quad \boldsymbol{K}(\boldsymbol{X}^*,\boldsymbol{X})\left(\boldsymbol{K}(\boldsymbol{X},\boldsymbol{X})+\sigma_n^2\mathbb{1}\right)^{-1}\boldsymbol{K}(\boldsymbol{X},\boldsymbol{X}^*).\end{aligned} \tag{32}$$

Note that the posterior covariance does not depend on the training labels and thus does not carry an index $p$.

##### 5.1.1 Calculating the mean

We again consider the limit of many neurons, in which the kernel matrix only shows three distinct entries. Compared to equation (26), we here have a total number of classes $P$, hence the homogeneous kernel has $P$ blocks, see Figure 7.

We here call the three distinct values of the inverse kernel matrix $p, q, r$.

$$
\boldsymbol{K}(\boldsymbol{X},\boldsymbol{X})\;(\boldsymbol{K}(\boldsymbol{X},\boldsymbol{X}))^{-1} = \underbrace{\left(\begin{array}{cccc|cccc|cccc|cccc}
a & b & \dots & b & c & \dots & \dots & c & & & & & c & \dots & \dots & c \\
b & \ddots & & \vdots & \vdots & \ddots & & \vdots & & \dots & \dots & & \vdots & \ddots & & \vdots \\
\vdots & & \ddots & b & \vdots & & \ddots & \vdots & & \dots & \dots & & \vdots & & \ddots & \vdots \\
b & \dots & b & a & c & \dots & \dots & c & & & & & c & \dots & \dots & c \\
\hline
c & \dots & \dots & c & & & & & & & & & & & & \\
\vdots & \ddots & & \vdots & & \ddots & \ddots & & & \ddots & \ddots & & & \vdots & \vdots & \\
\vdots & & \ddots & \vdots & & \ddots & \ddots & & & \ddots & \ddots & & & \vdots & \vdots & \\
c & \dots & \dots & c & & & & & & & & & & & & \\
\hline
& & & & & & & & & & & & c & \dots & \dots & c \\
& \vdots & \vdots & & & \ddots & \ddots & & & \ddots & \ddots & & \vdots & \ddots & & \vdots \\
& \vdots & \vdots & & & \ddots & \ddots & & & \ddots & \ddots & & \vdots & & \ddots & \vdots \\
& & & & & & & & & & & & c & \dots & \dots & c \\
\hline
c & \dots & \dots & c & & & & & c & \dots & \dots & c & a & b & \dots & b \\
\vdots & \ddots & & \vdots & & \dots & \dots & & \vdots & \ddots & & \vdots & b & \ddots & & \vdots \\
\vdots & & \ddots & \vdots & & \dots & \dots & & \vdots & & \ddots & \vdots & \vdots & & \ddots & b \\
c & \dots & \dots & c & & & & & c & \dots & \dots & c & b & \dots & b & a
\end{array}\right)}_{P \text{ blocks of width } N_{\mathrm{train}}}
$$

$$
\times \left(\begin{array}{cccc|cccc|cccc|cccc}
p & q & \dots & q & r & \dots & \dots & r & & & & & r & \dots & \dots & r \\
q & \ddots & & \vdots & \vdots & \ddots & & \vdots & & \dots & \dots & & \vdots & \ddots & & \vdots \\
\vdots & & \ddots & q & \vdots & & \ddots & \vdots & & \dots & \dots & & \vdots & & \ddots & \vdots \\
q & \dots & q & p & r & \dots & \dots & r & & & & & r & \dots & \dots & r \\
\hline
r & \dots & \dots & r & & & & & & & & & & & & \\
\vdots & \ddots & & \vdots & & \ddots & \ddots & & & \ddots & \ddots & & & \vdots & \vdots & \\
\vdots & & \ddots & \vdots & & \ddots & \ddots & & & \ddots & \ddots & & & \vdots & \vdots & \\
r & \dots & \dots & r & & & & & & & & & & & & \\
\hline
& & & & & & & & & & & & r & \dots & \dots & r \\
& \vdots & \vdots & & & \ddots & \ddots & & & \ddots & \ddots & & \vdots & \ddots & & \vdots \\
& \vdots & \vdots & & & \ddots & \ddots & & & \ddots & \ddots & & \vdots & & \ddots & \vdots \\
& & & & & & & & & & & & r & \dots & \dots & r \\
\hline
r & \dots & \dots & r & & & & & r & \dots & \dots & r & p & q & \dots & q \\
\vdots & \ddots & & \vdots & & \dots & \dots & & \vdots & \ddots & & \vdots & q & \ddots & & \vdots \\
\vdots & & \ddots & \vdots & & \dots & \dots & & \vdots & & \ddots & \vdots & \vdots & & \ddots & q \\
r & \dots & \dots & r & & & & & r & \dots & \dots & r & q & \dots & q & p
\end{array}\right) \stackrel{!}{=} 1\,.
$$

We obtain three equations from this inversion formula

$$
\begin{aligned}
ap + (N_{\mathrm{train}} - 1)bq + N_{\mathrm{train}}cr + N_{\mathrm{train}}(P-2)cr &= 1 \qquad \text{(I)} \\
aq + bp + (N_{\mathrm{train}} - 2)bq + N_{\mathrm{train}}cr + N_{\mathrm{train}}(P-2)cr &= 0 \qquad \text{(II)} \\
ar + (N_{\mathrm{train}} - 1)br + cp + (N_{\mathrm{train}} - 1)cq + N_{\mathrm{train}}(P-2)cr &= 0 \qquad \text{(III)}
\end{aligned} \tag{33}
$$

As we will see later, we only have to consider differences of means of readouts, not the individual means. The difference between the mean predictors of stimulus 1 and 2 is

$$
\begin{aligned}
\boldsymbol{\mu}_1(\boldsymbol{X}^*) - \boldsymbol{\mu}_2(\boldsymbol{X}^*) &= \boldsymbol{K}(\boldsymbol{X}^*, \boldsymbol{X}) \left(\boldsymbol{K}(\boldsymbol{X}, \boldsymbol{X})\right)^{-1} (\boldsymbol{y}_1 - \boldsymbol{y}_2) \\
&= \left(\begin{array}{c|c|c|c}
\begin{matrix} b & & \cdots & b \\ & \ddots & & \vdots \\ \vdots & & \ddots & \\ b & \cdots & & b \end{matrix} &
\begin{matrix} c & \cdots & \cdots & c \\ \vdots & \ddots & & \vdots \\ \vdots & & \ddots & \vdots \\ c & \cdots & \cdots & c \end{matrix} &
\begin{matrix} \cdots & \cdots \\ \cdots & \cdots \end{matrix} &
\begin{matrix} c & \cdots & \cdots & c \\ \vdots & \ddots & & \vdots \\ \vdots & & \ddots & \vdots \\ c & \cdots & \cdots & c \end{matrix} \\
\hline
\begin{matrix} c & \cdots & \cdots & c \\ \vdots & \ddots & & \vdots \\ \vdots & & \ddots & \vdots \\ c & \cdots & \cdots & c \end{matrix} &
\begin{matrix} \ddots & \ddots \\ \ddots & \ddots \end{matrix} &
\begin{matrix} \ddots & \ddots \\ \ddots & \ddots \end{matrix} &
\begin{matrix} \vdots & \vdots \\ \vdots & \vdots \end{matrix} \\
\hline
\begin{matrix} \vdots & \vdots \\ \vdots & \vdots \end{matrix} &
\begin{matrix} \ddots & \ddots \\ \ddots & \ddots \end{matrix} &
\begin{matrix} \ddots & \ddots \\ \ddots & \ddots \end{matrix} &
\begin{matrix} c & \cdots & \cdots & c \\ \vdots & \ddots & & \vdots \\ \vdots & & \ddots & \vdots \\ c & \cdots & \cdots & c \end{matrix} \\
\hline
\begin{matrix} c & \cdots & \cdots & c \\ \vdots & \ddots & & \vdots \\ \vdots & & \ddots & \vdots \\ c & \cdots & \cdots & c \end{matrix} &
\begin{matrix} \cdots & \cdots \\ \cdots & \cdots \end{matrix} &
\begin{matrix} c & \cdots & \cdots & c \\ \vdots & \ddots & & \vdots \\ \vdots & & \ddots & \vdots \\ c & \cdots & \cdots & c \end{matrix} &
\begin{matrix} b & & \cdots & b \\ & \ddots & & \vdots \\ \vdots & & \ddots & \\ b & \cdots & & b \end{matrix}
\end{array}\right) \\
&\times \left(\begin{array}{c|c|c|c}
\begin{matrix} p & q & \cdots & q \\ q & \ddots & & \vdots \\ \vdots & & \ddots & q \\ q & \cdots & q & p \end{matrix} &
\begin{matrix} r & \cdots & \cdots & r \\ \vdots & \ddots & & \vdots \\ \vdots & & \ddots & \vdots \\ r & \cdots & \cdots & r \end{matrix} &
\begin{matrix} \cdots & \cdots \\ \cdots & \cdots \end{matrix} &
\begin{matrix} r & \cdots & \cdots & r \\ \vdots & \ddots & & \vdots \\ \vdots & & \ddots & \vdots \\ r & \cdots & \cdots & r \end{matrix} \\
\hline
\begin{matrix} r & \cdots & \cdots & r \\ \vdots & \ddots & & \vdots \\ \vdots & & \ddots & \vdots \\ r & \cdots & \cdots & r \end{matrix} &
\begin{matrix} \ddots & \ddots \\ \ddots & \ddots \end{matrix} &
\begin{matrix} \ddots & \ddots \\ \ddots & \ddots \end{matrix} &
\begin{matrix} \vdots & \vdots \\ \vdots & \vdots \end{matrix} \\
\hline
\begin{matrix} \vdots & \vdots \\ \vdots & \vdots \end{matrix} &
\begin{matrix} \ddots & \ddots \\ \ddots & \ddots \end{matrix} &
\begin{matrix} \ddots & \ddots \\ \ddots & \ddots \end{matrix} &
\begin{matrix} r & \cdots & \cdots & r \\ \vdots & \ddots & & \vdots \\ \vdots & & \ddots & \vdots \\ r & \cdots & \cdots & r \end{matrix} \\
\hline
\begin{matrix} r & \cdots & \cdots & r \\ \vdots & \ddots & & \vdots \\ \vdots & & \ddots & \vdots \\ r & \cdots & \cdots & r \end{matrix} &
\begin{matrix} \cdots & \cdots \\ \cdots & \cdots \end{matrix} &
\begin{matrix} r & \cdots & \cdots & r \\ \vdots & \ddots & & \vdots \\ \vdots & & \ddots & \vdots \\ r & \cdots & \cdots & r \end{matrix} &
\begin{matrix} p & q & \cdots & q \\ q & \ddots & & \vdots \\ \vdots & & \ddots & q \\ q & \cdots & q & p \end{matrix}
\end{array}\right)
\left(\begin{array}{c} 1 \\ 1 \\ \vdots \\ 1 \\ \hline -1 \\ \vdots \\ \vdots \\ -1 \\ \hline 0 \\ \vdots \\ \vdots \\ 0 \\ \hline 0 \\ \vdots \\ \vdots \\ 0 \end{array}\right) \\
&= \left(\begin{array}{c} \mu^{(1)} - \mu^{(2)} \\ \vdots \\ \vdots \\ \mu^{(1)} - \mu^{(2)} \\ \hline -\mu^{(1)} + \mu^{(2)} \\ \vdots \\ \vdots \\ -\mu^{(1)} + \mu^{(2)} \\ \hline 0 \\ \vdots \\ \vdots \\ 0 \\ \hline 0 \\ \vdots \\ \vdots \\ 0 \end{array}\right)
= \left(\begin{array}{c} 1 \\ 1 \\ \vdots \\ 1 \\ \hline -1 \\ \vdots \\ \vdots \\ -1 \\ \hline 0 \\ \vdots \\ \vdots \\ 0 \\ \hline 0 \\ \vdots \\ \vdots \\ 0 \end{array}\right) \delta\mu
\end{aligned}
$$

with

$$
\begin{aligned}
\delta\mu &= N_{\text{train}}(b-c)(p+(N_{\text{train}}-1)q)-(b-c)N_{\text{train}}^2 r \\
&= N_{\text{train}}(b-c)(p+(N_{\text{train}}-1)q-N_{\text{train}}r) \\
&= N_{\text{train}}(b-c)\left(p-q+N_{\text{train}}(q-r)\right) . \qquad (34)
\end{aligned}
$$

So for (I)-(II) and (II)-(III) in equation (33) we get

$$
\begin{aligned}
p(a-b)+bq-aq=1 \qquad &\Rightarrow p-q=\frac{1}{a-b} \\
a\,(q-r)+bp-(N_{\text{train}}-1)\,br+(N_{\text{train}}-2)bq-cp-(N_{\text{train}}-1)cq+N_{\text{train}}cr=0\,.&
\end{aligned}
$$

Rearranging the second line

$$
(a-b+N_{\text{train}}(b-c))\,(q-r)+(b-c)(p-q)=0
$$

leads to

$$
(q-r)=-\,(b-c)(p-q)\frac{1}{(a-b+N_{\text{train}}(b-c))}.
$$

Inserting this into equation (34), we obtain

$$
\begin{aligned}
\delta\mu &= N_{\text{train}}(b-c)\left(p-q+N_{\text{train}}(q-r)\right) \\
&= N_{\text{train}}(p-q)(b-c)\left(1-\frac{N_{\text{train}}(b-c)}{(a-b+N_{\text{train}}(b-c))}\right) \\
&= N_{\text{train}}(p-q)(b-c)\left(\frac{a-b}{(a-b+N_{\text{train}}(b-c))}\right) \\
&= \frac{N_{\text{train}}(b-c)}{(a-b)}\left(\frac{a-b}{(a-b+N_{\text{train}}(b-c))}\right) \\
&= \left(1+\frac{1}{N_{\text{train}}}\frac{a-b}{b-c}\right)^{-1} \\
&= \left(1+\frac{1}{N_{\text{train}}}\frac{\kappa+R-Q^{\circlearrowleft}}{Q^{\circlearrowleft}-Q^{\leftrightarrow}(P)}\right)^{-1}, \qquad (35)
\end{aligned}
$$

which does not depend on the $P-2$ factor in equation (33). In other words, $\delta\mu$ only depends on $P$ through the overlap between stimuli $c=g_w^2 Q^{\leftrightarrow}(P)$. Apart from this, it is the same expression as the separability for two stimuli in equation (26).

##### 5.1.2 Calculating the variance

Each readout has variability in its responses. It is quantified through the posterior covariance, capturing the uncertainty from the noise $\epsilon$ and potentially insufficient data to constrain $\boldsymbol{w}$. Let us use the notation

$$
\boldsymbol{\Sigma}(\boldsymbol{X}^*)=\boldsymbol{K}_{**}-\boldsymbol{K}_{*D}\boldsymbol{K}_{DD}^{-1}\boldsymbol{K}_{D*}.
$$

The posterior covariance is a matrix, but here we are only interested in the diagonal entry which corresponds to the variance of one test point. We can use a trick for the calculation in assuming that we have as many test points for each class as training points. The entries of the posterior covariance are independent of how many test points there are. Thus with $\boldsymbol{K}_{**} = \boldsymbol{K}_{DD}$, $\boldsymbol{K}_{*D} = \boldsymbol{K}_{D*} = \boldsymbol{K}_{DD} - (a-b)\mathbb{1}$

$$\begin{aligned}\boldsymbol{\Sigma} &= \boldsymbol{K}_{**} - \boldsymbol{K}_{*D}\boldsymbol{K}_{DD}^{-1}\boldsymbol{K}_{D*} \\ &= \boldsymbol{K}_{DD} - \left(\boldsymbol{K}_{DD} - (a-b)\mathbb{1}\right)\boldsymbol{K}_{DD}^{-1}\left(\boldsymbol{K}_{DD} - (a-b)\mathbb{1}\right) \\ &= -(a-b)^2\boldsymbol{K}_{DD}^{-1} + 2(a-b)\mathbb{1} \\ &= (a-b)^2\left(\frac{2}{a-b}\mathbb{1} - \boldsymbol{K}_{DD}^{-1}\right).\end{aligned}$$

If we are now interested in the variance $\Sigma(\boldsymbol{x}_1^*)$ (i.e. the diagonal entry of $\boldsymbol{K}_{**}$), we obtain

$$\Sigma(\boldsymbol{x}_1^*) = (a-b)^2\left(\frac{2}{a-b} - p\right).$$

Using the equations in (33), we can solve for $p, q, r$ as functions of $a, b, c$:

$$r = -\frac{c}{N_{\mathrm{train}}^2(b-c)^2}\frac{1}{1+\frac{a-b}{N_{\mathrm{train}}(b-c)}}\frac{1}{1+\frac{a-b}{N_{\mathrm{train}}(b-c)}+P\frac{c}{b-c}}$$

$$q = -\frac{1}{N_{\mathrm{train}}}\frac{1}{a-b}\frac{1}{1+\frac{a-b}{N_{\mathrm{train}}(b-c)}}\left[1+\frac{\frac{a-b}{N_{\mathrm{train}}(b-c)}\frac{c}{b-c}}{1+\frac{a-b}{N_{\mathrm{train}}(b-c)}+P\frac{c}{b-c}}\right]$$

$$p = \frac{1}{a-b}\left[1-\frac{1}{N_{\mathrm{train}}}\frac{1}{1+\frac{a-b}{N_{\mathrm{train}}(b-c)}}\left[1+\frac{\frac{a-b}{N_{\mathrm{train}}(b-c)}\frac{c}{b-c}}{1+\frac{a-b}{N_{\mathrm{train}}(b-c)}+P\frac{c}{b-c}}\right]\right].$$

Thus the posterior covariance evaluates to

$$\begin{aligned}\Sigma(\boldsymbol{x}_1^*) &= (a-b)\left\{1+\frac{1}{N_{\mathrm{train}}}\frac{1}{1+\frac{a-b}{N_{\mathrm{train}}(b-c)}}\left[1+\frac{\frac{a-b}{N_{\mathrm{train}}(b-c)}\frac{c}{b-c}}{1+\frac{a-b}{N_{\mathrm{train}}(b-c)}+P\frac{c}{b-c}}\right]\right\} \\ &=: \Sigma.\end{aligned}$$

The only $P$ dependence appears in the last term. As we saw in the main text, this dependence on $P$ is negligible and one can even approximate the posterior variance as $\Sigma(\boldsymbol{x}_1^*) \simeq a-b$ quite well. The scalar $\Sigma(\boldsymbol{x}_1^*)$ is what from hereon call $\Sigma$ in the following for ease of notation.

#### 5.2 Mutual information

In this section, we calculate the mutual information about the stimulus identity $C$ that can be linearly decoded from the neural responses $\boldsymbol{x}$. To this end, we will use the optimal Bayesian posterior derived in the last section, which resulted from inference of the Bayes-optimal readout $\boldsymbol{w}$. We consider $P$ equally likely stimuli

$$p(C) = \frac{1}{P}.$$

The response $\boldsymbol{r} \in \mathbb{R}^P$ is given by the predictor statistics for all readouts to the same test point $\boldsymbol{r}|C = \left(y_1^*(\boldsymbol{x}^*), \ldots, y_P^*(\boldsymbol{x}^*)\right)^{\mathrm{T}}_{C(\boldsymbol{x}^*)=C}$, which factorizes according to equation (31)

$$\begin{aligned} p(\boldsymbol{r}|C) &= \mathcal{N}\left(\boldsymbol{r}|\boldsymbol{\mu}_C, \Sigma\right) \\ &= \prod_{p=1}^{P} \mathcal{N}\left(r_p|\left(\boldsymbol{\mu}_C\right)_p, \Sigma\right) \end{aligned} \tag{36}$$

In the mean-field limit all readouts have the same response except for the readout that has been conditioned on the presented stimulus $C$:

$$\left(\boldsymbol{\mu}_C\right)_p = \mu^1 \delta_{Cp} + \left(1 - \delta_{Cp}\right)\mu^2.$$

We define $\delta\mu = \mu^1 - \mu^2$ as in equation (35) and introduce $\Delta \coloneqq \delta\mu/\sqrt{\Sigma}$. The larger $\Delta$, the easier it is to infer which stimulus was applied. In the limit $\Delta \to \infty$, the responses would be given by a delta peak each. In this case, the information would be transmitted perfectly.

The mutual information (MI) between $\boldsymbol{r}$ and $C$ (which parametrically depends on $P$ and $\Delta$ only, as we will show) quantifies the amount of information transmitted by $\boldsymbol{r}$ about $C$ and is defined as

$$\begin{aligned} \mathrm{MI}\left(P, \Delta\right) &= D_{KL}\left(p(C, \boldsymbol{r}) \| p(C)p(\boldsymbol{r})\right) \\ &= H(C) - H(C|\boldsymbol{r}), \end{aligned}$$

where $H$ denotes the (conditional) entropy of a random variable.

The upper bound of the MI is the entropy of stimuli

$$\begin{aligned} H(C) &= -\sum_{C=1}^{P} p(C)\log_2 p(C) \\ &= -\sum_{C=1}^{P} \frac{1}{P}\log_2\frac{1}{P} \\ &= \log_2 P. \end{aligned}$$

We thus have to evaluate the conditional entropy

$$\begin{aligned} H(C|\boldsymbol{r}) &= -\sum_{C=1}^{P} \int d\boldsymbol{r}\, p(\boldsymbol{r}, C)\log_2\left(\frac{p(\boldsymbol{r}, C)}{p(\boldsymbol{r})}\right) \\ &= -\sum_{C=1}^{P} \int d\boldsymbol{r}\, p(\boldsymbol{r}, C)\log_2\left(\frac{p(\boldsymbol{r}, C)}{\sum_{C'=1}^{P} p(\boldsymbol{r}, C')}\right) \\ &= -\sum_{C=1}^{P} \int d\boldsymbol{r}\, p(\boldsymbol{r}|C)p(C)\log_2\left(\frac{p(\boldsymbol{r}|C)p(C)}{\sum_{C'=1}^{P} p(\boldsymbol{r}|C')p(C')}\right) \\ &= -\sum_{C=1}^{P} \int d\boldsymbol{r}\, p(\boldsymbol{r}|C)p(C)\log_2\left(\frac{p(\boldsymbol{r}|C)}{\sum_{C'=1}^{P} p(\boldsymbol{r}|C')}\right) \\ &= -\frac{1}{P}\sum_{C=1}^{P} \int d\boldsymbol{r}\, p(\boldsymbol{r}|C)\log_2\left(\frac{p(\boldsymbol{r}|C)}{\sum_{C'=1}^{P} p(\boldsymbol{r}|C')}\right) \\ &\overset{\text{symmetry in } C}{=} -\int d\boldsymbol{r}\, p(\boldsymbol{r}|C=1)\log_2\left(\frac{p(\boldsymbol{r}|C=1)}{\sum_{C'=1}^{P} p(\boldsymbol{r}|C')}\right). \end{aligned}$$

The symmetry over all stimuli is used in the last step. We choose $C$ to be 1 without loss of generality. This is the placeholder for the readout whose response is different from all others. Inserting equation (36) yields

$$
\begin{aligned}
H(C|\boldsymbol{r}) &= -\int d\boldsymbol{r}\, p(\boldsymbol{r}|C=1)\log_2\left(\frac{p(\boldsymbol{r}|C=1)}{\sum_{C'=1}^{P} p(\boldsymbol{r}|C')}\right)\\
&= -\int d\boldsymbol{r}\,\mathcal{N}\left(\boldsymbol{r}|\boldsymbol{\mu}_{C=1},\Sigma\right)\log_2\left(\frac{\mathcal{N}\left(\boldsymbol{r}|\boldsymbol{\mu}_{C=1},\Sigma\right)}{\sum_{C'=1}^{P}\mathcal{N}\left(\boldsymbol{r}|\boldsymbol{\mu}_{C'},\Sigma\right)}\right)\\
&= \int d\boldsymbol{r}\,\mathcal{N}\left(\boldsymbol{r}|\boldsymbol{\mu}_{C=1},\Sigma\right)\log_2\left(\frac{\sum_{C'=1}^{P}\mathcal{N}\left(\boldsymbol{r}|\boldsymbol{\mu}_{C'},\Sigma\right)}{\mathcal{N}\left(\boldsymbol{r}|\boldsymbol{\mu}_{C=1},\Sigma\right)}\right)\\
&= \int d\boldsymbol{r}\,\mathcal{N}\left(\boldsymbol{r}|\boldsymbol{\mu}_{C=1},\Sigma\right)\log_2\left(1+\frac{\sum_{C'=2}^{P}\mathcal{N}\left(\boldsymbol{r}|\boldsymbol{\mu}_{C'},\Sigma\right)}{\mathcal{N}\left(\boldsymbol{r}|\boldsymbol{\mu}_{C=1},\Sigma\right)}\right)\\
&\overset{(36)}{=} \int d\boldsymbol{r}\,\mathcal{N}\left(\boldsymbol{r}|\boldsymbol{\mu}_{C=1},\Sigma\right)\log_2\left(1+\sum_{C'=2}^{P}\frac{\mathcal{N}\left(r_1|\mu^2,\Sigma\right)}{\mathcal{N}\left(r_1|\mu^1,\Sigma\right)}\frac{\mathcal{N}\left(r_{C'}|\mu^1,\Sigma\right)}{\mathcal{N}\left(r_{C'}|\mu^2,\Sigma\right)}\right)\\
&\overset{\boldsymbol{r}\to\boldsymbol{r}+\mu^2}{=} \int d\boldsymbol{r}\,\mathcal{N}\left(\boldsymbol{r}|\boldsymbol{\mu}_{C=1}-\mu^2,\Sigma\right)\log_2\left(1+\sum_{C'=2}^{P}\frac{\mathcal{N}\left(r_1|0,\Sigma\right)}{\mathcal{N}\left(r_1|\delta\mu,\Sigma\right)}\frac{\mathcal{N}\left(r_{C'}|\delta\mu,\Sigma\right)}{\mathcal{N}\left(r_{C'}|0,\Sigma\right)}\right)\\
&= \int d\boldsymbol{r}\,\mathcal{N}(r_1|\delta\mu,\Sigma)\left[\prod_{C=2}^{P}\mathcal{N}(r_C|0,\Sigma)\right]\log_2\left(1+\sum_{C'=2}^{P}\frac{\mathcal{N}\left(r_1|0,\Sigma\right)}{\mathcal{N}\left(r_1|\delta\mu,\Sigma\right)}\frac{\mathcal{N}\left(r_{C'}|\delta\mu,\Sigma\right)}{\mathcal{N}\left(r_{C'}|0,\Sigma\right)}\right)
\end{aligned}
$$

Lets simplify the term in the logarithm

$$
\begin{aligned}
&\frac{\mathcal{N}\left(r_1|0,\Sigma\right)}{\mathcal{N}\left(r_1|\delta\mu,\Sigma\right)}\frac{\mathcal{N}\left(r_{C'}|\delta\mu,\Sigma\right)}{\mathcal{N}\left(r_{C'}|0,\Sigma\right)}\\
=&\frac{\exp\left(-\frac{1}{2}r_1^2/\Sigma\right)}{\exp\left(-\frac{1}{2}\left(r_1-\delta\mu\right)^2/\Sigma\right)}\sum_{C'=2}^{P}\frac{\exp\left(-\frac{1}{2}\left(r_{C'}-\delta\mu\right)^2/\Sigma\right)}{\exp\left(-\frac{1}{2}r_{C'}^2/\Sigma\right)}\\
=&\exp\left(-\delta\mu\, r_1/\Sigma+\frac{1}{2}(\delta\mu)^2/\Sigma\right)\sum_{C'=2}^{P}\exp\left(\delta\mu\, r_{C'}/\Sigma-\frac{1}{2}(\delta\mu)^2/\Sigma\right)\\
=&\exp\left(-\delta\mu\, r_1/\Sigma\right)\sum_{C'=2}^{P}\exp\left(\delta\mu\, r_{C'}/\Sigma\right)
\end{aligned}
$$

Thus with the redefinition of the integration variable $\boldsymbol{r} \leftarrow \frac{\boldsymbol{r}}{\sqrt{\Sigma}}$ we arrive at

$$
\begin{aligned}
H(C|\boldsymbol{r}) &= \int d\boldsymbol{r}\,\mathcal{N}(r_1|\Delta,1)\left[\prod_{C=2}^{P}\mathcal{N}(r_C|0,1)\right]\log_2\left(1+\exp\left(-\Delta\, r_1\right)\sum_{C'=2}^{P}\exp\left(\Delta\, r_{C'}\right)\right)\\
&= \log_2(e)\int dr_1\,\mathcal{N}(r_1|\Delta,1)\left(-\Delta\, r_1\right) + \int d\boldsymbol{r}\,\mathcal{N}(r_1|\Delta,1)\left[\prod_{C=2}^{P}\mathcal{N}(r_C|0,1)\right]\log_2\left(\sum_{C=1}^{P}\exp\left(\Delta\, r_C\right)\right)\\
&= -\log_2(e)\Delta^2 + \int d\boldsymbol{r}\,\mathcal{N}(r_1|\Delta,1)\left[\prod_{C=2}^{P}\mathcal{N}(r_C|0,1)\right]\log_2\left(\sum_{C=1}^{P}\exp\left(\Delta\, r_C\right)\right),
\end{aligned}
$$

where we factored the logarithm towards the second line.

Thus, the mutual information indeed only depends on $\Delta$ and $P$

$$\mathrm{MI}(P,\Delta) = \log_2(e)\Delta^2 - \int dr_1 \cdots \int dr_P\, \mathcal{N}(r_1|\Delta,1)\left[\prod_{C=2}^{P}\mathcal{N}(r_C|0,1)\right]\log_2\left(\frac{1}{P}\sum_{C=1}^{P}\exp\left(\Delta\, r_C\right)\right), \tag{37}$$

where we absorbed the entropy $H(C) = -\log_2(1/P)$ in the last factor.

The calculation of the mutual information thus boils down to evaluating the multi-dimensional integral in equation (37). This is very expensive to do numerically using Monte-Carlo techniques even after exploiting symmetries of the formula, or even intractable. We approximate the sum of the log-normal variables as a log-normal distribution and use the approach of [10] to rewrite

$$\begin{aligned}\mathrm{MI}(P,\Delta) &= \log_2(e)\Delta^2 - \int dr_1 \cdots \int dr_P\, \mathcal{N}(r_1|\Delta,1)\left[\prod_{C=2}^{P}\mathcal{N}(r_C|0,1)\right]\log_2\left(\frac{1}{P}\sum_{C=1}^{P}\exp\left(\Delta\, r_C\right)\right) \\ &\approx \log_2(e)\Delta^2 - \log_2(e)\, m(\Delta(P),P).\end{aligned} \tag{38}$$

with the two dimensional Gaussian integral

$$m(\Delta(P),P) = \int dr_1 \int dS\, \mathcal{N}(r_1|\Delta,1)\,\mathcal{N}(S|\mu_S,\Sigma_S)\log_2\left(\frac{1}{P}\exp\left(\Delta\, r_1\right) + \frac{P-1}{P}\exp\left(S\right)\right).$$

Here, we have denoted the sum of identically and independently distributed log-normal variables $\exp\left(\Delta\, r_C\right)$ as

$$\exp\left(S\right) = \frac{1}{P-1}\sum_{C=2}^{P}\exp\left(\Delta\, r_C\right) \tag{39}$$

and approximate its probability distribution as a lognormal distribution, by assuming that $p(S) = \mathcal{N}(S|\mu_S,\Sigma_S)$. To evaluate the integral, we need to determine its cumulants $\mu_S$ and $\Sigma_S$.

To this end, we generalize the Yeh-Schwartz approach [10] of two log-normal random variables to our case of $P-1$ variables. We first recapitulate their ansatz: It consists of approximating the sum of two log-normal variables as a single log-normal variable. For two Gaussian variables $Y_1 \sim \mathcal{N}\left(m_{Y_1},\sigma_{Y_1}^2\right)$ and $Y_2 \sim \mathcal{N}\left(m_{Y_2},\sigma_{Y_2}^2\right)$, we take

$$\begin{aligned} Z &= \ln\left(e^{Y_1} + e^{Y_2}\right) \\ &= e^{Y_1} + \ln\left(1 + e^{w}\right)\end{aligned}$$

with $w \sim \mathcal{N}\left(m_w,\sigma_w^2\right) = \mathcal{N}\left(m_{Y_2} - m_{Y_1}, \sigma_{Y_1}^2 + \sigma_{Y_2}^2\right)$. The paper provides the iterative equations for the cumulants of $Z \sim \mathcal{N}\left(m_Z,\sigma_Z^2\right)$,

$$\begin{aligned} m_Z &= m_{Y_1} + G_1\left(\sigma_w, m_w\right) \\ \sigma_Z^2 &= \sigma_{Y_1}^2 - G_1^2\left(\sigma_w,m_w\right) - 2\rho^2 G_3\left(\sigma_w,m_w\right) + G_2\left(\sigma_w,m_w\right),\end{aligned}$$

with $\rho = -\sigma_{Y_1}/\sigma_w$. To avoid the problem of arithmetic underflow we use $e^{s^2}\left(1 + \mathrm{erf}\left(-s\right)\right) = \mathrm{erfcx}\left(s\right)$ in the calculations for the $G_1$,$G_2$ and $G_3$, such that (in the notation of [10])

$$\begin{aligned} C_k &= \frac{(-1)^{k+1}}{k} \\ b_k &= \frac{2(-1)^{k+1}}{k+1}\sum_{j=1}^{k} j^{-1}\end{aligned}$$

$$G_1\left(\sigma_w, m_w\right) = m_w \Phi\left(\frac{m_w}{\sigma_w}\right) + \frac{\sigma_w}{\sqrt{2\pi}} \exp\left(-\frac{1}{2}\frac{m_w^2}{\sigma_w^2}\right) + \frac{1}{2}\exp\left(-\frac{1}{2}\frac{m_w^2}{\sigma_w^2}\right)\sum_{k=1}^{\infty} C_k \left[\operatorname{erfcx}\left(\frac{m_w + k\sigma_w^2}{\sqrt{2}\sigma_w}\right) + \operatorname{erfcx}\left(\frac{-m_w + k\sigma_w^2}{\sqrt{2}\sigma_w}\right)\right]$$

$$G_2\left(\sigma_w, m_w\right) = B_1 + B_3 + B_4 + B_5$$

$$B_1 = \frac{1}{2}\sum_{k=1}^{\infty} b_k \exp\left(-\frac{1}{2}\frac{m_w^2}{\sigma_w^2}\right) \operatorname{erfcx}\left(\frac{+m_w + (k+1)\sigma_w^2}{\sqrt{2}\sigma_w}\right)$$

$$B_3 = \left(m_w^2 + \sigma_w^2\right)\Phi\left(\frac{m_w}{\sigma_w}\right) + \frac{1}{\sqrt{2\pi}} m_w \sigma_w \exp\left(-\frac{1}{2}\frac{m_w^2}{\sigma_w^2}\right)$$

$$B_4 = -2\exp\left(-\frac{1}{2}\frac{m_w^2}{\sigma_w^2}\right)\sum_{k=1}^{\infty} C_k \left(-m_w + k\sigma_w^2\right) \operatorname{erfcx}\left(\frac{-m_w + k\sigma_w^2}{\sqrt{2}\sigma_w}\right)$$

$$B_5 = \frac{1}{2}\sum_{k=1}^{\infty} b_k \exp\left(-\frac{1}{2}\frac{m_w^2}{\sigma_w^2}\right) \operatorname{erfcx}\left(\frac{-m_w + (k+1)\sigma_w^2}{\sqrt{2}\sigma_w}\right)$$

$$2\rho^2 G_3\left(\sigma_w, m_w\right) = A_2' + A_2''$$

$$A_2' = 2\rho^2 \frac{1}{2}\sigma_w^2 \exp\left(-\frac{1}{2}\frac{m_w^2}{\sigma_w^2}\right)\sum_{k=0}^{\infty} (-1)^k \operatorname{erfcx}\left(\frac{m_w + (k+1)\sigma_w^2}{\sqrt{2}\sigma_w}\right)$$

$$A_2'' = 2\rho^2 \frac{1}{2}\sigma_w^2 \exp\left(-\frac{1}{2}\frac{m_w^2}{\sigma_w^2}\right)\sum_{k=0}^{\infty} (-1)^k \operatorname{erfcx}\left(\frac{-m_w + k\sigma_w^2}{\sqrt{2}\sigma_w}\right).$$

The sums over $k$ are carried out until the contributions become negligible, which is $k = 20$ for the plots in the manuscript. This procedure is extended to $P-1$ log-normal variables by iteration.

##### 5.2.1 The log-normal approximation: numerics

The above approximation technique is chosen over other possible approaches, such as a Gaussian approximation or an approximation using a third cumulant to account for the skewness. The reason is that we find it to empirically work also for smaller $P$ and larger $\Delta$.

The approximation fits well for small $P$ and converges to the right distribution for very large $P$, see Figure 8. The shape of the distribution may deviate for intermediate $P$ and larger $\Delta$. This is not a problem since $\Delta(P)$ depends on $P$ and decreases with increasing $P$ because of the crowding of stimuli in neuron space. For larger $\Delta$ the distribution only becomes peaked for larger $P$. The other commonly used Fenton-Wilkinson approximation [11] is insufficient as it becomes inaccurate for larger $\Delta$, see Figure 8.

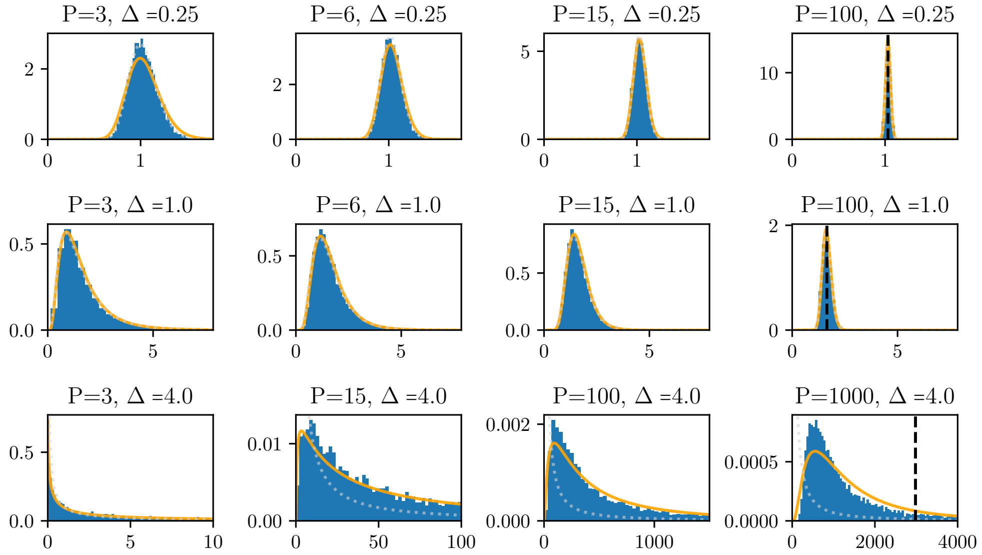

Figure 8: **Distribution for the sum of log-normal variables.** Distribution of $S$ appearing in equation (38) for different values of $\Delta$, $P$ for the evaluation of the mutual information. Numerics (blue) compared to theory (orange), see equation (39). The black, dashed line in the right-most panels indicate the value around which the distribution concentrates for infinitely many stimuli, see equation (40). The dotted light-gray line corresponds to the Fenton-Wilkinson approximation, which fixes the parameters of the lognormal distribution by matching it to the first two cumulants of equation (39).

#### 5.3 Mutual information saturates for large number of stimuli

Here we calculate the mutual information in the limit of many stimuli $P$. Sums of i.i.d. variables concentrate around some value for large $P$. Let us define the probability distribution in the limit of many stimuli $p(S)$

$$\mathrm{MI}(P,\Delta) \overset{P\to\infty}{\longrightarrow} \log_2(e)\Delta^2 - \int dS\, p(S)\, \log_2\left(\exp\left(S\right)\right)$$

For large $P$, the mean of $\sum_{C=1}^{P} \exp\left(\Delta\, r_C\right)$ will dominate over all other cumulants. Calculating the mean gives

$$\begin{aligned}
&\left\langle \frac{1}{P}\sum_{C=1}^{P} \exp\left(\Delta\, r_C\right)\right\rangle_{r_1,\ldots,r_P} \\
=&\frac{1}{P}\left[\left\langle \exp\left(\Delta\, r_1\right)\right\rangle_{r_1} + \sum_{C=2}^{P} \left\langle \exp\left(\Delta\, r_C\right)\right\rangle_{r_C}\right] \\
=&\frac{1}{P}\left[\left\langle \exp\left(\Delta\, r_1\right)\right\rangle_{r_1} + (P-1)\left\langle \exp\left(\Delta\, r_2\right)\right\rangle_{r_2}\right] \\
=&\frac{1}{P}\left[\exp\left(\frac{3}{2}\Delta^2\right) + (P-1)\exp\left(\frac{1}{2}\Delta^2\right)\right] \\
\to&\exp\left(\frac{1}{2}\Delta^2\right).
\end{aligned} \tag{40}$$

In the penultimate line we identified the moment generating functions of a Gaussian variable. Because this term has the particular form of $\frac{1}{P}\sum_{C=1}^{P} z_C$ with $z_C$ drawn i.i.d from regular probability distributions, the $k$-th

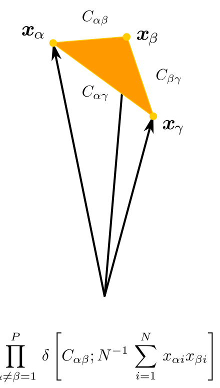

Figure 9: **Mutual overlaps of stimulus vectors.** Visualization of the second Dirac-$\delta$ term in equation (42), which constrains the mutual overlaps of binary vectors. The three binary vectors represent the average responses to three different stimuli, respectively. In other words, they correspond to the centers of the ellipsoids of the neural representations. In equation (42), we sum over all possible vector configurations and pick out the configuration that matches the previously specified target correlation matrix $C$.

cumulant goes as $\propto P^{-k+1}$, such that the mean dominates for large $P$. So eventually the distribution will have the mean calculated above,

$$\int dS\, p(S) \log_2 (\exp(S)) \overset{P\to\infty}{\longrightarrow} \log_2 \left( \exp\left( \frac{1}{2} \Delta^2 \right) \right).$$

and thus

$$\begin{aligned} \mathrm{MI}(P, \Delta) &\overset{P\to\infty}{\longrightarrow} \log_2(e)\Delta^2 - \frac{1}{2}\log_2(e)\Delta^2 \\ &= \frac{1}{2}\log_2(e)\Delta^2 . \end{aligned}$$

Note that lognormal distributions are notoriously slow in converging to a Gaussian due to their heavy tails. The larger $\Delta$, the more pronounced the tail and the more slowly it saturates, see Figure 8.

**Why does the mutual information saturate?**

An intuition for why mutual information eventually saturates can be drawn from the observation that deciphering the stimulus class relies heavily on analyzing the readouts, with particular focus on the first readout. It shows the highest mean response and is distributed according to $p(r_1) = \mathcal{N}\left(r_1|\mu^1, \Sigma\right)$. If $r_1$ takes on a particular value, the values of all other readouts distributed as $p(r_{2:P}) = \mathcal{N}\left(r_{2:P}|\mu^2, \Sigma\right)$ densely populate the vicinity of $r_1$ as $P$ goes to infinity. This blurs the information carried by the first readout and makes it harder to infer the stimulus class. One realization of one stimulus application carries no information anymore for infinitely many stimuli. On the other hand, we get infinitely many chances of inferring stimulus classes as $P$ goes to infinity. The balance between these two effects lets the mutual information saturate.

#### 5.4 Optimal overlap between stimulus classes

Here, we want to calculate the overlap between different stimuli under the assumption that the brain efficiently distributes different stimuli by allocating different regions in neuron space to them. In other words, our presumption is that the overlap between stimulus classes $Q^{\leftrightarrow}$ in neuron space is smaller than the average overlap one gets for uncorrelated random binary vectors. One stimulus is represented by a binary vector $\boldsymbol{x}_\alpha \in \mathbb{R}^N$. If the state of stimulus $\alpha$ of the $i$-th neuron is independently and identically distributed as $x_{\alpha i} \overset{\text{i.i.d}}{\sim} \text{Bernoulli}(R_i)$ with $R_i$ being the firing rate of neuron $i$, or equivalently, the probability of the neuron to fire, then we would have for $\alpha \neq \beta$

$$\langle x_{\alpha i} x_{\beta i} \rangle = R_i^2.$$

Thus the overlap between different stimuli would be

$$\begin{aligned} Q^{\leftrightarrow} &= \left\langle \frac{1}{N} \sum_{i=1}^{N} x_{\alpha i} x_{\beta i} \right\rangle \\ &= \frac{1}{N} \sum_{i=1}^{N} R_i^2. \end{aligned} \tag{41}$$

However, we experimentally observe a smaller overlap, suggesting that the distribution of stimuli is more efficient in the brain. But how much more efficient?

##### Number of vectors with fixed number of ones and overlap

To answer the previous question we need to count the number of possible representations that realize a more beneficial distribution in stimulus space. Analyzing how this number depends on the mutual overlap allows us to determine the minimal overlap possible that still leads to a solution. We would therefore like to determine the number of sets containing vectors $\boldsymbol{x}_{1\le\alpha\le P} \in \{0,1\}^N$ with the following properties:

1. The (normalized) pairwise overlap between any pair of vectors should be $Q^{\leftrightarrow} = N^{-1} \sum_{i=1}^{N} x_{\alpha i}\, x_{\beta i} \quad \forall \alpha \neq \beta$ .
2. The (normalized) number of ones for each neuron should be $R = \frac{1}{N} \sum_{i=1}^{N} x_{\alpha i} \quad \forall \alpha$.
3. Furthermore, we want to enforce the experimentally observed firing rate distribution $\{R_k\}_{1\le k\le N_{\text{exp}}}$:
   (a) We do this by dividing all $N$ neurons into $N_{\text{exp}}$ groups with $f = \frac{N}{N_{\text{exp}}}$ neurons each.
   (b) The $k$-th group $f_k$ should have a firing rate of $R_k = f^{-1} \sum_{i\in f_k} x_{\alpha i}$.
   (c) We order the neurons in $x_\alpha$ in such a way that the first $f$ entries belong to the first group $f_1$, the entries from there up to $2f$ to the second group $f_2$ and so on.

The number of solutions is then given by the partition function

$$Z(C,j) := \sum_{\{x_{\alpha i}\}\in\{0,1\}^{P\times N}} \left\{ \prod_{\alpha=1}^{P} \prod_{k=1}^{N_{\text{exp}}} \delta\left[ j_{\alpha k}; f^{-1} \sum_{i\in f_k} x_{\alpha i} \right] \right\} \left\{ \prod_{\alpha\neq\beta=1}^{P} \delta\left[ C_{\alpha\beta}; N^{-1} \sum_{i=1}^{N} x_{\alpha i} x_{\beta i} \right] \right\} \tag{42}$$

where $\delta[\circ;\circ]$ is the Kronecker-$\delta$. The $j_{\alpha k}$ tunes the firing rates of the neurons in group $f_k$ and the matrix $C$ tunes the mutual overlaps between neural vectors $\boldsymbol{x}_\alpha$. The first term in the curly braces enforces condition 2 and 3 and the second term condition 1, respectively.

The expression in equation (42) has the form of an unnormalized probability distribution for $j$, $C$, where the $x_{\alpha i}$ can be regarded as binary random variables with a flat measure.

#### Outline of derivation

In the following calculation, we will approximate $Z(C,j)$ by determining an effective action $\Gamma(C,j)$ such that $Z(C,j) \simeq e^{-\Gamma(C,j)}$ in the large $N$ limit up to proportionality. Intuitively, this is possible because $R = N^{-1}\sum_i^N x_{\alpha i}$ and $Q = N^{-1}\sum_i^N x_{\alpha i}x_{\beta i}$ are macroscopic variables comprised of a sum of single neuron terms which concentrates as $N \to \infty$. In this limit, the effective action is obtained as the Legendre transform of $\ln Z(C,j)$ in both $j$ and $C$, therefore sometimes called *second* Legendre transform [12, 13]. To determine the effective action, we adopt an approach from statistical field theory similar to [14, 4]: First, we will rewrite the $\delta$-constraints by introducing auxiliary fields $\tilde{j}$, $\tilde{C}$.

Because the first $\delta$-constraint on the firing rates $1/N\sum_i^N x_{\alpha i}$ is linear in $x$, it is possible to obtain the Legendre transform with respect to $\tilde{j}$ in analytical form for $\tilde{C} = 0$. For the second Legendre transform, we treat $\tilde{C}_{\alpha\beta}$ which appears as an effective coupling $x_{\alpha i}\tilde{C}_{\alpha\beta}x_{\beta i}$ perturbatively, paralleling the treatment of the Ising model in [12]. This will finally yield the effective action such that $Z(C,j) \propto p(j)\,p(C|j)$ with $p(C|j) \propto \mathcal{N}(C,j|\mu_C,\Sigma_C)$ in the form of a Gaussian (equation (49)), where $\Sigma_C(P)$ depends on the number of stimuli $P$. We will then use the experimentally observed $Q^{\leftrightarrow}(P=2)$ to fit a free parameter $E\left(Q^{\leftrightarrow}(P=2)\right)$ in this theory. Overall, this will enable us to extrapolate to $Q^{\leftrightarrow}(P>2)$, and thereby through equation (37), $\Delta = \delta\mu/\sqrt{\Sigma}$, and equation (35) calculate the mutual information for $P > 2$.

#### Transforming the $\delta$-constraint

Expressing the Kronecker-$\delta$ with help of the Dirac-$\delta$ as

$$\delta[a;b] = \lim_{\epsilon\searrow 0}\int_{-\epsilon}^{\epsilon}\delta(a-b-\iota)\,d\iota$$

and the latter by its Fourier representation $\delta(x) = \int_{-i\infty}^{i\infty}\frac{d\tilde{x}}{2\pi i}e^{\tilde{x}x}$, one has

$$\begin{aligned}Z(C,j) = \int d^{P\times P}\iota \int \mathcal{D}\tilde{j}\int\mathcal{D}\tilde{C}\sum_{\{x_{\alpha i}\}\in\{0,1\}^{P\times N}}&\exp\left(-\sum_{\alpha=1}^{P}\sum_{k=1}^{N_{\mathrm{exp}}}\tilde{j}_{\alpha k}(j_{\alpha k}+\iota_{\alpha k})-\sum_{\alpha\neq\beta=1}^{P}\tilde{C}_{\alpha\beta}(C_{\alpha\beta}+\iota_{\alpha\beta})\right)\\ &\times\exp\Big(\sum_{\alpha=1}^{P}\sum_{k=1}^{N_{\mathrm{exp}}}\tilde{j}_{\alpha k}\left[f^{-1}\sum_{i\in f_k}x_{\alpha i}\right]+\sum_{\alpha\neq\beta=1}^{P}\tilde{C}_{\alpha\beta}\,N^{-1}\sum_{i=1}^{N}x_{\alpha i}x_{\beta i}\Big)\end{aligned}$$

with $\int\mathcal{D}\tilde{j} = \prod_{\alpha=1}^{P}\prod_{k=1}^{N_{\mathrm{exp}}}\int\frac{d\tilde{j}_{\alpha k}}{2\pi i}$, $\int\mathcal{D}\tilde{C} = \prod_{\alpha\neq\beta=1}^{P}\int\frac{d\tilde{C}_{\alpha\beta}}{2\pi i}$, and $\int d^{P\times P}\iota = \lim_{\epsilon\searrow 0}\int_{-\epsilon}^{\epsilon}d^{P\times P}\iota$. We can simplify the expression in the second line by pulling out the sum over the neuron index $i$

$$\begin{aligned}&\sum_{\{x_{\alpha i}\}\in\{0,1\}^{P\times N}}\exp\Big(\sum_{\alpha=1}^{P}\sum_{k=1}^{N_{\mathrm{exp}}}\tilde{j}_{\alpha k}\left[f^{-1}\sum_{i\in f_k}x_{\alpha i}\right]+\sum_{\alpha\neq\beta=1}^{P}\tilde{C}_{\alpha\beta}\,N^{-1}\sum_{i=1}^{N}x_{\alpha i}x_{\beta i}\Big)\\ &=\sum_{\{x_{\alpha i}\}\in\{0,1\}^{P\times N}}\exp\Big(\sum_{\alpha=1}^{P}\sum_{i=1}^{N}f^{-1}\tilde{j}_{\alpha k_i}x_{\alpha i}+\sum_{\alpha\neq\beta=1}^{P}\tilde{C}_{\alpha\beta}\,N^{-1}\sum_{i=1}^{N}x_{\alpha i}x_{\beta i}\Big)\\ &=\sum_{\{x_{\alpha i}\}\in\{0,1\}^{P\times N}}\prod_{i=1}^{N}\exp\Big(\sum_{\alpha=1}^{P}f^{-1}\tilde{j}_{\alpha k_i}x_{\alpha i}+\sum_{\alpha\neq\beta=1}^{P}\tilde{C}_{\alpha\beta}\,N^{-1}x_{\alpha i}x_{\beta i}\Big).\end{aligned}$$

We can pull out this sum over neurons $\sum_{i=1}^{N}$ because we can reconstruct the group dependence $k$ of neuron $i$ indexed by $k_i = \lceil\frac{i}{f}\rceil$ (see condition 3). Therefore we could write $\sum_{k=1}^{N_{\mathrm{exp}}}\sum_{i\in f_k}\tilde{j}_{\alpha k} = \sum_{k=1}^{N_{\mathrm{exp}}}\sum_{i\in f_k}\tilde{j}_{\alpha k_i} = \sum_{i=1}^{N}\tilde{j}_{\alpha k_i}$ such that the sources $\tilde{j}_{\alpha k_i}$ only depend on $1\le i\le N$. We can swap the product and the sum to obtain the cumulant-generating function $\mathcal{W}\left(\tilde{C},\tilde{j}\right)$ from

$$
\begin{aligned}
&\sum_{\{x_{\alpha i}\}\in\{0,1\}^{P\times N}} \prod_{i=1}^{N} \exp\Big( \sum_{\alpha=1}^{P} f^{-1}\tilde{j}_{\alpha k_i} x_{\alpha i} + \sum_{\alpha\neq\beta=1}^{P} \tilde{C}_{\alpha\beta}\, N^{-1}\, x_{\alpha i} x_{\beta i}\Big) \\
=&\prod_{i=1}^{N} \sum_{\{x_{\alpha}\}\in\{0,1\}^{P}} \exp\Big( \sum_{\alpha=1}^{P} f^{-1}\tilde{j}_{\alpha k_i} x_{\alpha} + \sum_{\alpha\neq\beta=1}^{P} \tilde{C}_{\alpha\beta}\, N^{-1}\, x_{\alpha} x_{\beta}\Big) \\
=&\exp\left( \sum_{i=1}^{N} \ln\left( \sum_{\{x_{\alpha}\}\in\{0,1\}^{P}} \exp\Big( \sum_{\alpha=1}^{P} f^{-1}\tilde{j}_{\alpha k_i} x_{\alpha} + \sum_{\alpha\neq\beta=1}^{P} \tilde{C}_{\alpha\beta}\, N^{-1}\, x_{\alpha} x_{\beta}\Big)\right)\right) \\
=&\exp\big(\mathcal{W}\big(\tilde{C},\tilde{j}\big)\big)
\end{aligned}
$$

This cumulant-generating function contains all statistical information about the firing rates of all neurons and the mutual overlaps of neural state vectors. With the definition of the cumulant generating function, we arrive at the partition function

$$
Z(C,j) = \int d^{P\times P}\iota \int \mathcal{D}\tilde{j} \int \mathcal{D}\tilde{C} \sum_{\{x_{\alpha i}\}\in\{0,1\}^{P\times N}} \exp\left( -\sum_{\alpha=1}^{P}\sum_{k=1}^{N_{\text{exp}}} \tilde{j}_{\alpha k}(j_{\alpha k}+\iota_{\alpha k}) - \sum_{\alpha\neq\beta=1}^{P} \tilde{C}_{\alpha\beta}(C_{\alpha\beta}+\iota_{\alpha\beta})\right) \exp\big(\mathcal{W}\big(\tilde{C},\tilde{j}\big)\big)\,. \tag{43}
$$

**Reduction of the cumulant-generating function**

As stated above, we can decompose the total cumulant-generating function

$$
\begin{aligned}
\mathcal{W}\big(\tilde{C},\tilde{j}\big) &= \sum_{i=1}^{N} w\big(\tilde{C}/N, \big\{\tilde{j}_{\alpha k_i}\big\}_{\alpha}/N\big) \\
&= \sum_{k=1}^{N_{\text{exp}}} \sum_{i\in f_k} w\big(\tilde{C}/N, \big\{\tilde{j}_{\alpha k_i}\big\}_{\alpha}/N\big) \\
&= f \sum_{k=1}^{N_{\text{exp}}} w\big(\tilde{C}/N, \big\{\tilde{j}_{\alpha k}\big\}_{\alpha}/N\big)
\end{aligned}
$$

into a sum of small cumulant-generating functions with $k$ - dependent sources $\tilde{j}_{\alpha k}$

$$
w\big(\tilde{C}/N, \big\{\tilde{j}_{\alpha k}\big\}_{\alpha}/N\big) = \ln\left( \sum_{\{x_{\alpha}\}\in\{0,1\}^{P}} \exp\Big( \sum_{\alpha=1}^{P} \frac{N_{\text{exp}}}{N}\tilde{j}_{\alpha k} x_{\alpha} + \sum_{\alpha\neq\beta=1}^{P} \tilde{C}_{\alpha\beta}\, N^{-1}\, x_{\alpha} x_{\beta}\Big)\right). \tag{44}
$$

The expression looks like an Ising action with fields $f^{-1}\tilde{j}_{\alpha k_i}$ and interaction strengths $N^{-1}\tilde{C}_{\alpha\beta}$. In particular, a stronger $\tilde{C}_{\alpha\beta}$ (enforcing the imposed value $C_{\alpha\beta}$) tends to align $x_{\alpha}$ and $x_{\beta}$, increasing their correlation.

**Finding saddles for the large $N$ limit**

Since we can take the limit $N \to \infty$ of the scaled cumulant-generating function

$$
\begin{aligned}
\lambda(l,m) &= \lim_{N\to\infty}\frac{1}{N}\mathcal{W}(Nl,\,Nm)\\
&= \lim_{N\to\infty}\frac{1}{N}\frac{N}{N_{\mathrm{exp}}}\sum_{k=1}^{N_{\mathrm{exp}}} w\left(l,\{m_{\alpha k}\}_{\alpha}\right)\\
&= \lim_{N\to\infty}\frac{1}{N_{\mathrm{exp}}}\sum_{k=1}^{N_{\mathrm{exp}}} w\left(l,\{m_{\alpha k}\}_{\alpha}\right)\\
&= \frac{1}{N_{\mathrm{exp}}}\sum_{k=1}^{N_{\mathrm{exp}}} w\left(l,\{m_{\alpha k}\}_{\alpha}\right)
\end{aligned}
$$

the probability distribution over $\tilde{j}$ and $\tilde{C}$ concentrates in the limit of many neurons so that we can use the Gaertner-Ellis theorem [15] to replace the integrals over $\tilde{j}$ and $\tilde{C}$ by their most dominant values:

$$
\begin{aligned}
Z(C,j) &\overset{\text{l.d.p}}{\simeq} \int d^{P\times P}\iota\,\exp\big(-\Gamma(C+\iota,\,j+\iota)\big),\\
\Gamma(C,j) &:= \sup_{\tilde{C},\tilde{j}}\left\{\sum_{\alpha=1}^{P}\sum_{k=1}^{N_{\mathrm{exp}}}\tilde{j}_{\alpha k}j_{\alpha k}+\sum_{\alpha\neq\beta=1}^{P}\tilde{C}_{\alpha\beta}C_{\alpha\beta}-\mathcal{W}(\tilde{C},\tilde{j})\right\}
\end{aligned}
\tag{45}
$$

For finding the supremum we have to take the derivatives by $\tilde{j}$ and $\tilde{C}$. Because $\mathcal{W}\left(\tilde{C},\tilde{j}\right)$ is the cumulant-generating function, these derivatives yield the mean of $j$ and $C$, respectively. We use the Legendre transform with respect to $\tilde{j}$ in equation (45) to enforce the experimentally found firing rate distribution

$$
R_k \overset{!}{=} \frac{\partial\mathcal{W}\left(\tilde{C},\tilde{j}\right)}{\partial\tilde{j}_{\alpha k}}\quad\forall\,\alpha.
$$

**Uncoupled system ($\tilde{C}=0$)**

Because of the similarity to an Ising system in equation (44), we can think of $\tilde{C}/N$ as the coupling between spins. For an uncoupled system ($\tilde{C}=0$), the cumulant generating function is given by

$$
\begin{aligned}
&\mathcal{W}\big(0,\tilde{j}\big)\\
=&f\sum_{k=1}^{N_{\mathrm{exp}}} w\left(0,\left\{\tilde{j}_{\alpha k}\right\}_{\alpha}/N\right)\\
=&f\sum_{k=1}^{N_{\mathrm{exp}}} \ln\left(\sum_{\{x_\alpha\}\in\{0,1\}^P}\exp\Big(\sum_{\alpha=1}^{P}\frac{N_{\mathrm{exp}}}{N}\tilde{j}_{\alpha k}x_\alpha\Big)\right)\\
=&f\sum_{k=1}^{N_{\mathrm{exp}}} \ln\left(\sum_{\{x_\alpha\}\in\{0,1\}^P}\prod_{\alpha=1}^{P}\exp\Big(\frac{N_{\mathrm{exp}}}{N}\tilde{j}_{\alpha k}x_\alpha\Big)\right)\\
=&f\sum_{k=1}^{N_{\mathrm{exp}}}\sum_{\alpha=1}^{P} \ln\left(\sum_{\{x_\alpha\}\in\{0,1\}}\exp\Big(\frac{N_{\mathrm{exp}}}{N}\tilde{j}_{\alpha k}x_\alpha\Big)\right)\\
=&f\sum_{k=1}^{N_{\mathrm{exp}}}\sum_{\alpha=1}^{P} \ln\left(1+\exp\Big(\frac{N_{\mathrm{exp}}}{N}\tilde{j}_{\alpha k}\Big)\right)\\
=&f\sum_{k=1}^{N_{\mathrm{exp}}}\sum_{\alpha=1}^{P} \ln\Big(1+\exp\big(f^{-1}\tilde{j}_{\alpha k}\big)\Big)\,.
\end{aligned}
$$

Demanding

$$
R_k \overset{!}{=} \frac{\partial\mathcal{W}\big(\tilde{C},\tilde{j}\big)}{\partial\tilde{j}_{\alpha k}} = \frac{\exp\big(f^{-1}\tilde{j}_{\alpha k}\big)}{\big(1+\exp\big(f^{-1}\tilde{j}_{\alpha k}\big)\big)}
$$

lets us determine the value of the sources $\tilde{j}^{*}$ that fulfill this condition

$$
\Rightarrow\ \tilde{j}^{*}_{\alpha k}(R_k) = f\ln\left[R_k/(1-R_k)\right] \quad \forall\,\alpha.
$$

If we call the object after this first Legendre transform $\mathcal{V}$

$$
\begin{aligned}
\mathcal{V}\big(\tilde{C},\{R_k\}_k\big) &= \sup_{\tilde{j}_{\alpha k}}\left\{\sum_{\alpha,k}\tilde{j}_{\alpha k}R_k-\mathcal{W}(\tilde{C},\tilde{j})\right\}\\
&= \sum_{k=1}^{N_{\mathrm{exp}}}\sup_{\tilde{j}_{\alpha k}}\left\{\sum_{\alpha=1}^{P}\tilde{j}_{\alpha k}R_k-\sum_{i\in f_k} w\left(\tilde{C}/N,\left\{\tilde{j}_{\alpha k}\right\}_{\alpha}/N\right)\right\}\\
&= \sum_{k=1}^{N_{\mathrm{exp}}}\left\{\sum_{\alpha=1}^{P}\tilde{j}^{*}_{\alpha k}R_k-\sum_{i\in f_k} w\left(\tilde{C}/N,\left\{\tilde{j}^{*}_{\alpha k}\right\}_{\alpha}/N\right)\right\}\\
&= f\sum_{k=1}^{N_{\mathrm{exp}}} v\left(\tilde{C}/N,R_k\right),
\end{aligned}
$$

we see that it also decomposes into a sum of small $v$. For an uncoupled system ($\tilde{C}=0$) we arrive at

$$\begin{aligned}
\mathcal{V}\left(0,\{R_k\}_k\right) &= \sum_{k=1}^{N_{\mathrm{exp}}}\sum_{\alpha=1}^{P}\tilde{j}^*_{\alpha k}(R_k)R_k - f\sum_{k=1}^{N_{\mathrm{exp}}} w\left(0,\left\{\tilde{j}^*_{\alpha k}(R_k)\right\}_\alpha/N\right)\\
&= P\sum_{k=1}^{N_{\mathrm{exp}}}\left[\tilde{j}^*_{\alpha k}(R_k)R_k - f\,w\left(0,\left\{\tilde{j}^*_{\alpha k}(R_k)\right\}_\alpha/N\right)\right]\\
&= P\sum_{k=1}^{N_{\mathrm{exp}}}\left\{f\ln\left[R_k/(1-R_k)\right]R_k - f\ln\left[1+R_k/(1-R_k)\right]\right\}\\
&= P f\sum_{k=1}^{N_{\mathrm{exp}}}\left[\ln\left(R_k\right)R_k - \ln\left(1-R_k\right)R_k - \ln\left[1/(1-R_k)\right]\right]\\
&= P f\sum_{k=1}^{N_{\mathrm{exp}}}\left[\ln(R_k)\,R_k + (1-R_k)\ln(1-R_k)\right]\\
&= P\sum_{i=1}^{N}\left[\ln(R_{k_i})\,R_{k_i} + (1-R_{k_i})\ln(1-R_{k_i})\right],
\end{aligned}$$

which is the entropy of $NP$ independent binary variables with mean values $R_k$ for every $f$ neurons, as it should be, because the first Legendre transform constrains the mean only.

##### Perturbative treatment of coupling $\tilde{C}$

For a coupled system ($\tilde{C} \neq 0$), we do a perturbative approach:

To arrive at the exponent of the probability density we have to perform a second Legendre transform

$$\Gamma(C,\{R_k\}_k) = \sup_{\tilde{C}}\sum_{\alpha\neq\beta=1}^{P}\tilde{C}_{\alpha\beta}C_{\alpha\beta} + \mathcal{V}(\tilde{C},\{R_k\}_k). \tag{46}$$

Having fixed the mean of the probability distribution we now take the coupling into consideration. Using the diagrammatics of a non-Gaussian theory [12], we expand the first Legendre transform $\mathcal{V}(\tilde{C},\{R_k\}_k)$ in the couplings $\tilde{C}_{\alpha\beta}$. Due to the decomposition into a sum, we can do so for each of the $N$ terms. To evaluate the diagrammatic contributions up to second order, the necessary cumulants are the mean and the variance

$$\langle\!\langle x_{\alpha i}\rangle\!\rangle = \langle x_{\alpha i}\rangle = R_{k_i}$$

$$\begin{aligned}
&\langle\!\langle x_{\alpha i}^2\rangle\!\rangle\\
&=\left\langle x_{\alpha i}^2\right\rangle - \langle x_{\alpha i}\rangle^2\\
&=\langle x_{\alpha i}\rangle\left(1-\langle x_{\alpha i}\rangle\right)\\
&=R_{k_i}\left(1-R_{k_i}\right).
\end{aligned}$$

Up to second order (also referred to as TAP-approximation [16]), we get

$$\begin{aligned}\mathcal{V}\left(\tilde{C},\{R_k\}_k\right) = \mathcal{V}\left(0,\{R_k\}_k\right) \\ &- \sum_{\alpha\neq\beta=1}^{P} \frac{\tilde{C}_{\alpha\beta}}{N} \sum_{i=1}^{N} R_{k_i}^2 \\ &- \sum_{\alpha\neq\beta=1}^{P} \left(\frac{\tilde{C}_{\alpha\beta}}{N}\right)^2 \sum_{i=1}^{N} R_{k_i}^2 (1-R_{k_i})^2 + \mathcal{O}\left(\left(\frac{\tilde{C}_{\alpha\beta}}{N}\right)^3\right).\end{aligned} \tag{47}$$

With the second Legendre transform we want to enforce our desired correlation $Q$ between representations of different stimuli

$$Q \overset{!}{=} \frac{\partial \mathcal{W}\left(\tilde{C},\tilde{j}\right)}{\partial \tilde{C}_{\alpha\beta}} \;\forall\, \alpha \neq \beta\,.$$

The supremum condition hence yields

$$\begin{aligned}0 &\overset{!}{=} \frac{\partial}{\partial \tilde{C}_{\gamma\delta}} \Big( \sum_{\alpha\neq\beta=1}^{P} \tilde{C}_{\alpha\beta} Q + \mathcal{V}(\tilde{C},\{R_k\}_k)\Big) \\ &\simeq Q - \frac{1}{N}\sum_{i=1}^{N} R_{k_i}^2 - 2N^{-1}\tilde{C}_{\gamma\delta}\,\frac{1}{N}\sum_{i=1}^{N} R_{k_i}^2 (1-R_{k_i})^2,\end{aligned}$$

which is a linear equation for $\tilde{C}_{\alpha\beta} \quad \forall \alpha \neq \beta$ with the solution

$$\frac{\tilde{C}^*_{\gamma\delta}}{N} = \frac{1}{2}\,\frac{Q - \frac{1}{N}\sum_{i=1}^{N} R_{k_i}^2}{\frac{1}{N}\sum_{i=1}^{N} R_{k_i}^2 (1-R_{k_i})^2}, \tag{48}$$

which measures the deviation of the second moment $Q$ from the first moment squared relative to the squared variance of binary variables. In particular, we see that negative couplings $\tilde{C}_{\gamma\delta}/N$ are needed to keep $Q$ below the random overlap $\frac{1}{N}\sum_{i=1}^{N} R_{k_i}^2$, corresponding to an anti-correlating force $x_\alpha \tilde{C}_{\alpha\beta}/N\, x_\beta$ in the Ising action in equation (44).

So inserting $\tilde{C}^*_{\gamma\delta}$ into equation (46) we get

$$\begin{aligned}\Gamma(Q,\{R_k\}_k) = \mathcal{V}\left(0,\{R_k\}_k\right) \\ &+ P(P-1)\,\frac{N}{2}\,\frac{Q - \frac{1}{N}\sum_{i=1}^{N} R_{k_i}^2}{\frac{1}{N}\sum_{i=1}^{N} R_{k_i}^2 (1-R_{k_i})^2}\, Q \\ &- P(P-1)\,\frac{1}{2}\,\frac{Q - \frac{1}{N}\sum_{i=1}^{N} R_{k_i}^2}{\frac{1}{N}\sum_{i=1}^{N} R_{k_i}^2 (1-R_{k_i})^2} \sum_{i=1}^{N} R_{k_i}^2 \\ &- P(P-1)\left(\frac{1}{2}\,\frac{Q - \frac{1}{N}\sum_{i=1}^{N} R_{k_i}^2}{\frac{1}{N}\sum_{i=1}^{N} R_{k_i}^2 (1-R_{k_i})^2}\right)^2 \sum_{i=1}^{N} R_{k_i}^2 (1-R_{k_i})^2.\end{aligned}$$

Combining the terms

$$\begin{aligned}\Gamma(Q,\{R_k\}_k) = &\mathcal{V}(0,\{R_k\}_k)\\ &+ P(P-1)\,\frac{N}{2}\,\frac{Q-\frac{1}{N}\sum_{i=1}^{N}R_{k_i}^2}{\frac{1}{N}\sum_{i=1}^{N}R_{k_i}^2(1-R_{k_i})^2}\left(Q-\frac{1}{N}\sum_{i=1}^{N}R_{k_i}^2\right)\\ &- P(P-1)\,\frac{N}{4}\,\frac{\left(Q-\frac{1}{N}\sum_{i=1}^{N}R_{k_i}^2\right)^2}{\frac{1}{N}\sum_{i=1}^{N}R_{k_i}^2(1-R_{k_i})^2}\\ =&\mathcal{V}(0,\{R_k\}_k)\\ &+ P(P-1)\,\frac{N}{4}\,\frac{\left(Q-\frac{1}{N}\sum_{i=1}^{N}R_{k_i}^2\right)^2}{\frac{1}{N}\sum_{i=1}^{N}R_{k_i}^2(1-R_{k_i})^2},\end{aligned}$$

we observe that the probability distribution can be written as

$$p(Q,\{R_k\}_k) = p\left(Q|\,\{R_k\}_k\right)\,p\left(\{R_k\}_k\right) \propto \exp\left(-\Gamma(Q,\{R_k\}_k)\right) \tag{49}$$

with the probability for a firing rate distribution

$$p\left(\{R_k\}_k\right) \propto \exp\left(-\mathcal{V}\left(0,\{R_k\}_k\right)\right)$$

and the probability for a correlation given a firing rate distribution

$$\begin{aligned}p\left(Q|\,\{R_k\}_k\right) &\propto \mathcal{N}\left(Q\,|\,\frac{1}{N_{\text{exp}}}\sum_{k=1}^{N_{\text{exp}}}R_k^2,\ \frac{2}{N\,P(P-1)}\frac{1}{N_{\text{exp}}}\sum_{k=1}^{N_{\text{exp}}}R_k^2(1-R_k)^2\right)\\ &=: \mathcal{N}\left(Q\,|\,\mu_Q,\,\Sigma_Q\right)\end{aligned} \tag{50}$$

which has the form of a Gaussian. The mean and covariance can be calculated from the experiment and the variance decreases with $N$ and $P$. As expected, the most likely value for $Q$ is the overlap of two random binary vectors, which we already calculated above in equation (41). We assume now that the brain is more efficient in distributing the stimuli over the available neuron space. The measured overlap for two stimuli $P = 2$ from the experiment is smaller than the random overlap and therefore indicates some efficiency of the mouse brain in distributing stimuli. In the following, we formalize the notion of efficiency by the relative number of sets with overlaps smaller than $Q^{\leftrightarrow}$. The efficiency $E$ of the brain to realize one set of stimuli depends on how many possible sets of stimuli exist, see Figure 12. Intuitively, $Q^{\leftrightarrow} \le \mu_Q$ should be as small as possible to keep different stimuli apart, but this is increasingly difficult the larger their number $P$ grows, as the variance $\Sigma_Q$ decreases with $P$:

$$\begin{aligned}E^{-1} &\overset{!}{=} \int_0^{Q^{\leftrightarrow}} dQ\,p\left(Q|\,\{R_k\}_k\right)\\ &= \frac{1}{2}\left(1+\text{erf}\left(\frac{Q^{\leftrightarrow}-\mu_Q}{\sqrt{2\,\Sigma_Q}}\right)\right).\end{aligned}$$

The efficiency lies in $E \in [2,\infty)$. For $E = 2$, which presents a minimum efficiency, the measured overlap reaches the random overlap $Q^{\leftrightarrow} = \mu_Q$.

Solved for $Q^{\leftrightarrow}$ we obtain

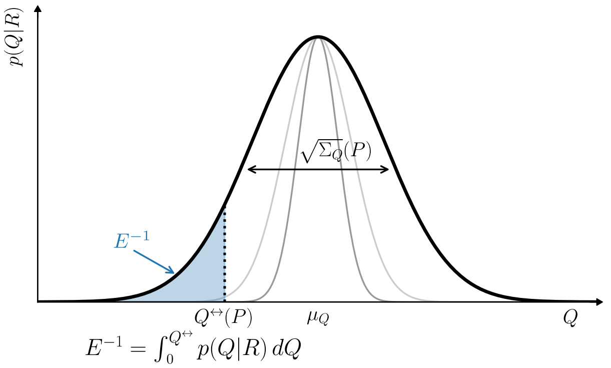

Figure 10: **Mutual overlaps of stimulus vectors.** Probability distribution of overlaps. The width of the distribution changes with the number of stimuli $P$, see equation (50). The efficiency $E$ is determined by demanding $Q^{\leftrightarrow}(2) \overset{!}{=} Q^{\leftrightarrow,\mathrm{exp}}$, which should be as large as possible to prevent overlap between stimulus classes. The larger the efficiency of the mouse brain, the fewer possible sets $\boldsymbol{x}_{1\leq\alpha\leq P} \in \{0,1\}^N$ (that fulfill conditions 1, 2 and 3) are needed to realize one set of them with the mutual overlap $Q$.

$$
\begin{aligned}
Q^{\leftrightarrow}(P) &= \frac{1}{N_{\mathrm{exp}}} \sum_{k=1}^{N_{\mathrm{exp}}} R_k^2 + \sqrt{2\,\Sigma_Q}\,\mathrm{erf}^{-1}\left(2E^{-1} - 1\right) \\
&= \frac{1}{N_{\mathrm{exp}}} \sum_{k=1}^{N_{\mathrm{exp}}} R_k^2 - 2\frac{\mathcal{E}}{\sqrt{N}} \frac{\sqrt{\frac{1}{N_{\mathrm{exp}}}\sum_{k=1}^{N_{\mathrm{exp}}} R_k^2(1-R_k)^2}}{\sqrt{P(P-1)}},
\end{aligned} \tag{51}
$$

where we rewrote the efficiency as $\mathcal{E} = -\mathrm{erf}^{-1}\left(2E^{-1} - 1\right) \in [0,\infty]$. We treat the so-determined $\mathcal{E}/\sqrt{N}$ as the efficiency of the mouse brain in relation to the available neuron space and infer it from the data by demanding $Q^{\leftrightarrow}(P = 2)$ to be the experimentally observed correlation. Note that this Gaussian approximation only holds in the vicinity of the mean $\mu_Q$, i.e. for an efficiency that is not too large.

This formula (51) provides us with the correction for a more efficient than random distribution of stimuli. In particular, we gain insights into how this correction scales with the number of stimuli $P$. If $P$ becomes very large, it becomes harder to efficiently distribute the stimuli such that $Q^{\leftrightarrow}$ will approach the average overlap $\mu_Q$ eventually.

#### Numerics

As seen in Figure 11 the distribution $p(Q|R)$ fits the simulation well and becomes more narrow for larger $N, P$. The coupling $\tilde{C}_{\alpha\beta}/N$ is small enough to justify the TAP-approximation in (47) and vanishes for the most likely overlap (see equation (48)).

In Figure 12 one can see how the inter-class correlation $Q^{\leftrightarrow}$ behaves according to equation (51). The absolute value of the coupling $\tilde{C}/N$ is sufficiently small, especially for larger $P$, justifying the perturbative treatment. For the smaller population mean and small $P$ the coupling is too large to justify the quadratic approximation for the effective action in equation (48) above. This, however, would only affect the mutual information for small $P$ for which we know that it has to be larger than the mutual information with larger population mean and smaller than the optimum $\log_2(P)$. In particular, the coupling will always become small enough for large $P$ such that the breakdown of the approximation does not affect the asymptotic information transfer of interest $\mathrm{MI}(\infty)$.

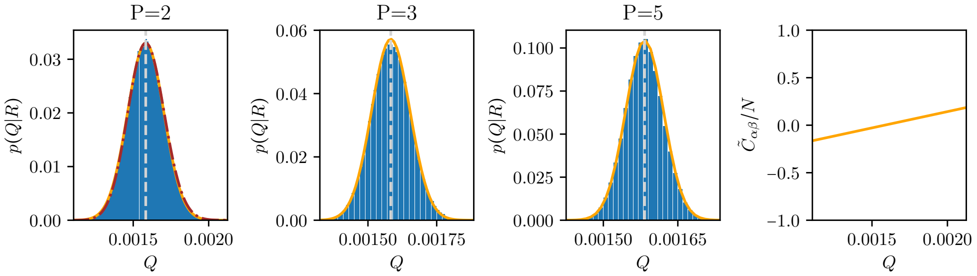

Figure 11: **Distribution for overlaps between stimuli.** Probability distribution for the overlap between stimuli (50) for different values of $P$. For simplicity we assume a homogeneous firing rate distribution $R_k = R$. We set $R$ close to the experimentally observed population mean $0.04$. Number of neurons $N = 10^5$. Blue: Numerical distribution by brute-force counting of overlaps for vectors of the same length $R$, orange: approximate distribution from mean-field theory. Light gray line indicates the overlap with highest probability $\frac{1}{N_{\mathrm{exp}}}\sum_{k=1}^{N_{\mathrm{exp}}} R_k^2 = R^2$. Rightmost panel shows the coupling $\tilde{C}_{\alpha\beta}$ in which we expanded to second order. Brown dashed-dotted line in the left panel for $P = 2$ shows exact probability distribution by solving equation (42) without approximations.

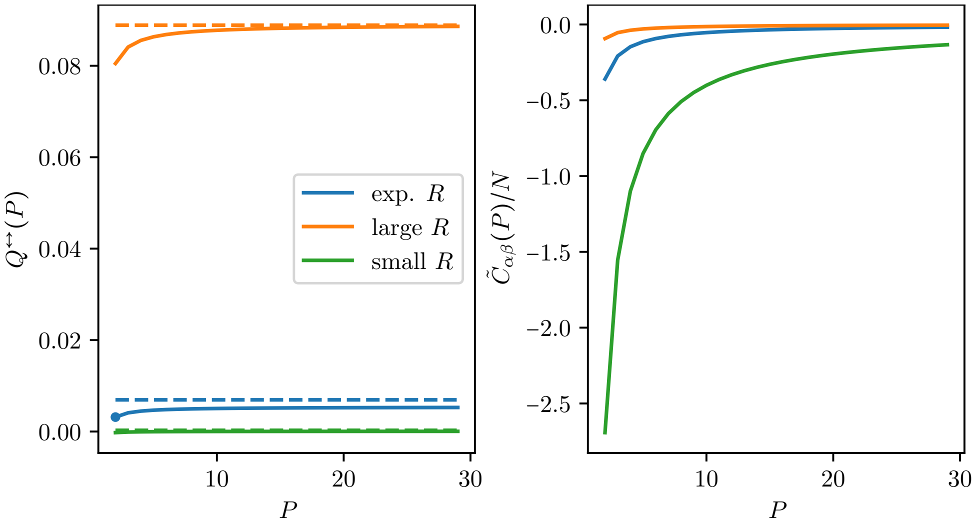

Figure 12: **Dependence of the inter-correlation $Q^{\leftrightarrow}$.** Different colors indicate different population means $R$. Dashed lines indicate intra-correlation $Q^{\circlearrowleft}$. For non-vanishing information transfer the inter-correlation has to be strictly smaller than the intra-correlation. Efficiency parameter is obtained by fixing the experimental inter-correlation for two stimuli (marked with a dot) $Q^{\leftrightarrow}(2) \overset{!}{=} Q^{\leftrightarrow,\mathrm{exp}} \Rightarrow E^{-1} = 0.13$. The inter-correlation reaches the value of intra-correlation at $P = 75$, marking the point of vanishing information transfer.

#### 5.5 Robustness of the vanishing regime

In the main text, we encountered the *vanishing regime*, where the mutual information asymptotically approaches zero as the number of stimuli increases. One assumption is that increasing the population mean preserves the shape of the firing rate distribution, scaling it by some factor to achieve the higher population mean. Here, we show that the vanishing regime is robust to modifications of the firing rate distribution *other than scaling*. These modifications arise from different regimes of the transfer function $\phi(h)$, which defines the relationship between input current $h$ and firing rate $R = \phi(h)$.

When examining the effect of changing the population mean $R$, a relationship between the overlaps $Q^{\circlearrowleft}$ and $Q^{\leftrightarrow}$ must be specified. While the between-classes overlap $Q^{\leftrightarrow}$ scales according to equation (51), for the within-class overlaps $Q^{\circlearrowleft}$, we assume that the selectivity of neurons across different classes $\rho_i$ remains constant as a function of the population mean $R$. This class selectivity for each neuron $i$ is captured by the Pearson correlation within classes, $C(\alpha) = C(\beta)$

$$
\begin{aligned}
\rho_i &= \frac{\mathrm{cov}(x_{\alpha i} x_{\beta i})_{(\alpha\beta)|\,C(\alpha)=C(\beta)}}{\sqrt{\mathrm{var}(x_{\alpha i})_\alpha \, \mathrm{var}(x_{\alpha i})_\beta}} \\
&= \frac{\mathrm{cov}(x_{\alpha i} x_{\beta i})_{(\alpha\beta)|\,C(\alpha)=C(\beta)}}{R_i\,(1 - R_i)}.
\end{aligned}
$$

The last equality follows from the binary nature of the states $\mathrm{var}(x_{\alpha i})_\alpha = \langle x_{\alpha i}\rangle_\alpha (1 - \langle x_{\alpha i}\rangle_\alpha)$ and $R_i = \langle x_{\alpha i}\rangle_\alpha$. $\mathrm{cov}(x_{\alpha i} x_{\beta i})_{(\alpha\beta)|\,C(\alpha)=C(\beta)}$ denotes the covariance over pairs of trials of the same stimulus class. We determine $\rho_i$ from the experiment. We can then express the overlap within classes using the Pearson correlation coefficient,

$$
\begin{aligned}
Q^{\circlearrowleft} &= \frac{1}{N}\sum_{i=1}^{N} \langle x_{\alpha i} x_{\beta i}\rangle_{(\alpha\beta)|\,C(\alpha)=C(\beta)} \\
&= \frac{1}{N}\sum_{i=1}^{N} \left[\mathrm{cov}(x_{\alpha i} x_{\beta i})_{(\alpha\beta)|\,C(\alpha)=C(\beta)} + \langle x_{\alpha i}\rangle_\alpha \langle x_{\beta i}\rangle_\beta\right] \\
&= \frac{1}{N}\sum_{i=1}^{N} \left[\rho_i\,(1 - R_i) + R_i\right] R_i \\
&= \frac{1}{N}\sum_{i=1}^{N} \left[R_i^2 + \rho_i R_i - \rho_i R_i^2\right].
\end{aligned}
$$

To calculate the population mean $R^*$, for which the mutual information vanishes in the limit $P \to \infty$, in which $Q^{\leftrightarrow} \to \frac{1}{N}\sum_{i=1}^{N} R_i^2$, we can equate within-class overlap to between-classes overlap,

$$
\begin{aligned}
& Q^{\circlearrowleft} = Q^{\leftrightarrow} \\
\Rightarrow \quad & \frac{1}{N}\sum_{i=1}^{N} \rho_i R_i = \frac{1}{N}\sum_{i=1}^{N} \rho_i R_i^2. \qquad (52)
\end{aligned}
$$

Notably, the resulting population mean marking the transition given is invariant under the rescaling $\rho_i \to \mathrm{const}\,\rho_i$, such that the lines for constant $\rho_i$, $\rho_i^{\mathrm{small}} = 0.2\,\rho_i$, and $\rho_i^{\mathrm{large}} = 2.5\,\rho_i$ (black and dashed lines) show the same point of transition $R^*$ in Figure 7a.

##### Exponential transfer function

If the population mean is low, the neural transfer function $R = \phi(h)$ has an exponential shape [17], leading to the scaling of the firing rates by a common factor when the inputs obtain an additive term $h \to h + \mathrm{const.} \Rightarrow \exp(h) \to \exp(\mathrm{const.})\exp(h)$.

When scaling the firing rate distribution with this scaling factor, which we call $l$ here ($l = R/R^{\mathrm{exp}}$), we find a population mean $R^*$ that marks the onset of the vanishing regime. Using equation (53), we obtain

$$R^* = R^{\mathrm{exp}} \frac{\frac{1}{N}\sum_{i=1}^{N} \rho_i R_i^{\mathrm{exp}}}{\frac{1}{N}\sum_{i=1}^{N} \rho_i \left(R_i^{\mathrm{exp}}\right)^2}.$$

If the degree of class-selectivity is similar for all neurons, $\rho_i \approx \rho$, $R^*$ takes the form

$$R^* = \frac{\left(\frac{1}{N}\sum_{i=1}^{N} R_i^{\mathrm{exp}}\right)^2}{\frac{1}{N}\sum_{i=1}^{N} \left(R_i^{\mathrm{exp}}\right)^2}. \tag{53}$$

It decreases the larger the random overlap $\frac{1}{N}\sum_{i=1}^{N}\left(R_i^{\mathrm{exp}}\right)^2$ (denominator) is compared to the squared population mean (numerator). Thus, the larger the heterogeneity in firing rates, the smaller the population mean has to be for the vanishing regime.

##### Linear transfer function

Neurons at sufficiently high firing rates typically show an affine linear dependence $R = \phi(h)$ [17, 18]. An additive term to their input then results in an additive term to every firing rate $h \to h + \mathrm{const.} \Rightarrow \phi(h) \to \phi(h) + \mathrm{const.}$. This transformation does not affect the variance $\langle\!\langle R_i^2 \rangle\!\rangle_i$. By rewriting equation (52) as $R(1-R) = \langle\!\langle R_i^2 \rangle\!\rangle_i$ we obtain a solution for the rate $R^*$ to transition into the vanishing regime that is larger than the $R^*$ in (53).

$$R^* = \frac{1}{2} + \sqrt{\frac{1}{4} - \langle\!\langle R_i^2 \rangle\!\rangle_i}.$$

In conclusion, for a realistic transfer function with an exponential shape at low rates and a linear part at higher rates, the transition into the vanishing regime lies between the two values of $R^*$ calculated for the exponential and linear case, respectively. Thus the transition to the vanishing regime persists.

##### 5.5.1 Mutual information limit $R \to 0$

If the population mean $R$ approaches the magnitude of the noise $\kappa$, the mutual information decreases:

$$\begin{aligned}
\delta\mu &= \left(1 + \frac{1}{N_{\mathrm{train}}} \frac{\kappa + R + Q^{\circlearrowleft}}{Q^{\circlearrowleft} - Q^{\leftrightarrow}}\right)^{-1} \\
&\leq \left(1 + \frac{1}{N_{\mathrm{train}}} \frac{\kappa}{R - Q^{\leftrightarrow}}\right)^{-1} \\
&\overset{R \to (Q^{\leftrightarrow})^+}{\longrightarrow} 0,
\end{aligned}$$

where we used that the intra-correlation is bounded by the population mean $Q^{\circlearrowleft} \leq R$.

Since $R \geq Q^{\leftrightarrow}$ we also have $\delta\mu \overset{R\to 0}{\longrightarrow} 0$. Together with $\Sigma \overset{R\to 0}{\longrightarrow} \mathrm{const.}$ we obtain $\Delta \overset{R\to 0}{\longrightarrow} 0$ and thus $\mathrm{MI}(P, \Delta) \overset{R\to 0}{\longrightarrow} 0$.

#### 5.6 Third regime for asymptotic information transfer

There actually is another regime for the asymptotic information transfer: Besides the vanishing regime, in which the mutual information converges to zero and the expanding regime, in which the mutual information monotonically grows as a function of the number of stimuli $P$, there is a third, small regime, in which the mutual information first grows with $P$, but then decreases and converges to a non-zero value Figure 13. Similar to the vanishing

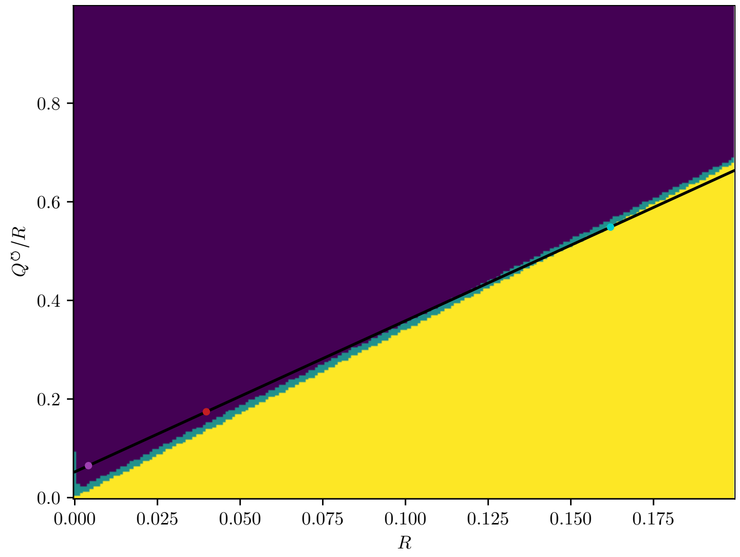

Figure 13: **Third regime for asymptotic information transfer.** Different regimes for asymptotic information transfer $\mathrm{MI}(\infty)$. Vanishing regime (yellow), expanding regime (indigo) and the third regime (green). Axis represent different population means $R$ and response variability within classes $Q^{\circlearrowleft}/R$. $Q^{\circlearrowleft}/R$ can be understood as the relative extent of the neural representation ellipsoid.

regime, for the third regime there exists a $P^*$ for optimal information transmission. This third regime connects the vanishing and expanding regime but is of negligible width.
